# GSE1 links the HDAC1/CoREST co-repressor complex to DNA damage

**DOI:** 10.1101/2023.03.13.532402

**Authors:** Terezia Vcelkova, Wolfgang Reiter, Martha Zylka, David M. Hollenstein, Stefan Schuckert, Markus Hartl, Christian Seiser

## Abstract

Post-translational modifications of histones are important regulators of the DNA damage response (DDR). By using affinity purification mass spectrometry (AP-MS) we discovered that genetic suppressor element 1 (GSE1) forms a complex with the HDAC1/CoREST deacetylase/demethylase co-repressor complex. In-depth phosphorylome analysis revealed that loss of GSE1 results in impaired DDR, ATR signalling and γH2AX formation upon DNA damage induction. Altered profiles of ATR target serine-glutamine motifs (SQ) on DDR-related hallmark proteins point to a defect in DNA damage sensing. In addition, GSE1 knock-out cells showed hampered DNA damage-induced phosphorylation on SQ motifs of regulators of histone post-translational modifications, suggesting altered histone modification. While loss of GSE1 does not affect the histone deacetylation activity of CoREST, GSE1 appears to be essential for binding of the deubiquitinase USP22 to CoREST and for the deubiquitination of H2B K120 in response to DNA damage. The combination of deacetylase, demethylase, and deubiquitinase activity makes the USP22-GSE1-CoREST subcomplex a multi enzymatic eraser that seems to play an important role during DDR. Since GSE1 has been previously associated with cancer progression and survival our findings are potentially of high medical relevance.

## Introduction

DNA damage represents a challenge to the genomic integrity on a daily basis. To address this constant threat, the eukaryotic cell has developed a system integrating multiple pathways that act synergistically to promote cell survival in response to DNA damage. Once DNA damage is sensed, a cell cycle arrest accompanied by DNA repair signalling is initiated. Impaired repair mechanisms have a severe impact on fitness of the organism, leading to the onset of diseases such as cancer (reviewed in (1)).

Post-translational modifications of histones such as phosphorylation, methylation, monoubiquitination and acetylation have been shown to be important regulators of the DNA damage response (DDR) (reviewed in (2)). For instance, protein kinases ATM (ataxia telangiectasia mutated), DNA-PK (DNA-dependent protein kinase) or ATR (ATM and Rad3 related) phosphorylate histone variant H2AX at serine 139 (γH2AX), which is one of the earliest markers of DNA damage and crucial for recruitment of downstream effector proteins such as MDC1, 53BP1, RAD50, RAD51 or BRCA1 to sites of DNA lesions (3-5). In addition, histone H2A is ubiquitinated at the site of these lesions by the E3 ubiquitin ligases RNF8/RNF168, and histone H2B is monoubiquitinated by the RNF20/RNF40 ubiquitin-protein ligase complex. Ubiquitination of the two histones then enables the assembly of the DNA repair machinery (6-10). However, formation of γH2AX may require dynamic regulation of H2B monoubiquitination, as loss of the H2B-specific deubiquitinase USP22 led to reduced levels of γH2AX (11). The lysine-specific demethylase LSD1/KDM1A can be recruited to sites of DNA damage by physical interaction with RNF168, catalysing demethylation of dimethylated lysine 4 of histone H3 (H3K4me2) (12). Alongside, H4K16 acetylation becomes increased upon ionising radiation (13) and deacetylation of H3K9 and H2K56 upon genotoxic stress has been reported (14). Experiments in diverse cancer cells have shown that treatment with HDAC inhibitors can impair DNA damage repair. Based on these results it has been proposed that specific histone deacetylation is necessary for the repair and for the restoration of local chromatin environment (15-19).

There is increasing evidence that all these PTMs must be orchestrated sequentially and many of them are interdependent for proper signal propagation (20-25). Moreover, to resolve DNA damage signalling, these modifications must also be removed consecutively, which requires coordinated cooperation of respective eraser enzymes. Nevertheless, it has not been understood how these enzymes uphold their activities and whether they function as a united entity within one or multiple specific complexes. Moreover, their capacity lies beyond histones since PTMs on non-histone proteins are being vastly identified and functionally annotated to the DDR in recent years (26-29).

In this study, we propose a potential mechanism for regulating the DDR by identifying a multi-enzymatic eraser sub-complex of HDAC1/CoREST (co-repressor of Repressor Element 1 Silencing Transcription Factor). This sub-complex combines the activities of the deacetylase HDAC1 and the demethylase activity LSD1 with the deubiquitinase activity of USP22. Using affinity-purification mass spectrometry (AP-MS) we identified GSE1 (genetic suppressor element 1) as a prominent interaction partner of HDAC1, and more specifically, as an interactor of the COREST complex. While loss of GSE1 did not affect the deacetylase activity of the CoREST complex, it resulted in reduced γH2AX formation and an impaired signalling response upon DNA damage. Further, by in-depth quantitative mass spectrometry, we determine common and individual effects caused by loss of GSE1 or HDAC1 activity. We show that GSE1 acts at an initial step of the DDR. Ultimately, we demonstrate that GSE1 is required for the recruitment of the deubiquitinase USP22 to the CoREST complex, potentially serving as a scaffold, and that loss of GSE1 inhibits deubiquitination of histone H2B at lysine 120, which has been previously shown to facilitate γH2AX formation (11,30).

Through biochemical and phenotypic characterisation of GSE1, we unravel its role in response to DNA damage as a part of multi-enzymatic eraser complex together with HDAC1/2, USP22 and LSD1, which maintains proper chromatin environment and downstream DNA repair signalling. Hence, we emphasise the importance of understanding the cooperation of not only writers but also eraser enzymes within the kinetic window of DNA damage and the chromatin challenges that have to be resolved during DNA repair.

## Materials and methods

### Cell culture

HAP1 cells were cultured in Iscove’s Modified Dulbecco Medium (Sigma Aldrich, I3390-500ML), 293T HEK and MEF in Dulbecco’s modified Eagle’s medium (Sigma Aldrich, D6046-500ML). The culture media was supplemented with 10% foetal bovine serum (Sigma Aldrich, F7524) and 1% penicillin/streptomycin (Sigma Aldrich, P4333-100ML). The cells were kept in an incubator at 37°C and 5% CO_2_. Treatments of the cells were performed with Mitomycin C, Etoposide or Hydroxyurea that were purchased from Sigma Aldrich. Mitomycin C (10107409001) was used at final concentration of 500 nM, Etoposide (E1383-25MG) at 0.1 μM and hydroxyurea (400046-5GM) at 200 μM if not stated otherwise. The cells were treated either for 6 or 24 hours with the chemicals in full culturing media. To harvest, cells were washed once with PBS on ice, scraped and collected by centrifugation (300 x g / 5 min, 4°C).

### Cell transfection

For transfection of HAP1 cells polyethyleneimine PEI/NaCl solution (100 μl of 150 mM NaCl, 7 μl of PEI) and DNA/NaCl solution (100 μl of 150 mM NaCl, 3 μg of plasmid DNA) were mixed, incubated at room temperature for 30 min, and added dropwise onto a plate of 0.5 x 10^6^ HAP1 cells. Human Gse1 plasmid was designed and purchased from VectorBuilder (pRP-hGse1).

### CRISPR-Cas9 gene editing

GSE1 KO HAP1 cell line (HZGHC007762c010) was obtained from Horizon Genomics (now Horizon Discovery). To tag endogenous human *Gse1* in HAP1 cells with V5-tag, Precise Integration into target chromosome (Crispr PITCh) was used as described before (31). The sequence of guide RNA was designed using an online web tool of IDTDNA company (https://eu.idtdna.com/site/order/designtool/index/CRISPR_SEQUENCE). The RNA duplex complex was assembled by adding 320 pmol guide RNA and 320 pmol tracrRNA (IDTDNA) in IDTE buffer and annealed in the following reaction: 94°C for 5 min, twelve times 93°C for 20 s with -1°C/cycle, 80°C for 4 min, three times 79°C for 20 s with -1°C/cycle, 75°C for 4 min, three times 74°C for 20 s with -1°C/cycle and finally at 70°C for 4 min. Next, the RNP complex was prepared using RNA duplex with 5 μg Cas9 protein in 2 μl cleavage buffer (20 mM Hepes, pH=7.5, 0.15 M KCl, 0.5 mM DTT, 0.1 EDTA) and incubated for 10 min at room temperature. After incubation, 2 x 10^6^ of HAP1 cells were electroporated using The Neon Transfection System^TM^ (MPK5000) set to 3 pulses, 10 ms width and 1575 V. The obtained single clones were cultured and screened for V5-tag integration by PCR (fw: 5’-GGT TTA CCC TTC CAT CCA GCA-3’; rv: 5’-AGG AGA GGG TTA GGG ATA GGC-3’).Guide RNA sequence: GCT ACC TGA AGG GAT ATC CC. Homology arm sequence: 5’-CTG GAC CAC TTA CGA AAG TGC CTT GCC TTG CCT GCA ATG CAC TGG CCT AGG GGC TAC CTG AAG GGA TAT CCC AGG GGT AAG CCT ATC CCT AAC CCT CTC CTC GGT CTC GAT TCT ACG TGA CGG TTT CCC TTG CAC TAG GCC GAA CCT ATA GTA TAG AAA TAT TAT CTA TTT TAT TAC CTT GAA TAT TTA ATA TTT-3’

### Cell cycle analysis

To monitor changes in cell cycle, fluorescent ubiquitination cell cycle indicator system (FUCCI) was used as described before (32). The 293T HEK cells were transfected with Lipofectamine^TM^ 3000 (Thermo Fisher, L3000075) to produce lentiviral particles containing pBOB-EF1-FastFUCCI-PURO according to the manufacturer’s protocol. 48 hours post transfection, media containing virus was collected, filtered, and added onto HAP1 cells. After 48 hours, the positive cells were selected by addition of 1 μg/ml puromycin for two days. FUCCI positive cells were cultured and washed once with phosphate-buffered saline (PBS, 300 x g / 5 min) and fixed with 4% methanol-free formaldehyde (Thermo Fisher Scientific, 28906) for 20 min at room temperature. Afterwards, the samples were washed twice with PBS and resuspended in the final volume of 200 μl PBS and taken for measurement. The samples were measured with Cytoflex Flow Cytometer (Beckman Coulter Life Sciences) and analysed with CytExpert software (Beckman Coulter Life Sciences) or FLowJo (TreeStar Inc.).

### Western blot analysis

To generate whole-cell lysates, cells were washed in 1 ml PBS on ice and centrifuged at 300 x g/ 5 min. The pellets were lysed in Hunt buffer (20 mM Tris/HCl ph=8.0, 100 mM NaCl, 1 mM EDTA, 0.5% NP-40) supplemented with Complete Protease inhibitor cocktail (Roche), Complete Phosphatase inhibitor cocktail (Roche, 11836145001), 10 mM sodium fluoride, 10 mM β-glycerophosphate, 0.1 mM sodium molybdate and 0.1 mM PMSF. The samples were frozen and thawed three times in liquid nitrogen and centrifuged (21 000 x g / 30 min, 4°C). The supernatants were transferred, and protein lysates concentration measured by Bradford assay. Detection of proteins was performed by SDS-PAGE and western blots using nitrocellulose membranes and chemiluminescence. The blots were developed using ECL Western Blot detection reagents (Ge Healthcare, RPN2106) and visualised via chemiluminescence using Fusion FX7-Spectra machine.

Antibodies: KIAA0182 (Proteintech, 24947-1-AP, 1:1000), HDAC1 SAT 208 (Seiser lab, polyclonal rabbit, 1: 10 000), HDAC2 clone 3F3 (Seiser/Ogris lab, monoclonal mouse, 1:1000), CoREST (Millipore, 07-455, 1:1000), β-actin (Abcam, ab8226, 1:1000), cleaved caspase-3 (Cell Signalling, 9661, 1:1000), γH2AX ( Millipore, JBW301, 1:1000), vinculin (Cell Signalling, 13901, 1:1000), LSD1 (Cell Signalling, 2139, 1:1000), USP22 (Novus Biologicals, NBP1-49644, 1:1000)

### Histone deacetylase activity measurements

The deacetylase assay was conducted on beads carrying precipitated proteins in 20 μl volume. 4 μl of [^3^H]-acetate-labelled chicken reticulocyte core histones (1.5-2 mg/ml) were added. Following 3 hours incubation on thermomixer (300 rpm, 30°C), 35 μl of Histone stop solution (1 M HCl, 0.4 M sodium acetate) and 800 μl of ethyl acetate were added and vortexed for 15 sec. The samples were centrifuged (11 700 x g / 4 min, RT) and 600 μl of organic upper phase was mixed with 3 ml of Scintillation Solution (5 g/l PPO, 0.5 g/l POPOP in toluene). Released acetate was measured in counts per minute (CPM) with a Liquid Scintillation Analyzer (Packard).

### Histone isolation

To isolate histones, all the buffers were pre-cooled and kept on ice throughout the whole procedure. Cells were washed once with PBS on ice and centrifuged (300 x g / 5 min). The pellets were lysed in histone lysis buffer (100 mM Tris-HCl, 8.6% sucrose, 10 mM MgCl_2_, 50 mM NaHSO_4_, 1% Triton X-100, adjusted to pH=6.5) supplemented with Complete Protease inhibitor cocktail (Roche), Complete Phosphatase inhibitor cocktail (Roche, 11836145001), 10 mM sodium fluoride, 10 mM β-glycerophosphate, 0.1 mM sodium molybdate and 0.1 mM PMSF and centrifuged (450 x g / 5 min, 4°C). The nuclear pellet was washed three times with 1 ml of histone lysis buffer, once with histone wash buffer (100 mM Tris-HCl, 13 mM Na_3_EDTA, adjusted to pH=7.4) and centrifuged (450 x g / 5 min, 4°C). After each washing step the cells were briefly vortexed. Next, the pellets were resuspended in 100 μl H_2_O and concentrated H_2_SO_4_ was added to final concentration of 0.4 N. After 1 hr incubation on ice, the cells were centrifuged (18 400 x g / 1 hr, 4°C). The supernatants were transferred into a new tube with ten times volume of acetone, vortexed for 1 min and precipitated o/n at -20°C. The precipitates were collected by centrifugation (18 400 x g / 15 min, 4°C), air-dried and resuspended in H_2_O. 2 μg of histones were used for SDS-PAGE and western blot.

### Indirect immunofluorescence analysis

The cells were grown on coverslips. The cells were pre-extracted with 0.1% Tween-20 in PBS (PBST) for 1 min and then fixed with 4% methanol-free formaldehyde (Thermo Fisher Scientific, 28906) for 20 min at room temperature. The samples were washed two times with PBS and permeabilized with 0.5% Triton-X in PBS for 10 min at room temperature. After three times washing with PBS, the samples were blocked with 5% bovine serum in PBST for 1 hour at room temperature. After blocking, the primary antibody was added in blocking solution and incubated overnight at 4°C. The samples were washed three times with 3% bovine serum in PBS for 5 min and stained with secondary antibody in blocking solution for 1 hour at room temperature in the dark. Next, samples were washed three times with 3% bovine serum in PBS for 5 min, incubated with DAPI in PBS for 10 min at room temperature and washed again two times with PBS. Afterwards, the slides were mounted using DAKO mounting medium and dried. Antibodies: DAPI (Sigma Aldrich, D9542-10MG,1:10000), γH2Ax (Millipore, JBW301,1:1000), H3Ser10ph (Santa Cruz, 8656, 1: 2000), Alexa Fluor 546 (Thermo Scientific, A11003,1:500), Alexa Fluor 488 (Thermo Scientific, A32731,1:500)

The staining was monitored by Olympus Confocal Microscope and the signals were quantified using ImageJ software. Either number of foci was counted and compared, or signal strength (integrated density) was measured. For quantification of G2 and M phase positive cells stained with H3Ser10ph antibody. As described in De La Barre *et al*, 2000 (33) the cells were counted as G2 positive when pericentromeric chromatin was visible as foci and as M positive when signal was fully saturated across the nucleus.

### Co-immunoprecipitation assays

Whole-cell lysates were prepared as described above (Protein lysates, western blot). 40 μl Protein A or Protein G Dynabeads (Thermo Fisher Scientific, 1002D / 1004D) per sample were washed three times with Hunt buffer, resuspended in Hunt buffer containing 10% bovine serum albumin and incubated for 1 hour rotating at 4°C for blocking. At the same time, the whole-cell extract (0.5-1 mg protein) was incubated with an antibody mix suitable for immunopurification of the protein of interest for 1 hour rotating at 4°C. After blocking, the beads were added to the cell lysate-antibody mix and incubated overnight at 4°C rotating. Next day, the beads were washed twice with TBS for 5 min and once for 15 min at 4°C. Subsequently, the purified proteins were eluted from the beads in 1x SDS loading dye by heat-incubation at 95°C for 5 min. The eluate was used for SDS-PAGE and western blotting.

Antibodies: 4 μg COREST (Millipore, 07-455), 4 μg USP22 (Santa Cruz, 390585), 2 μg GSE1 (515046, Santa Cruz), 30 μl of monoclonal HDAC1 (clone 10E2) and HDAC2 (clone 3F3) generated in collaboration with Egon Ogris (Medical University of Vienna)

### Interactomics

#### Cell extract and mass spectrometry sample preparation

Whole cell extracts were prepared as follows: cell pellets were resuspended in Hunt buffer (20 mM Tris/HCl ph=8.0, 100 mM NaCl, 1 mM EDTA, 0.5% NP-40), sonicated twice for 3 sec (Bandelin Sonoplus, power of 45-50%), and centrifuged at 21 000 x g for 15 min at 4°C. Protein concentration of supernatants was determined via Bradford. 1 mg of protein was used for each sample.

80 μl of V5-beads (MBL International, M167-10) per sample were used for isolation of Gse1-V5, 120 µl of crosslinked M2-Flag beads (Sigma Aldrich, M8823-5ML) were used for purification of HDAC1-Flag. Briefly, for antibody crosslinking the bead slurry was equilibrated with sodium borate buffer (200 mM sodium borate pH 9.0), resuspended in 20 mM dimethyl pimelimidate in sodium borate buffer, incubated rotating for 1 hour at room temperature, washed and incubated with 200 mM ethanolamine pH 8.0 for 2 hours, washed with 100 mM glycine and subsequently stored in 1 x TBS. Beads were washed three times with Hunt buffer, resuspended in whole cell extracts and incubated overnight rotating at 4°C. Next day, the beads were washed twice with TBS for 5 min and once for 15 min rotating at 4°C. Beads were transferred to a 0.2 ml PCR tube, re-buffered to and washed once with ammonium bicarbonate (ABC) (Sigma Aldrich, 09830-500G) buffer, pelleted by gentle centrifugation, and resuspended in 50 µl 2 M urea (VWR, 0568-500G), 50 mM ABC.

Predigestion of bound proteins was performed by addition of 200 ng / µl of a 1:1 LysC (FUJIFILM Wako Pure Chemical Corporation, 125-02543) trypsin (Trypsin Gold, Mass Spec Grade, Promega, V5280) mix, followed by an incubation at RT for 90 minutes in the dark. The eluate containing predigested proteins and peptides was separated from the beads, and beads were subsequently rinsed with 50 µl ABC. The eluates were united.

Predigested proteins and peptides were reduced with 10 mM dithiothreitol (DTT) (Roche, 10197777001) for 30 min at RT, and subsequently carbamidomethylated using 20 mM iodoacetamide (IAA) (Sigma Aldrich, I6125-5G) followed by incubation for 30 min at RT in the dark. The alkylation reaction was quenched by adding 5 mM DTT and incubation for 10 min. Samples were digested overnight at 37°C using 150 ng / µl of the 1:1 LysC and trypsin mix, acidified using 10% trifluoroacetic acid (TFA) (Thermo Scientific, 28903) to a final concentration of 0.5% and desalted on C18 (Empore, 2215-C18) StageTips as described in (34).

#### Mass spectrometry measurements

MS measurements were performed on a Q Exactive HF Orbitrap (with HCD, higher-energy collisional dissociation mode) mass spectrometer (Thermo Fisher), operated in data-dependent acquisition (DDA) mode with dynamic exclusion enabled. For peptide separation on the HPLC, the concentration of organic solvent (acetonitrile, VWR, 83639.320) was increased from 1.6% to 32% in 0.1% formic acid at a flow rate of 230 nl / min, using a 2 hours gradient time. Peptides were ionised with a spray voltage of 1,9 kV. MS1 survey scans were recorded with the following parameters: resolution 120,000, scan range 375-1,500 m/z, automatic gain control (AGC) target = 3e6, and maximum injection time (IT) = 60 ms. MS2 analysis was performed with the following parameters: resolution 30,000, AGC target = 1e5, maximum IT = 250 ms, isolation window 1.6 m/z. The mass spectrometer was configured to pick the eight most intense precursor ions for data-dependent MS2 scans, applying HCD for fragmentation with a normalised collision energy (NCE) of 28. Dynamic exclusion time was set to 30 s.

#### Mass spectrometry data analysis with MaxQuant

MS raw files were searched using MaxQuant (version 2.1.4.0) against the Homo sapiens (human) (HAP1 samples) or the Mus Musculus (mouse) (MEF samples) Uniprot database (release 2022.02) and a database of common laboratory contaminants. Enzyme specificity was set to “Trypsin/P”. Carbamidomethylation of cysteine was searched as a fixed modification. Protein N-term acetylation and Oxidation (M) were set as variable modifications. The data were filtered at 1% FDR (PSM and protein), reverse hits were removed. MaxLFQ and match-between-runs were activated. MaxQuant output was further processed in R using Cassiopeia (35). After removing contaminants and proteins with less than 2 razor & unique peptides, missing LFQ protein intensities were imputed if at least two valid values were present in one experimental group. Imputation was performed in log2 space by drawing random values from a normal distribution (µ = mean sample LFQ intensity - 1.8 standard deviations of the sample LFQ intensities, σ = 0.3 * standard deviation of the sample LFQ intensities). LIMMA was used to determine differentially enriched proteins at 1% FDR (Benjamini-Hochberg).

Inspection of the HDAC1-FLAG IP data obtained from HAP1 cell extracts (Figure 1A) revealed that MaxLFQ did not properly normalise the data due to the high number of enriched proteins, violating the assumption of MaxLFQ that the majority of proteins is not regulated. For this reason, in a first analysis Cassiopeia was used to determine a subset of background proteins (determined by k-means clustering in Cassiopeia) for the respective MQ output files. Then the Cassiopeia script was run again on the original proteinsGroups.txt file but specifying the background protein cluster to be used for normalisation in the Cassiopeia script. In contrast to the original Cassiopeia script LFQ values were used for renormalisation instead of raw intensities.

**Figure 1.**
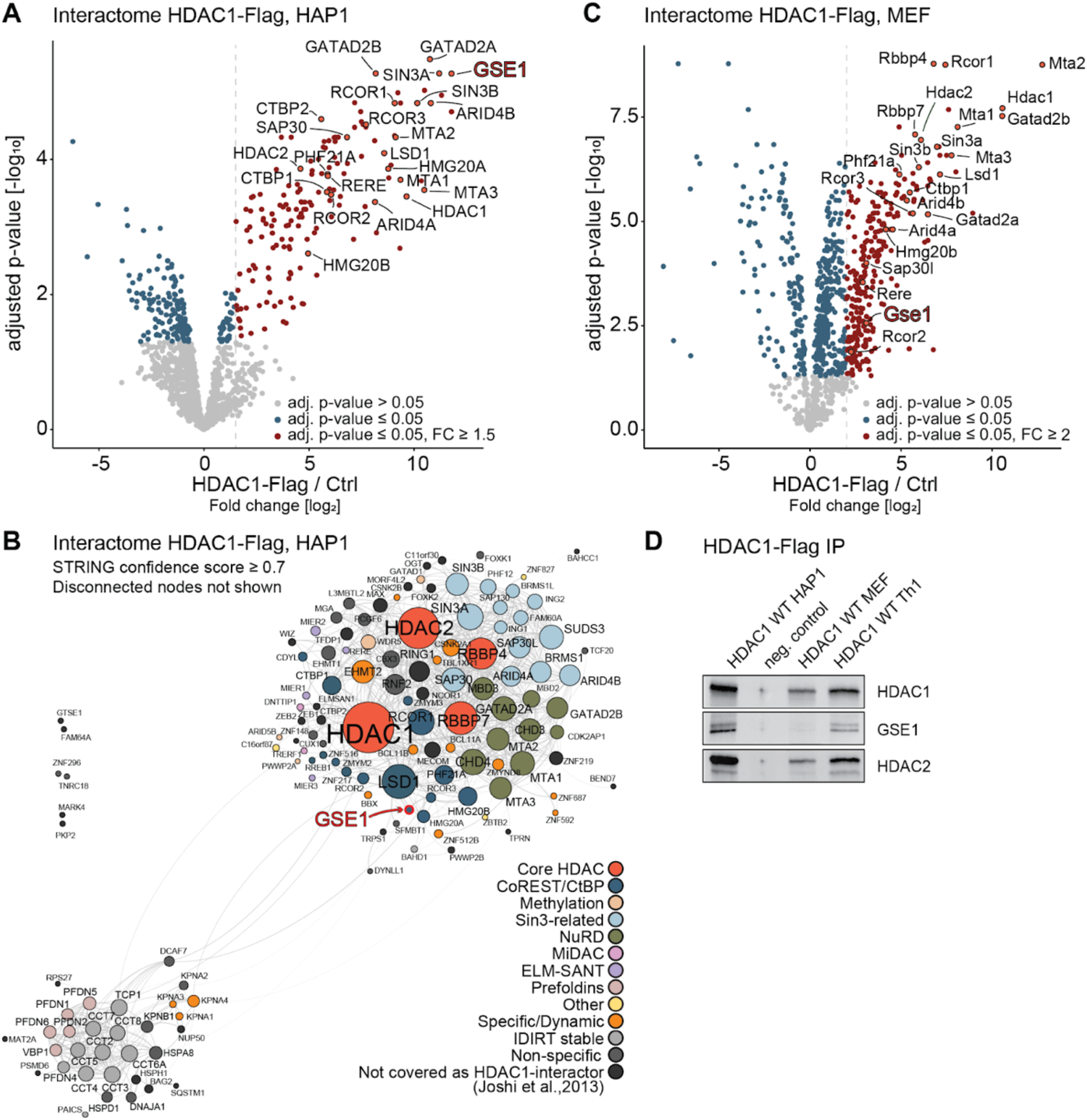
GSE1 as a prominent interactor of HDAC1 across species. **(A)** Volcano plot displaying the interactome of FLAG-tagged HDAC1 in HAP1 cells determined by AP-MS. Significant interactors (≥ 1.5-fold change [FC] over HDAC1 KO [Ctrl], adjusted p-value ≤ 0.05) in HDAC1-Flag cells are shown with red dots. Dashed line indicates fold change cut-off. Gene names of known subunits of the HDAC1 corepressor complexes are highlighted, such as CoREST, SIN3-related or NuRD. Among the binding partners GSE1 is highlighted with bold red text. **(B)** STRING DB-based protein interaction network of the HDAC1 interactome in HAP1 cells. High-confidence interactions (STRING confidence score ≥ 0.7) were selected. Disconnected nodes are not shown. The interactors were classified into functional protein groups in accordance with Joshi *et al.* (41). Strength of the associations is represented by the thickness of the edges. **(C)** Volcano plot displaying the interactome of FLAG-tagged HDAC1 in mouse embryonic fibroblasts (MEFs) cells determined by AP-MS. Significant interactors (≥ 1.5-fold change [FC] over wildtype cells [Ctrl], adjusted p-value ≤ 0.05) in HDAC1-Flag are shown with red dots. Dashed line indicates fold change cut-off. Gene names of known subunits of the HDAC1 corepressor complexes are highlighted, such as CoREST, SIN3-related or NuRD. Among the binding partners GSE1 is highlighted with bold red text. **(D)** Co-IP/western blot analysis illustrating HDAC1-GSE1 interaction in HAP1, MEFs and murine Th1 T-cells. HDAC1 KO HAP1 cells were used as negative control. Antibodies used for detection are depicted on the right.

### Proteome, acetylome, and phosphorylome analysis

#### Preparation of cells

Three biological replicates of each experimental condition were prepared. To induce DDR cells were treated with 0.1 µM etoposide (Sigma Aldrich, E1383-25MG) for 6 h. Cells were washed twice in 1x PBS, supplemented with 10 mM sodium butyrate (Sigma Aldrich, 303410) and 3 mM nicotinamide (NAM) (Sigma Aldrich, 303410), and pelleted prior to lysis.

#### Cell extract and mass spectrometry sample preparation

MS sample preparation was performed as previously described (36). Briefly, 2 x 10^7^ cells per replicate were resuspended in lysis buffer (8 M urea (VWR, 0568-500G), 50 mM Tris-HCl pH8.0, 150 mM NaCl, 1 mM PMSF, 5 mM sodium butyrate (Sigma-Aldrich,303410), 10 ng/ml NAM (Sigma Aldrich, 72340), 10 ng/ml trichostatin A (TSA) (Sigma Aldrich, T8552-1MG), 1x cOmplete protease inhibitor cocktail (+EDTA) (Roche, 11697498001), 250 U/replicate benzonase (Merck, 1.01695.0001)), lysed using a Bioruptor sonication device (Diagenode) (settings: 30 sec sonication (power level H, 30 sec cooling, 5 cycles), centrifuged at 15,000 x g for 10 min at 4°C and precipitated with 4x volumes of cold (-20°C) 100% acetone (overnight, 4°C). Proteins were washed with cold (-20°C) 80% acetone, air dried for 5 min, dissolved in 8 M urea, 50 mM ABC (Sigma Aldrich, 09830-500G), reduced with 10 mM DTT (Roche, 10197777001) for 45 min at RT, and subsequently carbamidomethylated with 20 mM IAA (Sigma Aldrich, I6125-5G) for 30 min at RT in the dark. The alkylation reaction was quenched by adding 5 mM DTT for 10 min. The urea concentration was reduced to 4 M using 50 mM ABC. Samples were pre-digested for 90 min at 37°C with Lys-C (FUJIFILM Wako Pure Chemical Corporation, 125-02543) at an enzyme-to-substrate ratio of 1:100 and subsequently digested overnight at 37°C with sequencing grade trypsin (Trypsin Gold, Mass Spec Grade, Promega, V5280) at an enzyme-to-substrate ratio of 1:50. Digests were stopped by acidification with TFA (Thermo Scientific, 28903) (0.5% final concentration) and desalted on a 50 mg tC18 Sep-Pak cartridge (Waters, WAT054960). Peptide concentrations were determined and adjusted according to UV chromatogram peaks obtained with an UltiMate 3000 Dual LC nano-HPLC System (Dionex, Thermo Fisher Scientific), equipped with a monolithic C18 column (Phenomenex). Desalted samples were dried in a SpeedVac concentrator (Eppendorf) and subsequently lyophilized overnight.

#### Isobaric labelling using TMT 10plex

500 μg of trypsin-digested and desalted peptides were used for each sample for isobaric labeling. Lyophilized peptides were dissolved in 20 μl 100 mM triethylammonium bicarbonate (TEAB) (Supelco, 18597-100ML). 60 μg of peptides in TEAB from each sample and replicate were additionally mixed (final volume 60 μl) as reference sample (REF). 500 μg of each TMT 10plex reagent (Thermo Scientific, 90110) were dissolved in 30 μl of acetonitrile (ACN) (VWR, 83639.320) and added to the peptide/TEAB mixes. The REF sample was mixed with 1.5 mg (90 μl) of TMT 10plex reagent 131N and subsequently split in 3 aliquots. Labels used: 126C: wildtype (wt) replicate (rep) 1; 127C: wt rep 2; 128C: wt rep 3; 129C: HDAC1 CI rep 1; 130C: HDAC1 CI rep 2; 127N: HDAC1 CI rep 3; 128N: GSE1-KO rep 1; 129N: GSE1-KO rep 2; 130N: GSE1-KO rep 3; 131N: REF. Untreated and etoposide treated samples were labelled using 2 separate TMT 10plex reagent kits which were subsequently processed individually. Samples were labelled for 60 min at RT. 0.1 μl of each sample were pooled, mixed with 10 μl 0.1% TFA and analysed by MS to check labelling efficiency. Reactions were quenched using hydroxylamine (Sigma Aldrich, 438227-50ML) (0.4% final concentration), pooled, desalted using Sep-Pak tC-18 (200 mg) cartridges (Waters, WAT054925), dried in a SpeedVac vacuum centrifuge (Eppendorf), and subsequently lyophilized overnight. Neutral pH fractionation was performed as previously described (36), using a 60 min gradient of 4.5 to 45% ACN (VWR, 83639.320) in 10 mM ammonium formate (1ml formic acid (26N) (Merck, 1.11670.1000), 3ml ammonia (13N) (1.05432.1000) in 300ml H_2_O, pH = 7 - 8, dilute 1:10) on an UltiMate 3000 Dual LC nano-HPLC System (Dionex, Thermo Fisher Scientific) equipped with a XBridge Peptide BEH C18 (130 Å, 3.5 μm, 4.6 mm x 250 mm) column (Waters) (flow rate of 1.0 ml / min). Fractions were collected and subsequently pooled in a non-contiguous manner into 8 pools, dried for 30 min using a SpeedVac concentrator (Eppendorf) and subsequently lyophilized overnight.

#### Acetylated peptide enrichment

Acetylated peptides were enriched, washed, eluted and desalted as described in (36), using the PTMScan Acetyl-Lysine Motif Kit (Cell Signaling Technology, 11663V1). The flow-through of the Acetyl-Lysine immunoprecipitation was kept for phosphorylated peptide enrichment. Eluted peptides were dried to completeness in a SpeedVac concentrator (Eppendorf) and subsequently resuspended in 0.1% TFA.

#### Phosphorylated peptide enrichment using TiO_2_

An aliquot (peptide: TiO_2_ resin = 1:6) of TiO_2_ (Titansphere TiO, GL Sciences, 5020-75000) was washed twice with 50% methanol (Fisher chemical, A456-212) and twice with glycolic acid solution (1 M glycolic acid (Sigma Aldrich, 124737-25G), 70% ACN (VWR, 83639.320), 3% TFA (Thermo Scientific, 28903)). Lyophilized peptides were dissolved in glycolic acid solution, mixed with the TiO_2_ resin, and incubated rotating at RT for 30 min. Samples were loaded onto Mobicol columns (MoBiTec, M1003, M2110) and shortly centrifuged in a table centrifuge to remove unphosphorylated peptides. Phosphorylated peptides will bind to the TiO_2_ resin. The resin was washed twice with glycolic acid solution, twice with 200 µl 70%, ACN 3%, TFA and twice with 1% ACN, 0,1% TFA. Phosphorylated peptides were eluted twice using 150 µl 300 mM ammonium hydroxide (VWR, 1.05432.1000), eluates were united and immediately acidified with conc. TFA to a pH of 2,5. Samples were desalted using a standard C18 (Empore, 2215-C18) StageTip protocol (34), dried using a SpeedVac (Eppendorf) and dissolved in 20 ml 2% ACN, 0,1% TFA.

#### Mass spectrometry measurements

DDA-SPS-MS3 method with online real-time database search (RTS) (for proteome analysis) was performed on an UltiMate 3000 Dual LC nano-HPLC System (Dionex, Thermo Fisher Scientific), containing both a trapping column for peptide concentration (PepMap C18, 5x 0.3 mm, 5 μm particle size) and an analytical column (PepMap C18, 500 x 0.075 μm, 2 μm particle size, Thermo Fisher Scientific), coupled to a Orbitrap Eclipse mass spectrometer (Thermo Fisher) via a FAIMS Pro Duo ion source (Thermo Fisher). The instruments were operated in data-dependent acquisition (DDA) mode with dynamic exclusion enabled. For peptide separation on the HPLC, the concentration of organic solvent (acetonitrile, VWR, 83639.320) was increased from 1.6% to 32% in 0.1% formic acid at a flow rate of 230 nl / min, using a 2 hours gradient time for proteome analysis. Peptides were ionised with a spray voltage of 2,4 kV. The instrument method included Orbitrap MS1 scans (resolution of 120,000; mass range 375−1,500 m/z; automatic gain control (AGC) target 4e5, max injection time of 50 ms, FAIMS CV -40V, -55V and -70V, dynamic exclusion 45 sec), and MS2 scans (CID collision energy of 30%; AGC target 1e4; rapid scan mode; max injection time of 50 ms, isolation window 1.2 m/z). RTS was enabled with full trypsin specificity, max. 2 missed cleavages, max. search time of 40 ms, max. 2 variable modifications, max. 10 peptides per protein, and considering the following modifications: Carbamidomethyl on cysteine and TMT-6-plex on Lys and peptide N-terminus as static, oxidation on Met and deamidation on Asn and Gln as variable modifications. The scoring thresholds were applied (1.4 for Xcorr, 0.1 for dCn, 10 ppm precursor tolerance) and the Homo sapiens Uniprot database (release 2020.01) was used for the search. Quantitative SPS-MS3 scans were performed in the Orbitrap with the following settings: HCD normalised collision energy 50%, resolution 50,000; AGC target 1.5e5; isolation window 1.2 m/Z, MS2 isolation window 3 m/z, 10 notches, max injection time of 150 ms. Total cycle time for each CV was set to 1.2 sec.

DDA-MS2 method (acetylome and phosphorylome analysis) was performed on an Orbitrap Exploris 480 mass spectrometer (Thermo Fisher) and similar HPLC and nanospray settings as described above, with the exception of a FAIMS Pro ion source and that 3 hours gradient time was applied. MS1 survey scans were recorded with the following parameters: resolution 60,000, scan range 375-1,500 m/z, automatic gain control (AGC) target = Custom, and maximum injection time (IT) mode = Auto, FAIMS voltages on, FAIMS CV -40V, -55V and -70V, dynamic exclusion was set to 45 sec. MS2 analysis was performed with the following parameters: resolution 45,000, normalised AGC Target 200%, AGC target = Custom, maximum IT mode = Custom, isolation window 0.7 m/z, and HCD normalised collision energy of 34%.

#### Mass spectrometry data analysis with MaxQuant

MS raw files were split according to CVs (-40V, -55V, -70V) using FreeStyle 1.7 software (Thermo Scientific). All raw MS data were commonly analysed using MaxQuant (37) software version 1.6.17.0, using default parameters with the following modifications. Proteome (MS3), acetylome and phosphorylome (MS2) raw data were included in the same search but in separate parameter groups. MS2 and MS3 spectra were searched against the Homo sapiens (human) Uniprot database (release 2020.01; with isoforms) and a database of common laboratory contaminants. Enzyme specificity was set to “Trypsin/P”, the minimal peptide length was set to 7 and the maximum number of missed cleavages was set to 2 for proteome and phosphorylome data and to 4 for the acetylome data. Carbamidomethylation of cysteine was searched as a fixed modification. “Acetyl (Protein N-term)”, “Oxidation (M)” were set as variable modifications. A maximum of 5 variable modifications per peptide was allowed. For search groups containing acetylome and phosphorylome files “Acetyl (K)” and “Phospho (STY)” were, respectively, set as variable modification. The identification and quantification information of acetylated peptides, sites, and proteins was obtained from the MaxQuant “Acetyl (K) Sites”, “Phospho (STY) Sites”, “ModificationSpecificPeptides”, and “ProteinGroups” tables.

Data were analysed in R (4.1.0) using custom scripts (Supplemental Material). The analysis procedure covered: correction for isotopic impurities of labels, within-plex median normalisation, between-plex IRS-normalisation (38), and statistical between-group comparisons using LIMMA (3.50). To account for differences of protein abundance on lysine acetylation-site level, site intensities were normalised to protein intensities before group comparisons with LIMMA.

### Statistical analysis and data visualisation

Perseus (version 1.6.15.0, (39)) Python (version 3.9) and R (version 4.0.4) were used for statistical analysis and visualisation of graphs and plots. GraphPad Prism version 8.0.1 for Windows (GraphPad Software, San Diego, California, USA) was used to generate bar graphs and viability curves in Figures 2 and 3 and Supplemental Figures 2 and 5. In case of histone deacetylase assays, immunofluorescence quantification or viability, unpaired parametric t-test was always used for comparison of one sample to wildtype.

**Figure 2.**
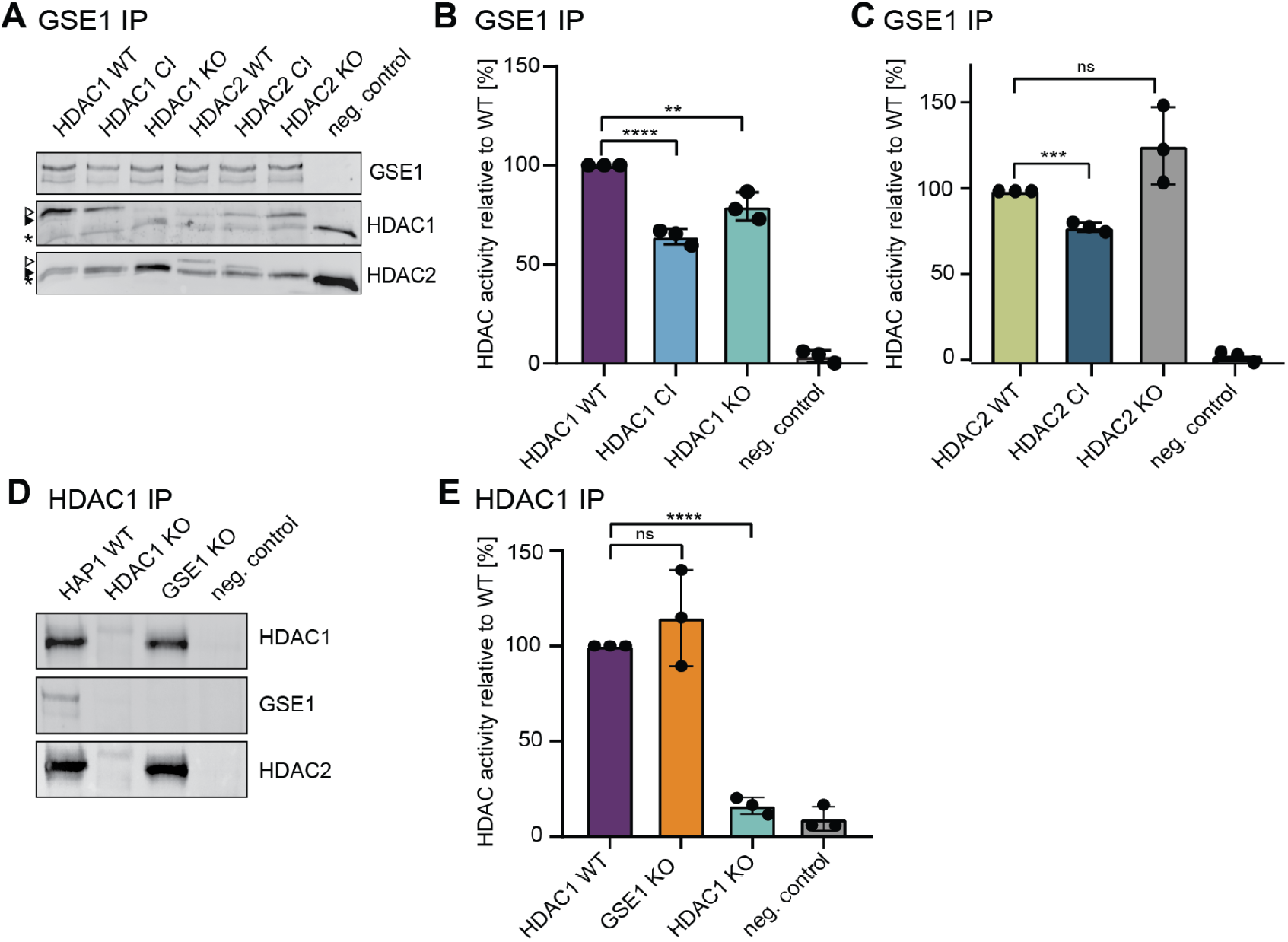
GSE1 as part of HDAC1/2 enzymatically active deacetylase complexes. **(A)** Western blot showing immunoprecipitated GSE1 complexes in HAP1 cells expressing HDAC1 and HDAC2 variants. Negative control: pull-downs performed using anti-mouse IgG antibodies. Antibodies used for detection are depicted on the right. Asterisk: unspecific band. White arrow: transgenic flag-tagged HADC1 or HDAC2 Black arrow: endogenous HADC1 or HDAC2 **(B)** Deacetylase activity measurements of GSE1-HDAC1/2 containing complexes. Panel shows the impact of HDAC1 deletion and inactivation on deacetylation activity. The data represent the mean values ± standard deviation (SD) of 3 biological replicates. Deacetylation activity obtained with HDAC1 WT extracts was set to 100%. Significance was determined by unpaired parametric t-test by comparison to wildtype. *p < 0.05, **p < 0.01, ***p < 0.001. **(C)** The impact of HDAC2 deletion and inactivation was monitored. Panel shows the impact of HDAC2 deletion and inactivation on deacetylation activity. The data represent the mean values ± standard deviation (SD) of 3 biological replicates. Deacetylation activity of HDAC2 WT extracts was set to 100%. Significance was determined by unpaired parametric t-test by comparison to wildtype. *p < 0.05, **p < 0.01, ***p < 0.001. **(D)** Western blot illustrating immunoprecipitated endogenous HDAC1 in HAP1 WT, GSE1 KO or HDAC1 KO cells. Antibodies used for detection are depicted on the left. Negative control: pull-downs performed using anti-rabbit IgA antibodies. **(E)** The impact of GSE1 deletion was monitored. The data represent the mean values ± standard deviation (SD) of 3 biological replicates. Deacetylation activity obtained with HAP1 WT extracts was set to 100%. Significance was determined by unpaired parametric t-test by comparison to wildtype. *p < 0.05, **p < 0.01, ***p < 0.001.

**Figure 3.**
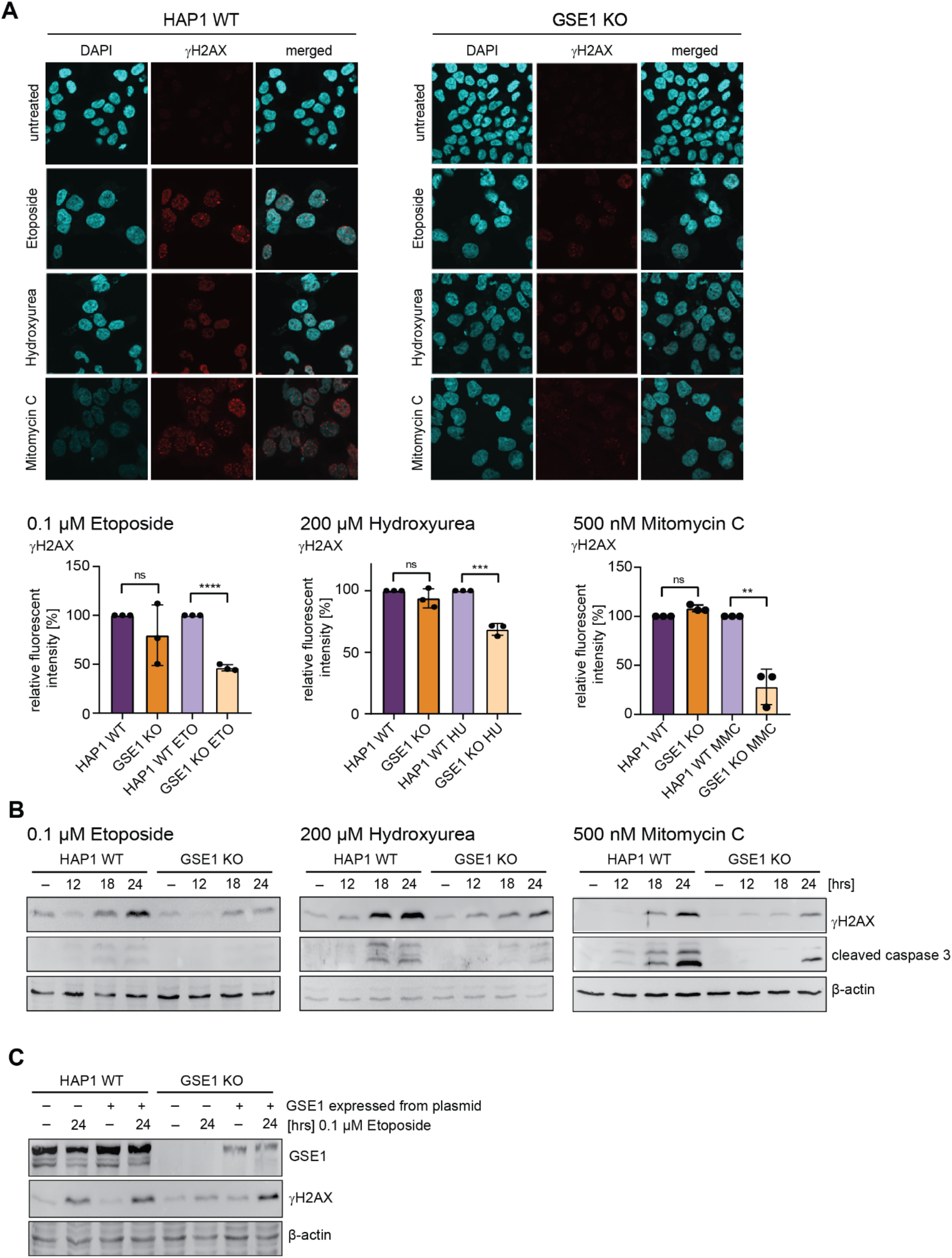
Loss of GSE1 leads to impaired DNA damage response signalling. **(A)** Phosphorylation of H2AX histone (γH2AX) was analysed by immunofluorescent staining for γH2AX. The nucleus was stained with DAPI. Relative fluorescent intensity was plotted, and significance determined by unpaired parametric t-test by comparison to wildtype cells. *p < 0.05, **p < 0.01, ***p < 0.001. HAP1 WT cells were set to 100% as a control. **(B)** Downregulation of γH2AX in GSE1 KO was confirmed by western blot analysis upon treatment for 12, 18 and 24 hours with etoposide, mitomycin C or hydroxyurea. Apoptosis was determined by cleavage of caspase 3 and actin was used as loading control. **(C)** Phosphorylation of H2AX histone (γH2AX) was analysed by western blot upon re-introduction of GSE1 in untreated or 24 hours etoposide treated HAP1 WT and GSE1 KO cells. Apoptosis was determined by cleavage of caspase 3 and actin was used as loading control. GSE1 expression was analysed using GSE1 specific antibody.

For OMICS data, the individual statistical analysis is indicated in corresponding method parts. Heatmaps shown in Figures 4 and 5 and Supplementary Figures 3 and 4 were created in Perseus (version 1.6.15.0), for clustering the k-means algorithm was used. TMT reporter ion intensities were log2 transformed and z-scored. Mapping of human to mouse proteins was done using the ortholog database from OrthoInspector 3.0 (database version 2019) (40), proteins with no annotated ortholog between human and mouse were excluded from the analysis (Supplementary Figure 1). For bar plots shown in Figure 4C and 5B TMT reporter ion intensities were log2 transformed and z-scored using inhouse Python scripts. Bands of Western blot images were cut out using Adobe Photoshop version 10.0 and figures were prepared using Adobe Illustrator version 16.0.0.

**Figure 4.**
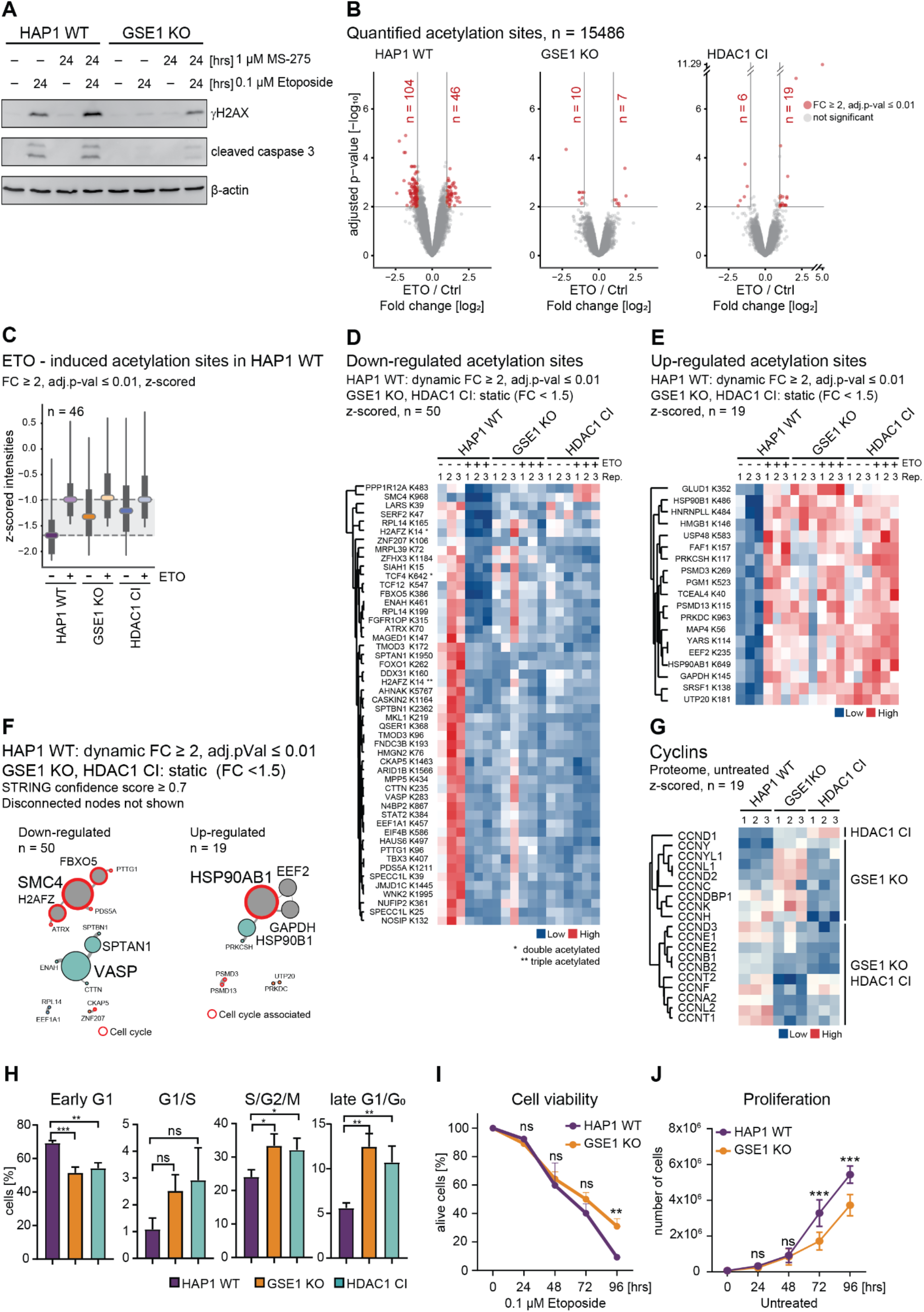
GSE1 KO and HDAC1 CI show distinct acetylation and protein signatures affecting cell cycle. **(A)** Western blot illustrating synergistic effect of MS-275 and etoposide on DNA damage induction (γH2AX) in HAP1 WT and GSE1 KO cells. Actin was used as a loading control. **(B)** Volcano plots displaying acetylome changes in HAP1 WT, HDAC1 CI and GSE1 KO cells upon 6 hours treatment with 0.1 µM etoposide. Significantly upregulated and downregulated acetylation sites are shown in red (≥ 2-fold change over untreated control, adjusted p-value ≤ 0.01). **(C)** Boxplots of z-scored acetylation site intensities in different cell lines and conditions of sites that are etoposide induced in HAP1 WT cells. Coloured horizontal lines represent medians, boxes represent the 25th and 75th percentiles, whiskers expand to the 5th and 95th percentile, outliers are not shown. **(D-E)** Heatmaps showing acetylation profiles of selected downregulated and upregulated sites in HAP1 WT upon etoposide treatment that are not responsive in GSE1 KO and HDAC1 CI mutants. Corresponding gene names and positions of acetyl-lysine are indicated on the left. * and ** mark quantification of acetylation sites arising from *dual or **triple acetylated peptides where only a single site was confidently assigned. **(F)** STRING DB-based network of proteins associated with etoposide-responsive acetylation sites in HAP1 WT cells that are stable in GSE1 KO and HDAC1 CI. High-confidence interactions (confidence score ≥ 0.7) were selected. Disconnected nodes are not shown. Colouring of nodes is according to modularity classes. Proteins functionally associated with cell cycle regulation are highlighted in red. Strength of the associations is represented by the thickness of the edges. **(G)** Heatmap illustrating protein abundance of selected cyclins in HAP1 WT, GSE1 KO and HDAC1 CI. Corresponding gene names are indicated on the left. **(H)** Quantified cell cycle phase distribution of HAP1 WT, HDAC1 CI and GSE1 KO FUCCI cells using flow cytometry. The data represent the percentage of cells in each cell cycle phase ± standard deviation (SD) of 3 biological replicates. Significance is determined by unpaired parametric t-test by comparison to wildtype. *p < 0.05, **p < 0.01, ***p < 0.001. **(I)** Cell viability was calculated by counting of the cells every 24 hours cultured in the presence of etoposide up to 96 hours. Significance is determined by unpaired parametric t-test by comparison to wildtype. *p < 0.05, **p < 0.01, ***p < 0.001. **(J)** Cell proliferation was calculated by counting of the cells every 24 hours cultured in full media. Significance is determined by unpaired parametric t-test by comparison to wildtype. *p < 0.05, **p < 0.01, ***p < 0.001.

**Figure 5.**
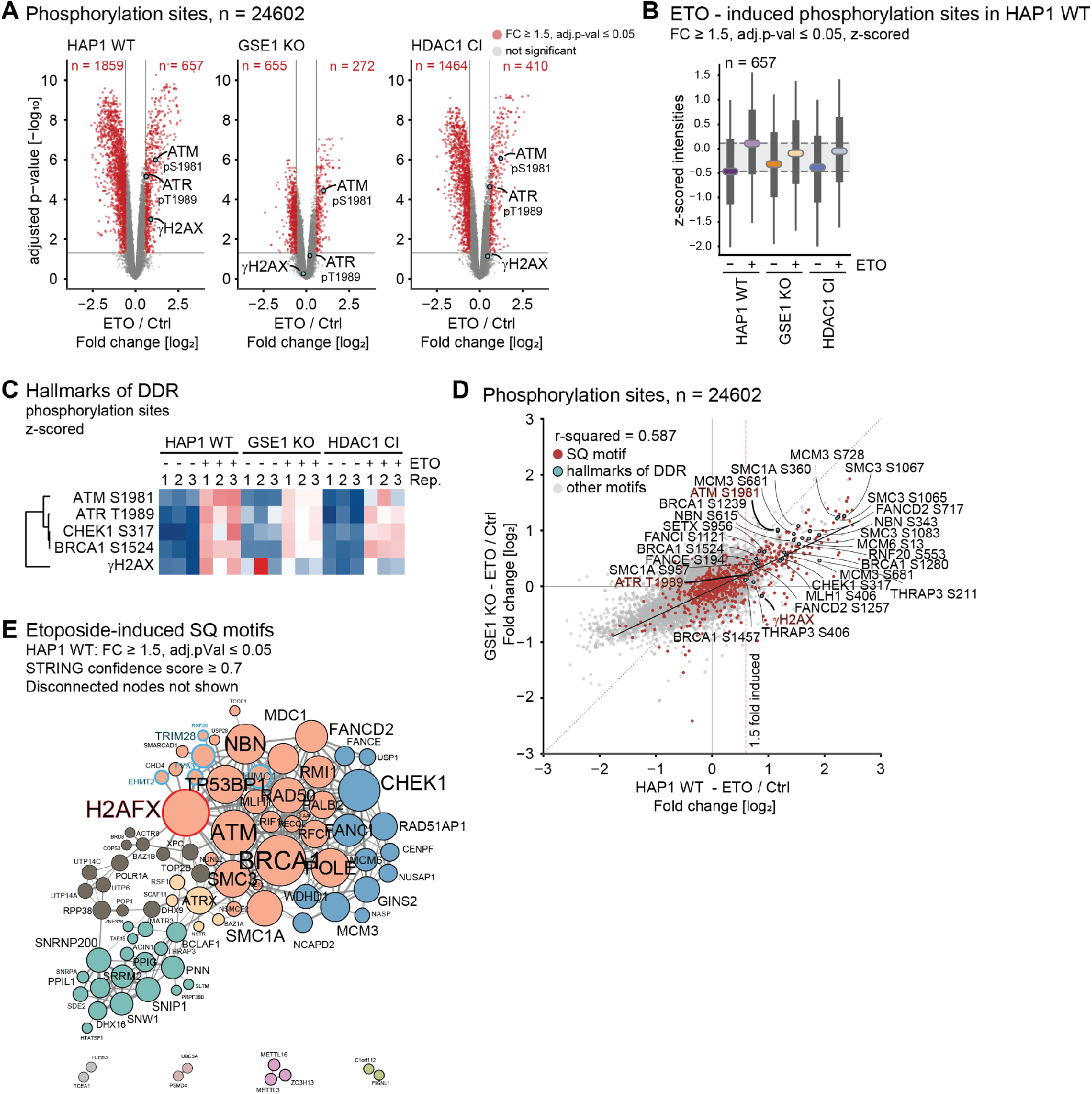
Global impact of GSE1 ablation on DNA damage sensing and downstream phosphorylation signalling. **(A)** Volcano plots displaying phosphorylome changes in HAP1 WT, HDAC1 CI and GSE1 KO cells upon 6 hours treatment with 0.1 µM etoposide. Significantly upregulated and downregulated phosphorylation sites are shown in red (≥ 1.5-fold change over untreated control, adjusted p-value ≤ 0.05). The DDR hallmark modifications γH2AX (representing pS139 of histone variant H2AX) and autophosphorylation of ATM (pS1981) or ATR (pT1989) are highlighted in blue. **(B)** Boxplots of z-scored phosphorylation site intensities in different cell lines and conditions of sites that are etoposide induced in HAP1 WT cells. Coloured horizontal lines represent medians, boxes represent the 25th and 75th percentiles, whiskers expand to the 5th and 95th percentile, outliers are not shown. **(C)** Heatmap displaying phosphorylation profiles of selected hallmarks of the DDR in HAP1 WT, GSE1 KO and HDAC1 CI. Corresponding gene names and positions of phosphorylated residues are indicated on the left. **(D)** Scatter plot comparing etoposide induced phosphorylation between HAP1 wildtype cells and GSE1 KO cells. Sites harbouring an SQ motive are shown in red, sites representing hallmarks of the DDR are shown in blue and annotated. **(E)** STRING DB-based network of proteins associated with etoposide-responsive SQ-motives in HAP1 WT. High-confidence interactions (STRING confidence score ≥ 0.7) were selected. Disconnected nodes are not shown. Colouring of nodes according to modularity classes. Proteins functionally associated with histone modification are highlighted with blue frames. yH2AX is highlighted in a red frame. Strength of the associations is represented by the thickness of the edges.

### Integration of published mass spectrometry datasets

For integration of published datasets, we used in-house Python scripts. The list of previously described HDAC1 interactors in CEM T cells was obtained from Joshi *et al.* (41) “Supplementary Table 2”, by selecting entries with a value greater than 0.9 in the “HDAC1_Prob” column, as described by the authors.

SILAC ratios of acetylome and phosphorylome changes in response to 10 µM etoposide treatment for 24 hours in U2OS cells were obtained from Beli et al., 2012 (42). Phosphorylation, acetylation, and protein SILAC ratios were extracted from the “Etoposide” sheet in “Table S1”, “Table S2” and “Table S3” using the column “Ratio H/L Etoposide” or “Ratio H/L Etoposide Combined”. SILAC ratios of phosphorylation and acetylation sites were normalised to protein SILAC ratios, sites of not quantified proteins were discarded. Phosphorylation and acetylation sites were mapped to the Homo sapiens (human) Uniprot database (release 2020.01) by using the “Gene Names”, “Position”, and “Sequence Window” columns, and sites with ambiguous mapping were removed. Normalised SILAC ratios were then integrated into Supplementary Tables 3 and 4 as column “Ratio H/L etoposide [log2]”.

### Protein network analysis

Protein-protein interaction networks were generated using the STRING database v11.5 (43), network parameters were exported and modified as follows: column “node1” was defined as “source”, column node 2 was defined as “target” and “combined STRING score” was defined as weight of the network edge. Network type was undirected. The modified table was imported into Gephi v0.9.7 (44) for network visualisation. Node sizes were set proportional to the degree value and edge thickness to the combined interaction score. Network was visualised using the Fruchteman Reingold function with default settings and ForceAtlas 2 function (45) using balanced scaling and gravity settings, and the “prevent overlaps” option being activated. All other settings were default. Nodes were either coloured according to the classification presented in (41) (Figure 1B and Supplemental Figure 1B) or according to modularity class (Figures 5E and 6C).

**Figure 6.**
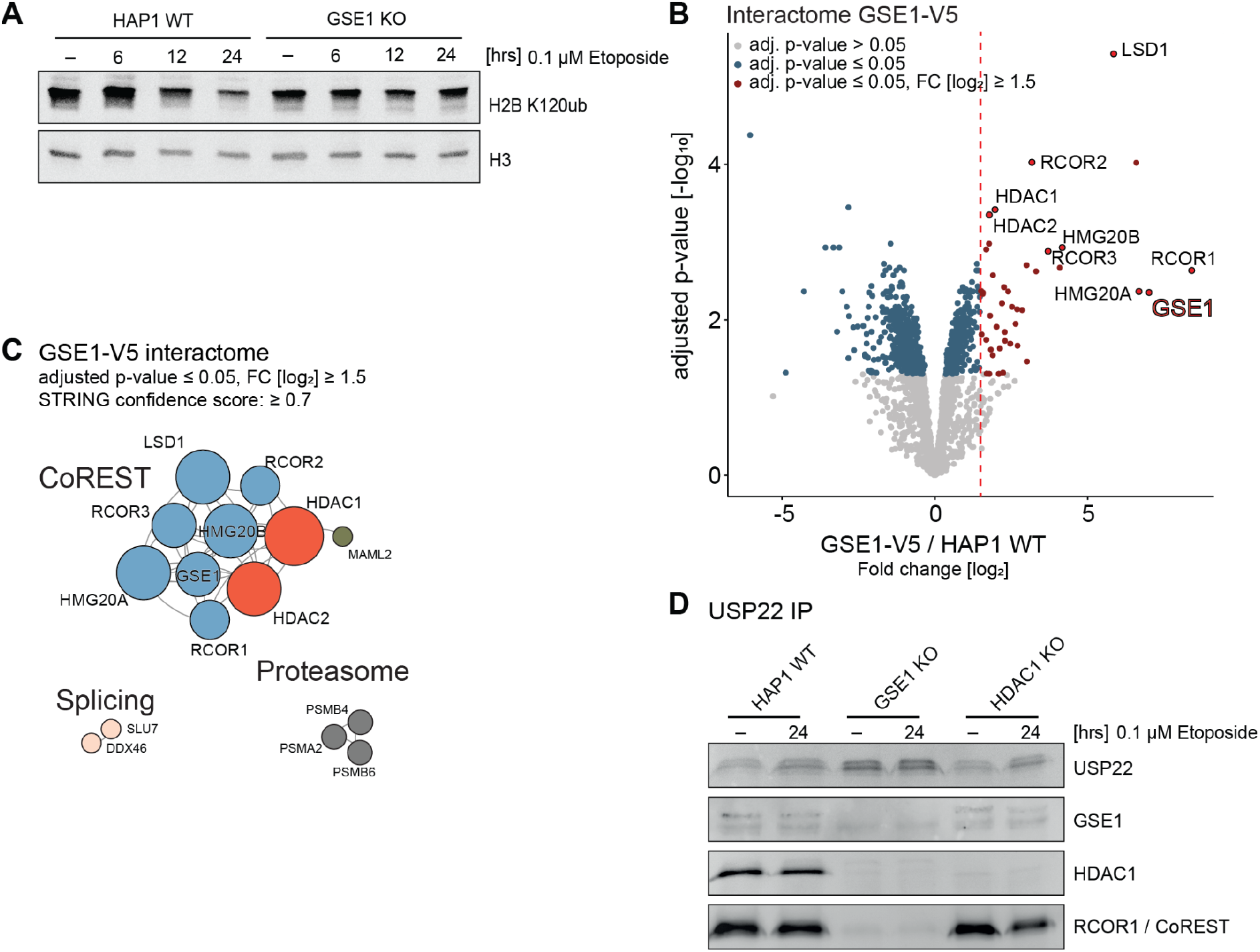
Identification of specific GSE1/CoREST complex with multi-enzymatic eraser activities. **(A)** Histone western blot detecting levels of ubiquitination of H2B at K120 upon treatment with etoposide for 6, 12 and 24 hours. C-terminal histone H3 was used as a loading control. **(B)** Volcano plot displaying the interactome of GSE1 in HAP1 cells determined by AP-MS. Significant interactors (≥ 1.5-fold change [FC] over wildtype cells [dashed line], adjusted p-value ≤ 0.05) in GSE1 V5-tagged cells over HAP1 WT cells are shown with red dots. Gene names of known subunits of the HDAC1/CoREST complex are highlighted, GSE1 is highlighted with bold red text. **(C)** STRING DB-based protein interaction network of the GSE1 interactome. High-confidence interactions (confidence score ≥ 0.7) were selected. Strength of the associations is represented by the thickness of the edges. **(D)** Western blot depicting immunoprecipitated endogenous USP22 in HAP1 WT, HDAC1 KO and GSE1 KO cells. Antibodies used for detection are depicted on the right.

## Results

### GSE1 is a prominent interactor of HDAC1

The class I histone deacetylase HDAC1 is the catalytic component of several multiprotein co-repressor complexes, present in virtually all mammalian cells. To gain a deeper understanding of the functional landscape of HDAC1, we examined the composition of HDAC1-associated proteins in different cellular backgrounds. First, we affinity purified (AP) C-terminally FLAG-tagged HDAC1 from the nearly haploid human cell line HAP1 (36,46) and analysed the interactome using quantitative mass spectrometry (MS). In total, we identified 161 proteins as potential HDAC1 interaction partners [logFC > 1.5, adjusted p-value < 0.05], thereby providing a comprehensive analysis of the interactions of the enzyme (Figure 1A, Supplementary Table 1).

The presence of well-known subunits of co-repressor complexes such as NuRD, Sin3-related, MiDAC or CoREST (41) in the HDAC1 interactome indicated that our analysis adequately covered anticipated binding partners of the deacetylase. In addition, we found the HDAC1 substrates H2AFV, H2AFX (47) and the prefoldin/CTT complex (48) among the co-precipitated proteins. We also confirmed interaction between HDAC1 and a group of importins, zinc finger proteins or transcriptional regulators possessing an ELM-SANT domain (Figure 1B).

When comparing our results to those obtained from a similar approach in human CEM T-cell lines (41), we observed a high level of overlap in the identified binding partners of HDAC1 (Supplementary Figure 1A). 78 of the 91 HDAC1-interacting proteins reported by Joshi *et al* were also identified in the HAP1 dataset. This suggests stable HDAC1-complex composition across different cell types. However, we have covered additional 83 proteins as HDAC1 interactors that were not reported by Joshi *et al*, providing an in-depth view on potential HDAC1 binders and cell type specific differences. Among these we found many transcriptional regulators such as TFDP1, MAX, ZEB2, CTBP2 and MECOM, and negative regulators of ubiquitination BAG2, OGT or SQSTM1 (Figure 1B).

To examine cross-species conservation in the composition of HDAC1 complexes we additionally analysed HDAC1-FLAG purifications from mouse embryonic fibroblasts (MEFs) (Figure 1C and Supplementary Figure 1B, Supplementary Table 2). Again, we observed a high level of overlap (68 proteins) in the identified interaction partners. Those in common contained mainly known corepressors subunits, as expected. Interestingly, a wide fraction of MEF-specific HDAC1 binders contained proteins involved in proteasomal degradation, transport or mitochondria related processes (Supplementary Figure 1C, Supplementary Figure 1D). We thus recovered generally accepted hallmarks of the HDAC1 interactome and demonstrate data consistency with previous studies. Additionally, we provide a view on HDAC1 interactome across different species and cell types showing a high level of conservation among core corepressor complex subunits as well as variations specific for species or cell type. It is worth mentioning that our findings align closely with those of a comparable study on the HDAC1 interactome in Th1 T-cells (Ci, Stolz *et al*., manuscript in preparation).

Remarkably, a previously reported interaction between HDAC1 and GSE1 (41,49-52), that has been not fully appreciated so far, was found across all experimental conditions tested ([fold change [log_2_] = 12.37, adjusted p-value < 0.001], Figure 1A, Figure 1B Supplementary Table 1, Supplementary Table 2). We additionally confirmed this interaction by co-IP experiments in combination with western blot analysis in murine Th1 T-cells (Figure 1D), thereby also confirming cross-species conservation of a GSE1-HDAC1 complex. Since the binding of GSE1 to HDAC1 appeared prominent we decided to characterise this interaction in more detail.

We first investigated whether GSE1 binding possibly affects the enzymatic activity of HDAC complexes. To this end, we employed and expanded a toolbox of transgenic human tumour cell lines (46), previously created in our research group (36). In particular, we used HAP1 cells expressing either C-terminally FLAG-tagged wildtype (WT) or catalytic inactive (CI) mutants of HDAC1 or its paralog HDAC2 in the respective knockout background (Supplementary Figure 2A and Supplementary Figure 2B). Using a GSE1 specific antibody, we purified GSE1-containing HDAC complexes from transgenic HAP1 cells and determined the associated deacetylase activity using H^3^-acetylated histones as substrate (36,53). We observed a robust deacetylase activity in cells expressing wildtype HDAC1 (HDAC1 WT) as well as HDAC2 (HDAC2 WT), demonstrating that GSE1-containing HDAC complexes are active (Figure 2A). Moreover, this activity was significantly reduced in cells expressing catalytically inactive HDAC1 (HDAC1 H141A, HDAC1 CI) allele, but not in cells expressing catalytically inactive HDAC2 (HDAC2 H142A, HDAC2 CI). HDAC1 KO cells showed only mild reduction in deacetylase activity, whereas HDAC2 KO showed increased activity when compared to HDAC2 WT (Figure 2B and 2C). This is most likely due to a compensatory effect of upregulated HDAC1 in the absence of HDAC2 (36). In summary, these data suggest that HDAC1 is the main deacetylase of GSE1-containing complexes.

Next, we used GSE1 KO cells to test whether loss of GSE1 affects HDAC1 and HDAC2 deacetylase activity. Activity assays of HDAC1- or HDAC2-associated complexes, isolated from wildtype and GSE1 KO cells, showed no significant difference (Figure 2D and 2E and Supplementary Figure 2C), suggesting that deacetylase activity is not dependent on GSE1. As expected, an HDAC1 KO had no effect on the activity of HDAC2-associated complexes whereas HDAC1-associated complex activity was abrogated in HDAC1 KO cells (Figure 2E and Supplementary Figure 2C).

In summary, we find that HDAC1 interacts with known co-repressor complexes and various other proteins, among which GSE1 was identified as a prominent binding partner within different cell types (HAP1, MEFs and Th1 T-cells). In addition, we demonstrate that GSE1-HDAC1/HDAC2 complexes are catalytically active, and their deacetylase potential is dependent on HDAC1 and HDAC2 activity. Moreover, GSE1 ablation has no impact on deacetylase activity of HDAC1 and HDAC2.

### GSE1 KO leads to reduced response to DNA damage

A genetic screen performed by Gudkov *et al.* was first to identify GSE1 as a potential mediator of cellular sensitivity to the topoisomerase II inhibitor etoposide (54). However, this study did not investigate the molecular function of GSE1. To test whether GSE1 plays a role in the DDR, we treated HAP1 wildtype and GSE1 knock-out (KO) cells with etoposide for 24 hours and determined the level of phosphorylation at S139 of histone H2AX (designated as γH2AX), a marker of DNA double strand breaks. Using indirect immunofluorescence microscopy and western blot analysis, we observed an increase in γH2AX in wildtype cells upon etoposide treatment. This increase, however, was severely reduced in GSE1-negative cells (Figure 3A and 3B). In addition, we followed caspase 3 cleavage to determine the level of apoptotic cells by western blot analysis in cells treated with etoposide for 12, 18 and 24 hours. We detected reduced cleavage of caspase 3 in GSE1 KO cells compared to the control, indicating a reduced level of DNA damage induced apoptosis in GSE1 KO cells (Figure 3B). Moreover, complementation of GSE1 fully restored etoposide-induced γH2AX formation upon 24 hours of etoposide treatment (Figure 3C).

Etoposide is known to form a complex with topoisomerase II that inhibits DNA synthesis and in turn induces DNA damage via double strand breaks (55). There are, however, various types of DNA damage that can be repaired by different mechanisms during different stages of the cell cycle (56). To test whether deletion of GSE1 also affects other repair mechanisms that are not induced by DNA double strand breaks, we used the alkylation reagent mitomycin C, as well as the DNA synthesis inhibitor hydroxyurea as alternative DNA damage inducers. Similar to the effect of etoposide treatment, we observed a significant reduction in γH2AX formation and caspase 3 cleavage in GSE1 KO cells upon treatment with mitomycin C or hydroxyurea (Figure 3A and 3B).

These data strongly suggest that GSE1 plays a central role in γH2AX formation independent of the type of DNA damage.

### DNA damage does not impact GSE1-HDAC1 complex deacetylase activity

Increased sensitization of cancer cells with decreased homologous recombination repair capacity has been observed upon HDAC inhibition in combination with antitumor drugs (18,57-59). Specifically, inhibition of class I HDACs by high dosage of Entinostat (MS-275) induced γH2AX, significantly inhibited cellular growth and triggered apoptosis (60-63). To avoid side effects caused by the high dose of HDAC1 inhibitor (64), we decided to apply a mild dose of MS-275 (1 μM) that functionally inhibits class I HDACs yet does not cause detrimental DNA damage. We tested whether HDAC inhibition has a synergistic effect with etoposide on the DDR in our cellular system and whether this response is altered by loss of GSE1. As expected, we could not detect a significant induction of γH2AX with MS-275 alone in HAP1 WT or GSE1 KO cells (Figure 4A). However, in combination with etoposide, γH2AX signal became increased in both cell lines when compared to etoposide alone pointing towards synergistic influence on DDR induction. Interestingly, these data show that inactivation of HDACs by MS-275 partly restores γH2AX signalling in GSE1 KO cells. A possible explanation for this is that GSE1 may negatively regulate the activity of HDAC1 in the context of DNA damage. To address this, we treated HAP1 wildtype cells for 6 hours with etoposide, immunoprecipitated GSE1-HDAC1 complexes and measured deacetylase activity. We did not detect any changes in activity of GSE1-HDAC1 complexes upon etoposide treatment (Supplementary Figure 2D). This suggests that the enzymatic activity of GSE1-HDAC1 complexes is rather stable upon DDR induction, and that GSE1 possibly affects γH2AX formation independent of changes in HDAC1 activity.

### GSE1 does not regulate the DNA damage response via acetylation

DDR is vastly orchestrated by a sequential cascade of multiple signalling events that are tightly regulated. In addition, a functional role between phosphorylation and acetylation crosstalk has been established in cardiomyocytes where inhibition of HDACs can lead to increased phosphorylation of proteins such as p38, ERK1, and AKT (reviewed in (65)).

To further investigate how HDAC1 and GSE1 can regulate DDR and whether their mode of action might be overlapping, we next performed a proteomic analysis using quantitative mass spectrometry based on the recently developed TMTpro 16-plex isobaric labelling technology (66). We determined global proteome, acetylome and phosphorylome profiles in HAP1 WT, GSE1 KO and HDAC1 CI cells and compared patterns between untreated and etoposide-treated (0.1 µM) cells (Supplementary Figure 3A, Supplementary Table 3 and 4). To avoid adaptive responses and bystander effects, etoposide treatment was limited to 6 hours. In total, we covered 9051 proteins, 15,486 acetylation sites and 24,602 phosphorylation sites in our quantitative analysis. Changes in phosphorylation and acetylation site abundance were further normalised to the corresponding protein levels. Principal component analysis of the proteome, acetylome and phosphorylome separated out distinct clusters for wildtype, GSE1 KO and HDAC1 CI cells, pointing towards quantitative differences between the genetic backgrounds and indicating a high level of reproducibility between replicates (Supplementary Figure 3B). Phosphorylation sites showing a fold change of ≥ 1.5 over wildtype cells and an adjusted p-value of ≤ 0.05 were considered as significantly regulated. However, a higher level of variance was observed in untreated acetylome samples. We therefore used more stringent filtering for the acetylome dataset (fold change ≥ 2 over wildtype cells, adjusted p-value ≤ 0.01).

First, we investigated etoposide-induced acetylome changes in HAP1 WT, GSE1 KO and HDAC1 CI cells. We detected a rather small population of increased acetylation of sites (46) in HAP1 WT upon etoposide treatment, which is in accordance with the previous studies, reporting only minor changes in acetylation levels upon 24 hours of etoposide treatment (42) or in response to irradiation (67). The number of acetylation sites affected by etoposide treatment was reduced in both HDAC1 CI (19 sites) and GSE1 KO (7 sites) cells (Figure 4B). Next, we analysed how the mutants affect the set of DNA damage-induced acetylation sites. We selected the set of 46 sites induced in the wildtype and studied their behaviour at basal level and after etoposide treatment. After etoposide treatment, acetylation of those sites reached a similar level in the mutants as in the wildtype. When looking at untreated conditions, however, this set of sites displayed elevated acetylation levels in both, HDAC1 CI as well as in GSE1 KO cells, (Figure 4C), indicating that both mutants cause dysregulation in DDR signalling. Therefore, we tested whether both mutants had the same effect on the global acetylome. When comparing global acetylation patterns of HDAC1 CI to GSE1 KO cells, we found 102 sites of different abundance (fold change ≥ 2, adjusted p-value ≤ 0.01), suggesting that the mutants have to a certain extent individual effects on the acetylome (Supplementary Figure 3C). In line with this, recently described substrate sites of HDAC1 (36) show higher acetylation levels in the HDAC1 CI mutant (Supplementary Figure 3C), which also supports our observation that GSE1 does not affect HDAC activity (Figure 2D). We next wanted to get an insight into DNA damage-induced processes that are commonly dysregulated in GSE1 KO and HDAC1 CI. We focussed on acetylation sites that are etoposide-responsive (fold change ≥ 2, adjusted p-value ≤ 0.01, in wildtype cells) in wildtype cells but remain unaffected in both mutants (Figure 4D and 4E). The set of corresponding proteins was submitted to a protein network analysis using the STRING database tool (43). We found that proteins harbouring these acetylation sites are part of cell cycle control networks (Figure 4F), pointing to an affected cell cycle regulation in these mutants. Based on this finding, we asked whether protein levels of cyclins and cyclin dependent kinases are altered in the mutants. Indeed, in both mutants we observed strong differences in the protein abundance of several cyclins known to regulate cell cycle transition points (68-71) (Figure 4G). Decreased abundance of cyclin dependent kinases CDK1, CDK2, and CDK6 was observed in GSE1 KO as well as HDAC1 CI cells (Supplementary Figure 3E). To monitor potential changes in cell cycle distribution, we used the fluorescent ubiquitination cell cycle indicator (FUCCI) system developed by Sakaue-Sawano *et al.* (32). We observed a similar cell cycle phase distribution between GSE1 KO and HDAC1 CI cells that was different from HAP1 WT cells (Figure 4H, Supplementary Figure 3F). GSE1 KO and HDAC1 CI showed an increased fraction of cells in S/G2/M and G1/G0 phase and a reduced number of early G1 cells compared to HAP1 WT (Figure 4H). In response to treatment with etoposide, the wildtype and both mutants showed an increase in S/G2/M phase (Supplementary Figure 3F), which is in line with previous reports (72-74). Based on these findings, we wanted to clarify whether deletion of GSE1 leads to an accumulation of G2 cells, which could indicate DNA repair incompetence. Cells were stained for phosphorylation of histone H3 at Ser10 that is known to correlate with chromosome condensation during cell division. Quantifying the percentage of cells in G2 and M phase revealed a significantly increased number of GSE1 KO cells in G2 phase when compared to HAP1 WT (Supplementary Figure 3G). These data indicate that the incompetent DNA repair in GSE1 KO cells causes a G2/M arrest. We also observed a decreased proliferation rate in the GSE1 KO compared to HAP1 WT (Figure 4I). In addition, when cultivating the cells in etoposide containing media, GSE1 KO cells acquired etoposide resistance and restored their growth after 96 hours in presence of the drug, whereas the vast majority of the control cells were dead (Figure 4J). These findings are consistent with the previous report from Gudkov *et al.* (54).

Taken together, we find that the overall impact of the DDR on the acetylome seems to be rather neglectable. Etoposide-responsive sites that are commonly affected in both mutants are most likely caused by cell cycle defects. Therefore, we conclude that the mode of action of GSE1 in regulation of DDR is independent of the changes in the acetylome.

### Loss of GSE1 affects early events of DDR and attenuates downstream DNA repair signalling

Since GSE1’s role in response to DNA damage does not depend on protein acetylation, we next examined DNA damage-induced phosphorylation changes after 6 hours of etoposide treatment. In total, we identified 657 (2.7%) up- and 1,859 (7,6%) down-regulated phosphorylation sites in HAP1 WT cells (fold change ≥ 1.5, adjusted p-value ≤ 0.05) in response to DNA damage (Figure 5A). We observed a high level of correlation between our results and those of Beli et al. (42), monitoring etoposide-induced phosphorylation patterns 24 hours after induction with etoposide in U2OS cells (Supplementary Figure 4A). Notably, 53.6% of commonly induced sites constitute serine glutamine (SQ) motifs, which are putative ATR / ATM substrates and a hallmark of the DDR (75). About a quarter of the commonly affected SQ sites are found on DDR-related proteins such as BRCA1, FANCI, MCM6, MLH1, NBN and SMC1A. We also confirmed the etoposide-induced phosphorylation of THRAP3 serine 211, which was identified as a novel DDR regulator by Beli et al. (42). Phosphorylation sites that were only affected after 24 hours of etoposide treatment might correspond to downstream effects of the DDR or secondary effects caused by prolonged DNA damage.

Next, we focused on how a GSE1 knock-out or loss of HDAC1 catalytic activity affects DNA damage-induced phospho-signalling. We observed a reduced number of etoposide-affected phosphorylation sites in GSE1 KO cells (41.4% upregulated and 35.2% downregulated sites compared to wildtype cells) and to a lesser extent also in the HDAC1 CI mutant (62.4% upregulated, 78.8% downregulated sites compared to wildtype cells) (Figure 5A). Notably, the set of 657 sites that become induced by etoposide treatment in wildtype cells were already slightly elevated in phosphorylation at basal conditions in GSE1 KO and HDAC1 CI cells (Figure 5B), which could be due to a partial constitutive activation of the DDR. However, when the mutants were exposed to etoposide, these sites were not phosphorylated to the same magnitude as in the wildtype. Moreover, both mutants showed a reduced etoposide-induced phosphorylation at several hallmark sites of the DDR, such as γH2AX, pT1989 of ATR, pS317 of CHEK1, when compared to wildtype, indicating that DNA damage-signalling is hampered (Figure 5C and Supplementary Figure 4B). On the other hand, activating autophosphorylation at S1981 of ATM kinase was induced upon etoposide in HAP1 WT and HDAC1 CI but to a lesser extent in GSE1 KO (Figure 5A and Figure 5C). Therefore, we next asked whether the GSE1 KO or HDAC1 CI mutant affects the entire DNA damage-induced phosphorylation response or just a specific set of sites indicative for individual pathways. In fact, the GSE1 KO mutant showed generally hampered phosphorylation at SQ motifs (r-squared = 0.587) and non-SQ motifs in response to etoposide treatment, including the entire set of targets of ATM / ATR kinases (Figure 5D). On a side note, HDAC1 CI showed similar results, albeit to a lesser degree (R-squared = 0.784) and without dysregulation of ATM / ATR kinase regulatory sites (Supplementary Figure 4C). In conclusion, GSE1 appears to play a role in the initial stages of DNA damage detection and recognition, which are essential for full activation of phosphorylation cascades downstream.

To identify such upstream regulators that could be affected by GSE1 we created a STRING database-based protein network of the etoposide phosphorylation response (proteins harbouring SQ motifs, FC ≥ 1.5, padj-value of ≤ 0.05, in wildtype cells). Analysis of the networks revealed proteins mostly associated with DDR and DNA repair, but also with spliceosome, ubiquitination machinery, and histone modifications (Figure 5E). The latter seemed particularly relevant in the context of DNA damage sensing (76), encompassing factors such as EYA3, associated with dephosphorylation of histone H2AX Y142 (77), EHMT2 and TRIM28, both associated with methylation of K9 of histone H3 (78,79), UIMC1, selectively binding K63 of histones H2A and H2AX (80), and RNF20 mediating monoubiquitination of K120 of histone H2B (81). We speculated that ablation of GSE1 might potentially negatively affect the activity of one of these factors or its associated mechanisms. Notably, phosphorylation at S533 of RNF20 was similarly reduced in both GSE1 KO and HDAC CI compared to wildtype, indicating that a fully assembled GSE1-HDAC1 complex is required. Since monoubiquitination/deubiquitination of H2B K120 was previously described as a prerequisite for DDR induction, we turned our attention to this particular lead.

### GSE1 forms a multi-enzymatic COREST subcomplex with deubiquitinases

To test whether H2B K120 monoubiquitination (H2Bub) levels are affected in GSE1 KO cells, we followed this modification in a time-resolved manner using western blotting in combination with an anti-H2Bub antibody. In wildtype cells we observed a prominent and continuous decline in H2Bub levels from the 6 hours to the 24 hours time point of etoposide treatment. Interestingly, in GSE1 KO cells H2Bub levels remained constant throughout the same timeframe (Figure 6A), demonstrating that deubiquitination of H2B K120 is compromised by loss of GSE1. In light of previous findings that deubiquitination of H2B K120 is crucial for DDR induction (11,30), a defect in this step could explain the observed phenotype of GSE1 KO cells. Following this observation, our next goal was to understand if GSE1 directly regulates H2B deubiquitination. To this end, we generated by CRISPR/Cas-9 mediated knock-in a HAP1 cell line expressing a V5-tagged version of the endogenous GSE1 protein and used AP-MS to determine the GSE1 interactome. We purified GSE1 from untreated and 6 hours etoposide treated cells and found the entire set of CoREST complex members to be co-purified (Figure 6B).

Interestingly, GSE1 seems to exclusively bind to the CoREST complex as subunits of other known corepressor complexes such as SIN3, NURD or MiDAC were not enriched by the GSE1 pull down (Supplementary Figure 5A). In addition to CoREST complex members, the proteasome subunits PSMA2 and PSMB4/6, and the splicing factors DDX46 and SLU7 were found to co-purify with GSE1 (Figure 6C, Supplementary Table 5). Notably, we could not detect differences in GSE1-CoREST complex composition in response to etoposide treatment, indicating a stable composition of the complex core that is most likely not affected by DDR induction (Supplementary Figure 5B). As GSE1 is associated with CoREST, we wanted to know whether its deletion affects CoREST-associated deacetylase activity. As expected, we detected no significant change in enzymatic activity associated with CoREST in GSE1 KO cells when compared to wildtype, while HDAC1 ablation significantly reduced CoREST-associated deacetylase activity (Supplementary Figure 5C). Therefore, DNA damage-mediated deubiquitination of H2BBub is not regulated via etoposide-induced changes in GSE1-CoREST complex composition or deacetylation activity.

We did, however, detect traces of UCH proteinases USP22 and USP15 in the affinity purification samples of GSE1-V5, but not in the background controls (see MS/MS counts of USP22 and USP15 in Supplementary Table 5). These low confidence hits were either removed from the dataset or did not pass the applied cut-offs. However, in particular USP22 has been shown to remove monoubiquitin specifically from K120 of H2B during DNA damage (11). We therefore wanted to independently test whether UCH proteinases bind to GSE1 or even to the GSE1-CoREST complex. Indeed, co-IP experiments using USP22 as bait resulted in successful co-isolation of GSE1, HDAC1, HDAC2 and CoREST upon basal conditions and in response to etoposide treatment (Figure 6D). Binding of USP22 to CoREST was independent of HDAC1. However, none of these interactions were observed in GSE1 KO cells, showing that GSE1 is indeed required for the binding of USP22 to the CoREST complex (Figure 6D). We therefore show that GSE1-CoREST complex ultimately combines several enzymatic properties essential to the DDR, such as deacetylation, demethylation, and deubiquitination.

In summary, we propose that GSE1 functions as a scaffolder for assembly of a multi-eraser CoREST complex, harbouring the activities of HDAC1/2, LSD1 and USP22. Presence of USP22 might be key for K120 deubiquitination upon induction of DDR, a primary event in complex PTM rearrangements of histones required for DDR (Figure 7). We thereby provide a first insight into a mechanistic rationale of how the GSE1-HDAC1-CoREST complex modulates the response to DNA damage.

**Figure 7.**
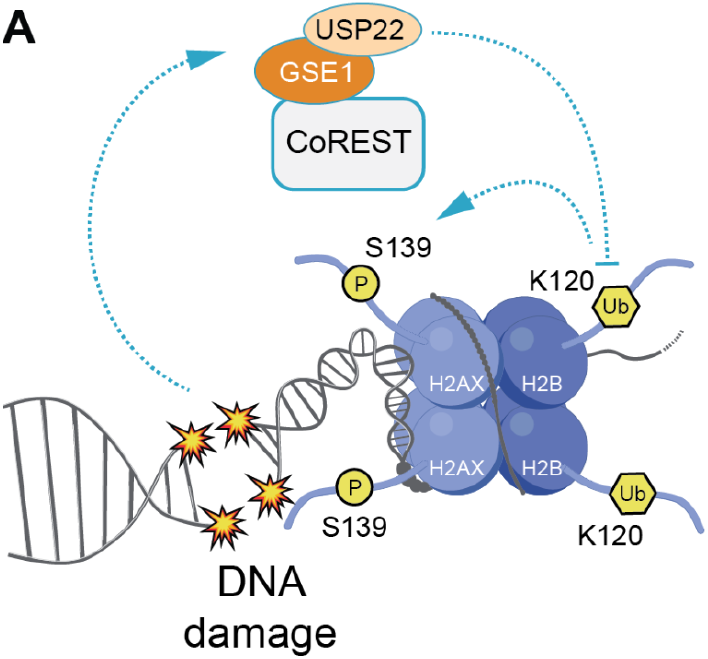
Potential mechanism of DNA damage regulation by GSE1. Proper initiation and orchestration of downstream signalling during DNA damage is highly dependent on the induction of Serine 139 phosphorylation of histone variant H2AX. To sustain proper signalling resulting in efficient DNA repair, a chromatin environment suitable for the recruitment of effector molecules of DNA damage signalling pathway is one of the essential requirements. Here we show a novel role of a neglected protein GSE1, an unit of CoREST/HDAC1 complex, in DDR. We propose that during DNA damage, GSE1/CoREST/HDAC1 complex can bind USP22 that can in return deubiquitinate histone H2B at lysine 120. This eraser event is necessary for sufficient γH2AX signalling as reported previously (11,30). In the absence of GSE1, USP22 cannot bind to GSE1/CoREST/HDAC1 complex and H2B lysine 120 remains ubiquitinated. This inadequate removal of the ubiquitin mark might interfere with γH2AX formation leading to reduced γH2AX signal and downstream signalling.

## Discussion

We have identified GSE1 as an important interactor of HDAC1 as a part of a novel subvariant of the CoREST complex that is involved in regulating the initiation of the DDR. The GSE1/CoREST sub-complex comprises a multi enzymatic eraser combining the deacetylation activity of HDAC1, demethylation activity of LSD1, and deubiquitination activity of USP22. Notably, GSE1 plays a central role in this process, as it constitutively binds to CoREST and is required for USP22 binding. Classical CoREST is primarily associated with silencing of neural genes and has been implicated to govern stem cell differentiation and development (82). These processes are primarily controlled by the enzymatic activities of its known catalytically active subunits HDAC1 and LSD1 (83). Their dysregulation, together with additional CoREST subunits, has been associated with various neurodegenerative diseases as well as cancer (84-86). However, to our knowledge, CoREST has hardly been associated with DNA damage (12) and therefore represents a potentially highly interesting but until now mostly neglected target in the field of medical research.

Despite being largely overlooked in previous research, GSE1 has caught our attention due to its robust interaction with HDAC1 and its previous association with cancer. GSE1 has been identified as a putative subunit of the CoREST and BRAF-HDAC (BHC) complexes (41,49-52). Here, we determine the interactome of GSE1 in human HAP1 cells by direct pull downs and show that GSE1 interacts with core CoREST complex components, including HMB20A/B, HDAC1, HDAC2, RCOR1/2/3, and LSD1, however, we did not find components of other HDAC1 subcomplexes. Of note, we observed that GSE1 is essential for the binding of USP22 to CoREST, suggesting a role as a scaffolder.

Why has this multi-eraser complex gone unnoticed in previous research? We believe that USP22-GSE1-CoREST represents only a small fraction of CoREST complexes. Our IBAQ values (Supplementary Material), providing an estimate of protein abundance, as well as data provided at ProteomicsDB (87), ImmProt (88), and ImmPress (89) report very distinct protein abundance levels for HDAC1, RCOR1, GSE1, and USP22, indicating that these components are not expressed in equimolar levels. This is in agreement with findings of Macinkovic et al., who demonstrated the heterogeneous composition and functional diversity of cellular CoREST complexes (90). Consistent with this, we report that deacetylase activity associated with CoREST is not dependent on the presence of GSE1, implying that GSE1 may not be a constitutive subunit of all CoREST complexes. The GSE1-USP22-CoREST complex seems to be stable, as we do not see changes in its composition following DNA damage induction. These results align with the findings of Barnes *et al*., who observed stable GSE1-CoREST interaction during stem cell differentiation (49).

What is the biological function of the GSE1-USP22-CoREST complex? Here we show that loss of GSE1 results in hampered γH2AX formation, impaired phosphorylation of key sites in DNA damage signalling and an overall compromised DDR. These findings point to a connection between GSE1-USP22-CoREST and the regulation of DNA damage initiation. One of key requirements in this process seems to be the GSE1-dependent binding of USP22 to CoREST. We hypothesise that during DNA damage USP22 is essential for removal of H2B K120ub, which seems to be a prerequisite signal for γH2AX formation (Figure 7). It has been previously reported that B cells lacking USP22 showed impaired removal of H2Bub which caused reduced ATM/ATR and DNA-PK-induced γH2AX formation in antibody class-switch recombination (11,30). In agreement with this we also do not see deubiquitination of H2Bub upon DNA damage in GSE1 KO cells. We suspect that the inefficient removal of H2B K120ub might lead to impaired kinetics of DNA damage-specific chromatin establishment necessary for recruitment of downstream effector molecules. Multiple mechanisms have been discussed in this context, for example, persistent presence of H2Bub could lead to extensive H2A Y142 phosphorylation. Phosphorylation of H2A Y142 in turn interferes with recruitment of MDC1 which is known to shield γH2AX signal from phosphatases (10,11,91). Additionally, it was postulated that H2B K120 undergoes a monoubiquitination to acetylation switch at DNA double strand breaks (DSBs) loci. This switch is needed for relaxation of the chromatin, facilitating resection and accessibility of DNA repair factors. Hence, removal of H2B monoubiquitination is inevitable for proper DDR (76). Notably, our study shows that the function of GSE1 is not limited to DSBs since we observed reduced γH2AX signal upon treatment with various agents including mitomycin C or hydroxyurea (Figure 3A and 3B).

It also should be mentioned, however, that other studies reported an increase of H2B monoubiquitination levels in response to DNA -damage (6-10). We could not observe such an effect in our experiments. It is possible that such discrepancies might be due to differential doses of DNA damaging agents used, different kinetic windows chosen or regions of chromatin affected. It could simply be that we have missed such an increase in H2Bub levels with our experimental settings.

Our identification of the GSE-HDAC1-USP22 multi-eraser complex and its function in DNA damage might be of relevance for cancer research. Various studies have recently connected GSE1 function to tumour growth in distinct types of cancer and highlighted its potential oncogenic and tumour suppressive functions (92-98). Overexpression of GSE1 was shown to promote proliferation of human breast cancer cells (93) whereas its downregulation positively correlated with decreased tumour invasion and metastasis in lung (94) or gastric cancer (98). Moreover, mutations in GSE1 were linked to a specific subtype of medulloblastomas accelerating their growth (95). In our study, we show that GSE1 is essential to integrate different enzymatic activities. It is, however, still an open question how these activities are regulated in response to DNA damage. We can rule out changes in deacetylase activity of HDAC1. We do not know how and whether the activity of USP22 is regulated, or whether the complex becomes localised to specific sites of damaged chromatin. However, we also dismiss DNA-damage induced changes in complex composition. Future work will certainly focus on these aspects. The fact that inhibitors of separate enzyme activities combined in the GSE1-HDAC1-USP22 complex are currently applied and tested in the clinics for cancer patients is potentially putting our findings into the context of clinical relevance (99,100).

## Supporting information

Supplementary Table 1. Interactome of FLAG-tagged HDAC1 in HAP1 cells

Supplementary Table 2. Interactome of FLAG-tagged HDAC1 in MEF cells

Supplementary Table 3. Acetylome data

Supplementary Table 4. Phosphorylome data

Supplementary Table 5. Interactome of V5-tagged GSE1 in HAP1 cells

Scripts used for MS shotgun data analysis

## Data Availability

The mass spectrometry proteomics data have been deposited to the ProteomeXchange Consortium (http://proteomecentral.proteomexchange.org) via the PRIDE partner repository (101) with the dataset identifier PXD040307 (interactomes) and PXD040339 (MS shotgun).

## Acknowledgements

We thank N. Hartl for the help with MS sample preparations, G. Brosch for the generous gift of acetylated histones, W. Chen for the support with AP-MS measurements and VBCF for providing the LC–MS instrument pool.

## CRediT author statement

Terezia Vcelkova: Conceptualization, Investigation, Methodology, Validation, Visualization, Writing – original draft. Wolfgang Reiter: Data curation, Formal analysis, Methodology, Visualisation, Writing – original draft. David Hollenstein: Software, Data curation, Formal analysis, Visualisation, Writing – original draft. Martha Zylka: Investigation, Validation. Stefan Schuckert: Investigation, Validation. Markus Hartl: Data curation, Formal analysis, Investigation, Methodology, Software, Validation, Writing – Review & Editing. Christian Seiser: Conceptualization, Funding acquisition, Project administration, Supervision, Writing – Review & Editing.

## Supplementary Figure legends

**Supplementary Figure 1.**
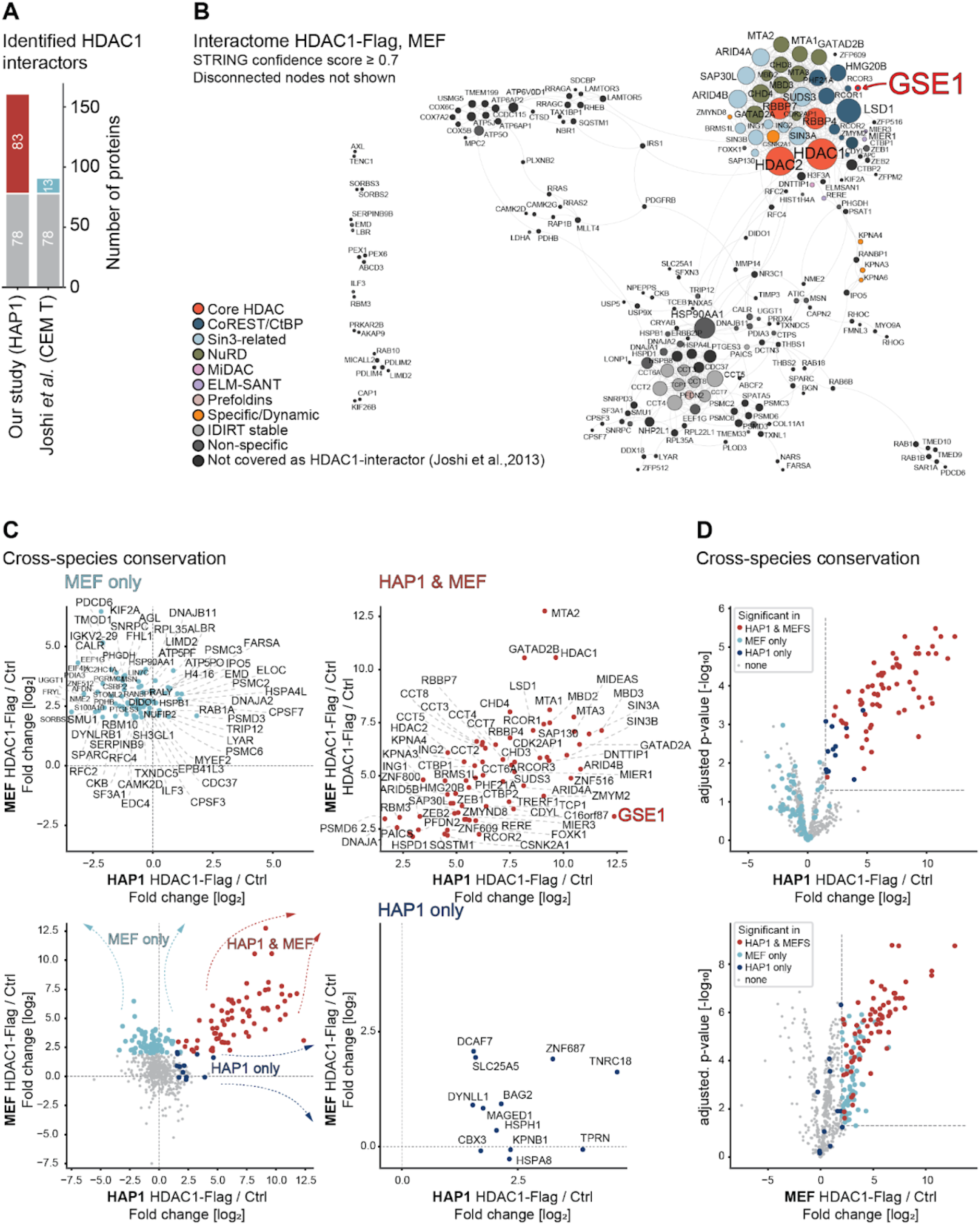
**(A)** Bar plot displaying number of proteins interacting with HDAC1 identified by Joshi *et al.* and our study. The red and blue boxes represent interactors found only in our study or by *Joshi et al. (41).* **(B)** STRING DB-based protein interaction network of the HDAC1 interactome in mouse embryonic fibroblasts (MEF). High-confidence interactions (confidence score ≥ 0.7) were selected. Disconnected nodes are not shown. The interactors were classified into functional protein groups in accordance with Joshi *et al.* (41). Strength of the associations are represented by the thickness of the edges. **(C)** Scatter plots illustrating cross-species conservation of the HDAC1 interactome determined by AP-MS in MEF and HAP1 cells. Proteins designated as interactors (as described in panel D) only in MEF cells are shown in light blue, only in HAP1 cells are shown in dark blue, and in both cell types are shown in red. **(D)** Volcano plot displaying the interactome of FLAG-tagged HDAC1 in HAP1 cells (top) and MEF cells (bottom) determined by AP-MS. Proteins designated as interactors only in MEF cells (≥ 2 log2-fold change [FC], adjusted p-value ≤ 0.05) are shown in light blue, only in HAP1 cells (≥ 1.5 log2-fold change [FC], adjusted p-value ≤ 0.05) are shown in dark blue, and in both cell types are shown in red.

**Supplementary Figure 2.**
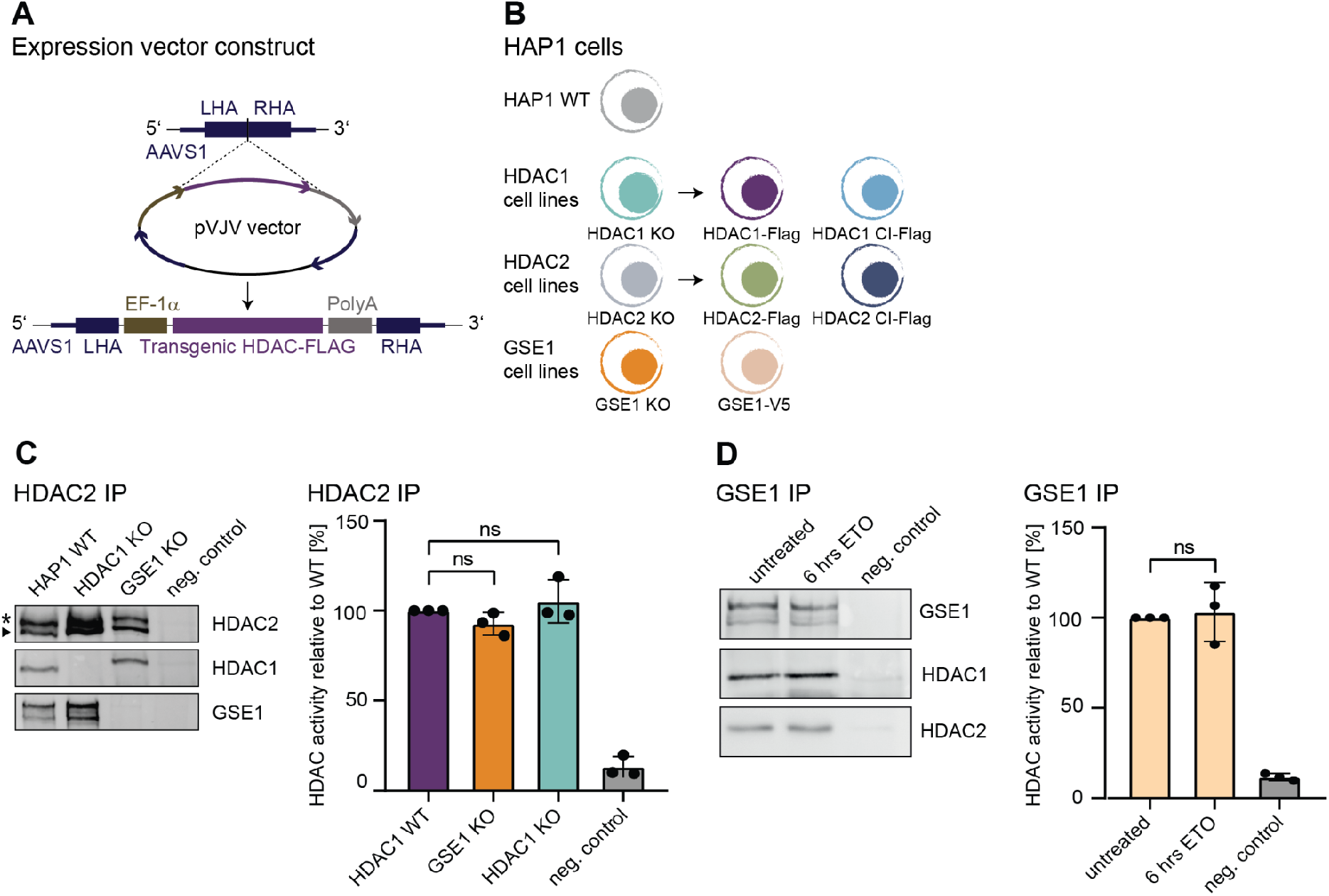
**(A)** Illustration of targeting strategy for establishment of transgenic cell lines expressing wildtype or inactive flag-tagged versions of HDAC1 or HDAC2. The transgenes (purple) were cloned into pVJV vector and integrated into safe harbour AAVS1 locus within HAP1 genome (dark blue) by CRISPR/Cas9 technology (LHA – left homology arm, RHA – right homology arm). The expression of transgenes was under the control of EF-1a promoter (army green). **(B)** Schematic illustration of cell lines used in this study. Original HAP1 WT cell line was used to generate HDAC1, HDAC2 and GSE1 knockout cell lines as well as the V5-tagged GSE1 cell line. Transgenic cell lines expressing wildtype and inactive versions of HDAC1 or HDAC2 were generated in their respective knockout backgrounds. **(C)** Western blot illustrating immunoprecipitation of endogenous HDAC2 complexes in HAP1 WT, GSE1 KO or HDAC1 KO cells. Band corresponding to HDAC2 is marked by an arrow. Asterisk: unspecific band. Antibodies used for detection are depicted on the right. Negative control: pull-downs performed using anti-mouse IgG antibodies. Impact of GSE1 ablation on the deacetylase activity of GSE1-HDAC2-containing complexes is shown in the bar graph. The data represent the mean values ± standard deviation (SD) of 3 biological replicates. Deacetylation activity obtained with HAP1 WT extracts was set to 100%. Significance was determined by unpaired parametric t-test by comparison to wildtype. *p < 0.05, **p < 0.01, ***p < 0.001. **(D)** Western blot illustrating immunoprecipitated endogenous GSE1 complexes in HAP1 WT cells treated with etoposide for 6 hours or left untreated. Antibodies used for detection are depicted on the right. Negative control: pull-downs performed using anti-mouse IgG antibodies. Effect of etoposide treatment on deacetylase activity of GSE1-HDAC complexes in HAP1 WT cells is shown in the bar graph. The data represent the mean values ± standard deviation (SD) of 3 biological replicates. Deacetylation activity obtained with untreated HAP1 WT extracts was set to 100%. Significance was determined by unpaired parametric t-test by comparison to wildtype. *p < 0.05, **p < 0.01, ***p < 0.001.

**Supplementary Figure 3.**
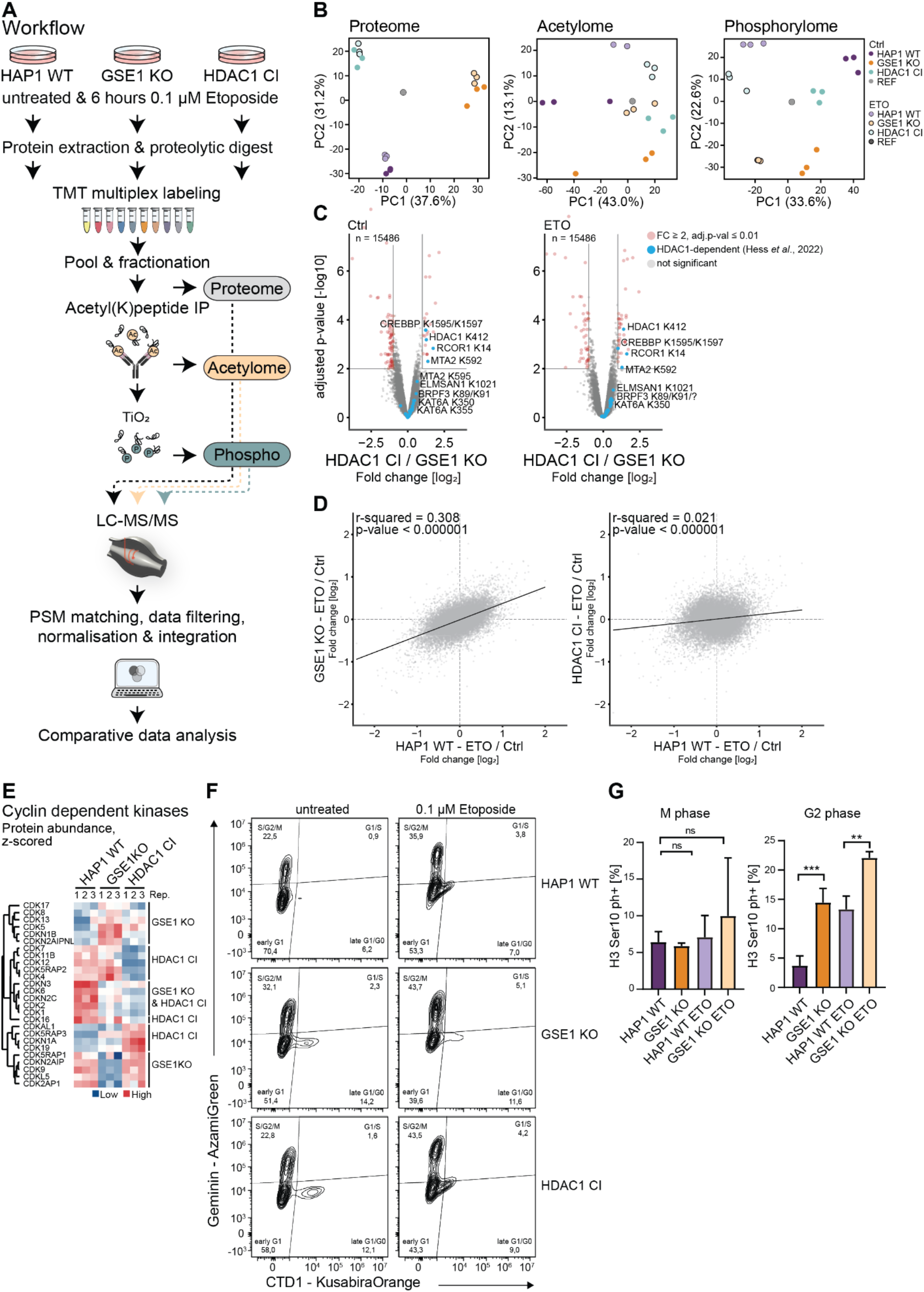
**(A)** Quantitative mass spectrometry workflow. Cells were treated for 6 hours with etoposide or left untreated, proteins were extracted and digested by trypsin. Peptides were labelled TMTpro 10plex, pooled and fractionated using high pH HPLC. Proteome aliquots were removed for MS measurement. Acetylated peptides were enriched by acetyl-lysine immunoprecipitation. Acetyl-IP flowthrough was used for enrichment of phosphorylated peptides by titanium dioxide precipitation. Finally, proteomes, acetylomes and phosphorylomes were analysed by LC-MS/MS. After data processing, the phosphorylation and acetylation sites were normalised to protein abundance and the effects of HDAC1 inactivation and GSE1 ablation were assessed in comparative analysis of untreated and etoposide treated conditions. **(B)** Principal Component Analysis (PCA) plots illustrating relatedness of proteome, acetylome and phosphorylome datasets. Acetylome profiles of untreated HAP1 WT and GSE1 KO samples showed higher variance between replicas. **(C)** Volcano plots displaying acetylome differences between HDAC1 CI and GSE1 KO cells, either in untreated conditions (left) or upon 6 hours treatment with 0.1 µM etoposide (right). Significantly different acetylation sites are shown in red (≥ 2-fold change, adjusted p-value ≤ 0.01), HDAC1-dependent acetylation sites previously described by Hess et al. are shown in blue. **(D)** Scatter plot comparing etoposide affected acetylation between HAP1 wildtype cells and GSE1 KO cells (right) or HDAC1 CI cells (left), related to Figure 4B. **(E)** Heatmap showing the protein abundance profiles of selected cyclin dependent kinases in untreated HAP1 WT, GSE1 KO, and HDAC1 CI cells. Corresponding gene names are indicated on the left. Values are z-scored. **(F)** Representative flow cytometry plots showing the cell cycle distribution of HAP1 WT, HDAC1 CI and GSE1 KO cells untreated or treated with etoposide for 24 hours. The data represent the percentage of cells in each cell cycle phase ± standard deviation (SD) of 3 biological replicates. Significance is determined by unpaired parametric t-test by comparison to wildtype. *p < 0.05, **p < 0.01, ***p < 0.001. **(G)** Immunofluorescence staining of H3 Ser10 phosphorylation quantified by counting of positive cells either in M-phase (robust signal in nucleus) and G2 phase (foci in the nucleus) are depicted. The data represent the percentage of cells in each M and G2 phase ± standard deviation (SD) of 3 biological replicates. On average 250 nuclei were counted per focal view. Significance is determined by unpaired parametric t-test by comparison to wildtype. *p < 0.05, **p < 0.01, ***p < 0.001.

**Supplementary Figure 4.**
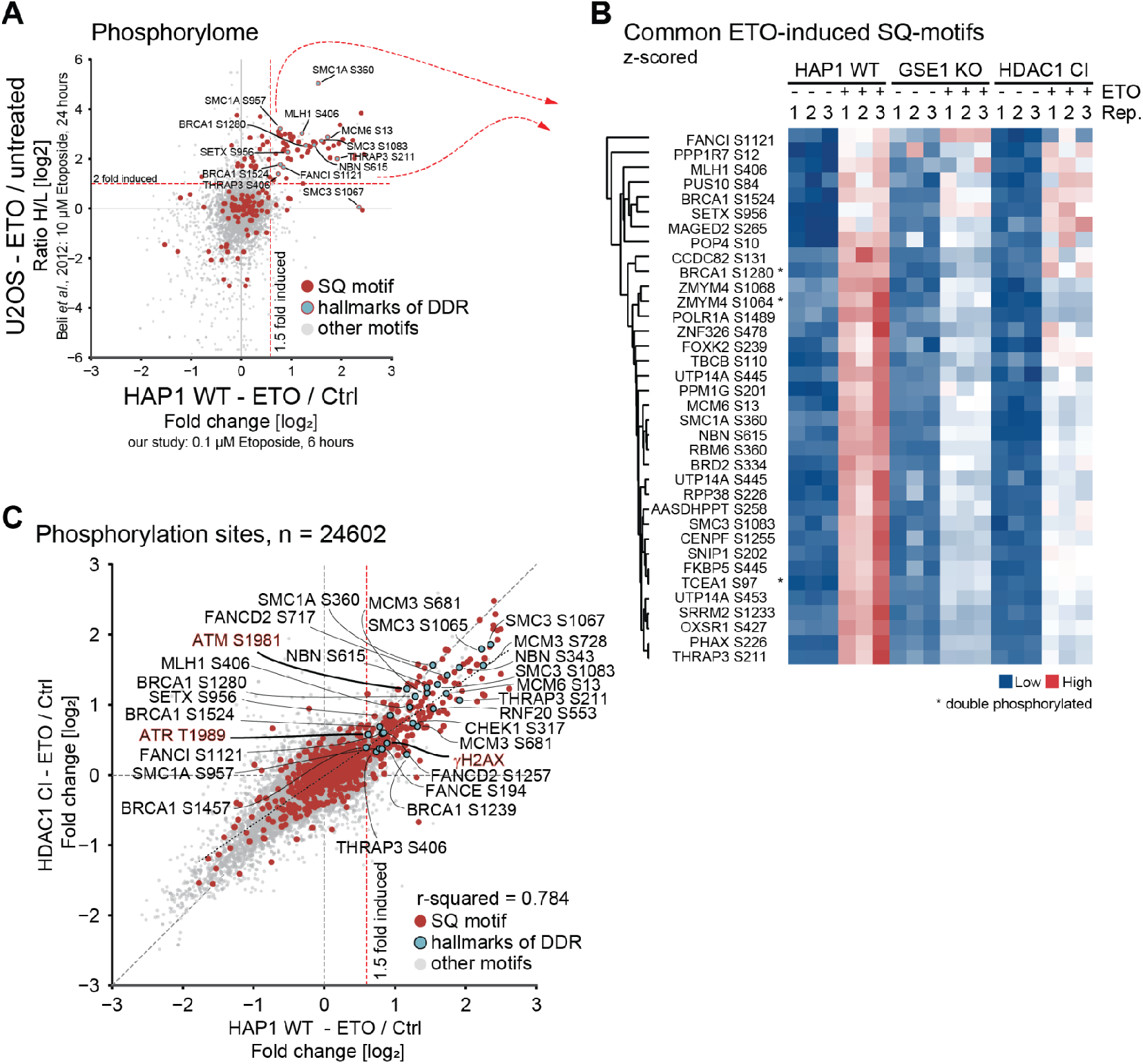
**(A)** Scatter plot of phosphorylation sites commonly quantified in our study and the study of Beli *et al.* (42), comparing the effects of long time (24 hours) and short time (6 hours) etoposide treatment. Sites harbouring an SQ motive are shown in red, sites representing hallmarks of the DDR are shown in blue. **(B)** Heatmap illustrating phosphorylation profiles in HAP1 WT, GSE1 KO and HDAC1 CI cells of SQ sites induced by 6 hours and 24 hours etoposide treatment (dataset of Beli et al. (42) cut-off ≥ 2-fold change, our study ≥ 1.5-fold change). * marks quantification of phosphorylation sites arising from double phosphorylated peptides where only a single site was confidently assigned. **(C)** Scatter plot comparing etoposide affected phosphorylation between HAP1 wildtype cells and HDAC1 CI cells, related to Figure 5A. Sites harbouring an SQ motive are shown in red, sites representing hallmarks of the DDR are shown in blue.

**Supplementary Figure 5.**
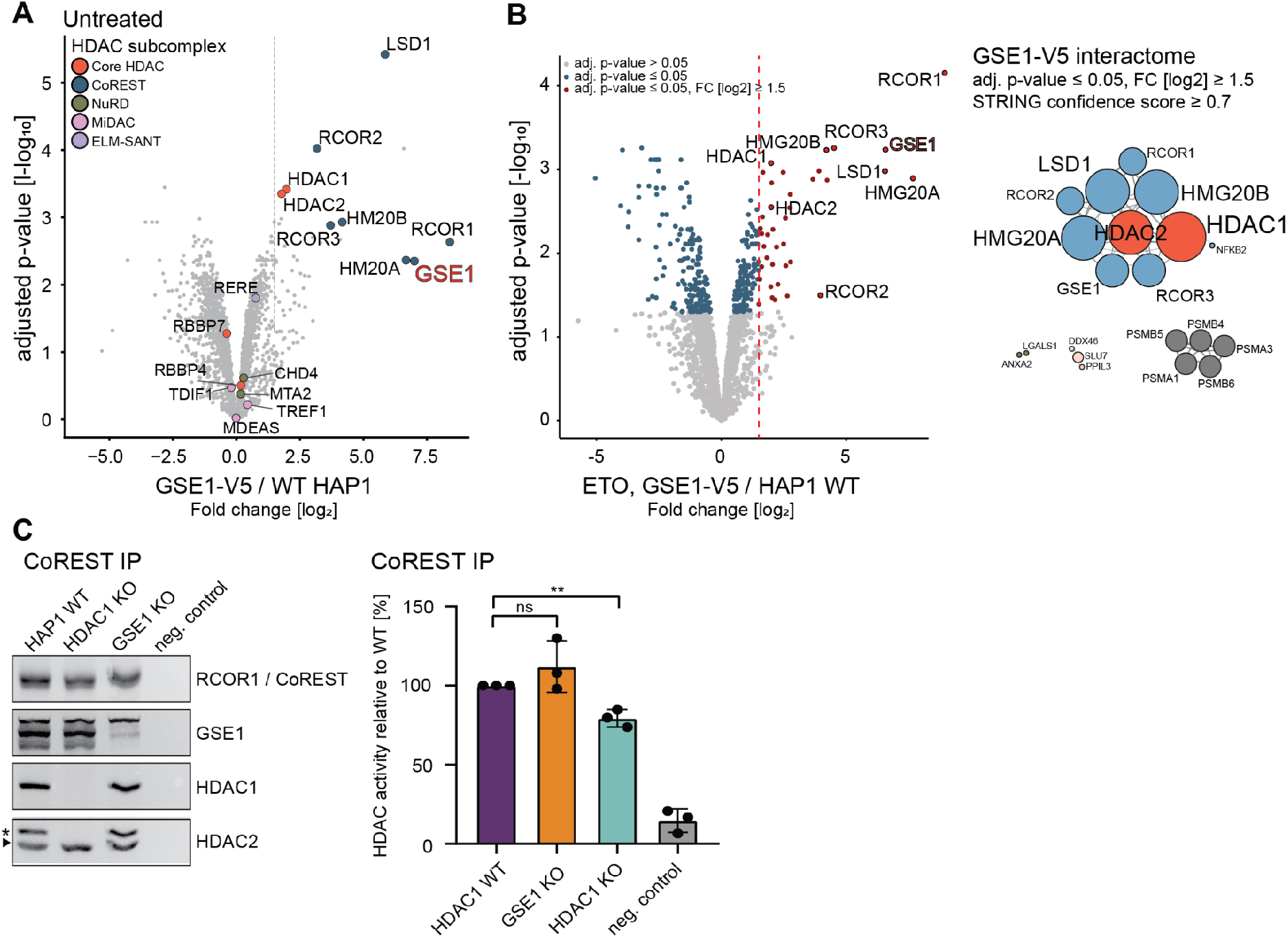
**(A)** Volcano plot displaying the interactome of GSE1-V5 in HAP1 cells determined by AP-MS. Gene names of known subunits of the HDAC1 corepressor complexes are highlighted, such as CoREST, SIN3-related or NuRD. GSE1 is highlighted with bold red text. **(B)** Left: Volcano plot displaying the interactome of GSE1-V5 in HAP1 cells treated with etoposide (0.1 µM, 6 hours) determined by AP-MS. Significant interactors (≥ 1.5-fold change [FC] over wildtype cells [HAP1 WT], adjusted p-value ≤ 0.05) in GSE1 V5-tagged cells are shown with red dots. Gene names of known subunits of the HDAC1/CoREST complex are highlighted, GSE1 is highlighted with bold red text. Right; protein interaction network of the GSE1 interactome in etoposide-treated cells (0.1 µM, 6 hours), based on STRING DB with a confidence cut-off score of ≥ 0.7. Disconnected nodes are not displayed. Nodes are coloured according to modularity classes, and the strength of the associations is represented by the thickness of the edges. **(C)** Western blot illustrating immunoprecipitated endogenous CoREST in HAP1 WT, GSE1 KO or HDAC1 KO cells. Antibodies used for detection are depicted on the left. Asterisk: unspecific band. Negative control: pull-downs performed using anti-rabbit IgA antibodies. Impact of GSE1 deletion on the deacetylase activity of GSE1-CoREST complexes is shown in the bar graph. The data represent the mean values ± standard deviation (SD) of 3 biological replicates. Deacetylation activity obtained with HAP1 WT extracts was set to 100%. Significance is determined by unpaired parametric t-test by comparison to wildtype. *p < 0.05, **p < 0.01, ***p < 0.001.

## Supplementary Table legends

**Supplementary Table 1. Interactome of FLAG-tagged HDAC1 in HAP1 cells**

List of proteins that have been co-purified in anti-FLAG AP/MS experiments. HDAC1-FLAG was purified from HAP1 cells (HDAC1-Flag). CTRL: HDAC1 KO cells. The table depicts MS/MS counts, log2 raw intensities and log2 normalised LFQ intensities. Missing LFQ values were imputed. The table also includes results of a differential expression (DE) analysis providing e.g., fold change (FC) and adjusted p-values for proteins enriched in the HDAC1-FLAG samples.

**Supplementary Table 2. Interactome of FLAG-tagged HDAC1 in MEF cells**

Summary of proteins that have been co-purified in anti-FLAG AP/MS experiments. HDAC1-FLAG was purified from MEFs (HDAC1-Flag). CTRL: untagged cells. The table depicts MS/MS counts, log2 raw intensities and log2 normalised LFQ intensities. Missing LFQ values were imputed. The table also includes results of a differential expression (DE) analysis providing e.g., fold change (FC) and adjusted p-values for proteins enriched in the HDAC1-FLAG samples.

**Supplementary Table 3. Acetylome data**

Summary of quantified acetylation sites over MS shotgun experiments of our study. The table includes log2 ratios of site to protein normalised TMT reporter ion intensities of quantified acetylation sites as well as results of the DE analysis. Conditions measured: untreated and treated with 0.1 µM etoposide for 6 hours. Backgrounds used: HAP1 WT, GSE1 KO, HDAC1 CI. Results from Beli et al., 2012 (42) are shown in column “Ratio H/L Etoposide [log2]”.

**Supplementary Table 4. Phosphorylome data**

Summary of quantified phosphorylation sites over MS shotgun experiments of our study. The table includes log2 ratios of site to protein normalised TMT reporter ion intensities of quantified phosphorylation sites as well as results of the DE analysis. Conditions measured: untreated and treated with 0.1 µM etoposide for 6 hours. Backgrounds used: HAP1 WT, GSE1 KO, HDAC1 CI. Results from Beli et al., 2012 (42) are shown in column “Ratio H/L Etoposide [log2]”.

**Supplementary Table 5. Interactome of V5-tagged GSE1 in HAP1 cells**

Summary of proteins that have been co-purified in anti-FLAG AP/MS experiments. GSE1-V5 was purified from HAP1 cells (HDAC1-Flag). CTRL: untagged HAP1 WT cells. Tested conditions: untreated and treated with 0.1µM etoposide for 6 hours.

## Graphical abstract

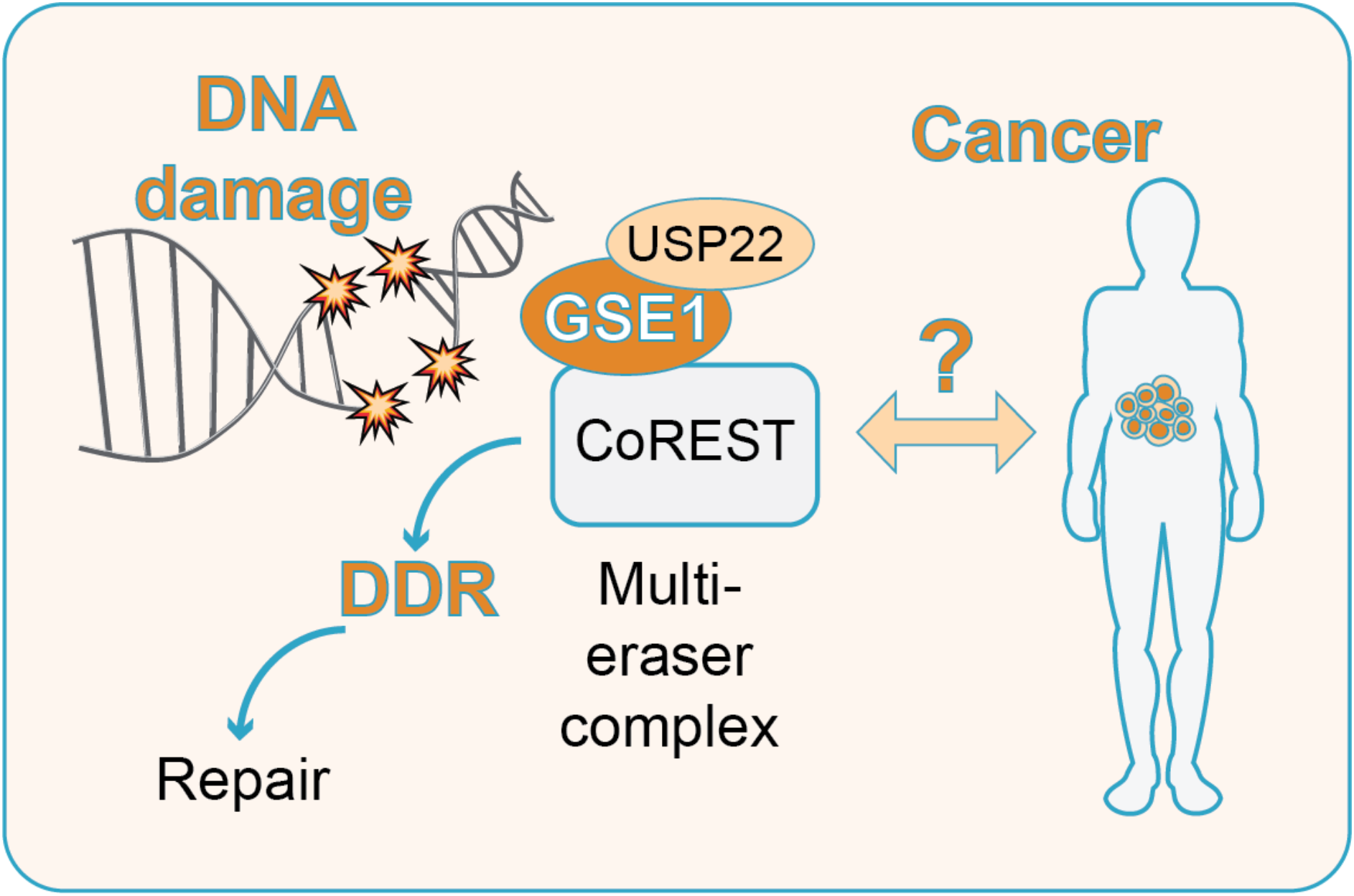

## References

1. Alhmoud, J.F., Woolley, J.F., Al Moustafa, A.E. and Malki, M.I. (2020) DNA Damage/Repair Management in Cancers. Cancers (Basel), 12.

2. Mendez-Acuna, L., Di Tomaso, M.V., Palitti, F. and Martinez-Lopez, W. (2010) Histone post-translational modifications in DNA damage response. Cytogenet Genome Res, 128, 28–36.

3. Paull, T.T., Rogakou, E.P., Yamazaki, V., Kirchgessner, C.U., Gellert, M. and Bonner, W.M. (2000) A critical role for histone H2AX in recruitment of repair factors to nuclear foci after DNA damage. Curr Biol, 10, 886–895.

4. Salguero, I., Belotserkovskaya, R., Coates, J., Sczaniecka-Clift, M., Demir, M., Jhujh, S., Wilson, M.D. and Jackson, S.P. (2019) MDC1 PST-repeat region promotes histone H2AX-independent chromatin association and DNA damage tolerance. Nat Commun, 10, 5191.

5. Ward, I.M., Minn, K., Jorda, K.G. and Chen, J. (2003) Accumulation of checkpoint protein 53BP1 at DNA breaks involves its binding to phosphorylated histone H2AX. J Biol Chem, 278, 19579–19582.

6. Doil, C., Mailand, N., Bekker-Jensen, S., Menard, P., Larsen, D.H., Pepperkok, R., Ellenberg, J., Panier, S., Durocher, D., Bartek, J. et al. (2009) RNF168 binds and amplifies ubiquitin conjugates on damaged chromosomes to allow accumulation of repair proteins. Cell, 136, 435–446.

7. Huyen, Y., Zgheib, O., Ditullio, R.A., Jr., Gorgoulis, V.G., Zacharatos, P., Petty, T.J., Sheston, E.A., Mellert, H.S., Stavridi, E.S. and Halazonetis, T.D. (2004) Methylated lysine 79 of histone H3 targets 53BP1 to DNA double-strand breaks. Nature, 432, 406–411.

8. Mailand, N., Bekker-Jensen, S., Faustrup, H., Melander, F., Bartek, J., Lukas, C. and Lukas, J. (2007) RNF8 ubiquitylates histones at DNA double-strand breaks and promotes assembly of repair proteins. Cell, 131, 887–900.

9. Moyal, L., Lerenthal, Y., Gana-Weisz, M., Mass, G., So, S., Wang, S.Y., Eppink, B., Chung, Y.M., Shalev, G., Shema, E. et al. (2011) Requirement of ATM-dependent monoubiquitylation of histone H2B for timely repair of DNA double-strand breaks. Mol Cell, 41, 529–542.

10. Nakamura, K., Kato, A., Kobayashi, J., Yanagihara, H., Sakamoto, S., Oliveira, D.V., Shimada, M., Tauchi, H., Suzuki, H., Tashiro, S. et al. (2011) Regulation of homologous recombination by RNF20-dependent H2B ubiquitination. Mol Cell, 41, 515–528.

11. Ramachandran, S., Haddad, D., Li, C., Le, M.X., Ling, A.K., So, C.C., Nepal, R.M., Gommerman, J.L., Yu, K., Ketela, T. et al. (2016) The SAGA Deubiquitination Module Promotes DNA Repair and Class Switch Recombination through ATM and DNAPK-Mediated gammaH2AX Formation. Cell Rep, 15, 1554–1565.

12. Mosammaparast, N., Kim, H., Laurent, B., Zhao, Y., Lim, H.J., Majid, M.C., Dango, S., Luo, Y., Hempel, K., Sowa, M.E. et al. (2013) The histone demethylase LSD1/KDM1A promotes the DNA damage response. J Cell Biol, 203, 457–470.

13. Sharma, G.G., So, S., Gupta, A., Kumar, R., Cayrou, C., Avvakumov, N., Bhadra, U., Pandita, R.K., Porteus, M.H., Chen, D.J. et al. (2010) MOF and histone H4 acetylation at lysine 16 are critical for DNA damage response and double-strand break repair. Mol Cell Biol, 30, 3582–3595.

14. Tjeertes, J.V., Miller, K.M. and Jackson, S.P. (2009) Screen for DNA-damage-responsive histone modifications identifies H3K9Ac and H3K56Ac in human cells. EMBO J, 28, 1878–1889.

15. Blattmann, C., Oertel, S., Ehemann, V., Thiemann, M., Huber, P.E., Bischof, M., Witt, O., Deubzer, H.E., Kulozik, A.E., Debus, J. et al. (2010) Enhancement of radiation response in osteosarcoma and rhabdomyosarcoma cell lines by histone deacetylase inhibition. Int J Radiat Oncol Biol Phys, 78, 237–245.

16. Camphausen, K., Burgan, W., Cerra, M., Oswald, K.A., Trepel, J.B., Lee, M.J. and Tofilon, P.J. (2004) Enhanced radiation-induced cell killing and prolongation of gammaH2AX foci expression by the histone deacetylase inhibitor MS-275. Cancer Res, 64, 316–321.

17. Krumm, A., Barckhausen, C., Kucuk, P., Tomaszowski, K.H., Loquai, C., Fahrer, J., Kramer, O.H., Kaina, B. and Roos, W.P. (2016) Enhanced Histone Deacetylase Activity in Malignant Melanoma Provokes RAD51 and FANCD2-Triggered Drug Resistance. Cancer Res, 76, 3067–3077.

18. Munshi, A., Kurland, J.F., Nishikawa, T., Tanaka, T., Hobbs, M.L., Tucker, S.L., Ismail, S., Stevens, C. and Meyn, R.E. (2005) Histone deacetylase inhibitors radiosensitize human melanoma cells by suppressing DNA repair activity. Clin Cancer Res, 11, 4912–4922.

19. Zhang, F., Zhang, T., Teng, Z.H., Zhang, R., Wang, J.B. and Mei, Q.B. (2009) Sensitization to gamma-irradiation-induced cell cycle arrest and apoptosis by the histone deacetylase inhibitor trichostatin A in non-small cell lung cancer (NSCLC) cells. Cancer Biol Ther, 8, 823–831.

20. Fradet-Turcotte, A., Canny, M.D., Escribano-Diaz, C., Orthwein, A., Leung, C.C., Huang, H., Landry, M.C., Kitevski-LeBlanc, J., Noordermeer, S.M., Sicheri, F. et al. (2013) 53BP1 is a reader of the DNA-damage-induced H2A Lys 15 ubiquitin mark. Nature, 499, 50–54.

21. Jacquet, K., Fradet-Turcotte, A., Avvakumov, N., Lambert, J.P., Roques, C., Pandita, R.K., Paquet, E., Herst, P., Gingras, A.C., Pandita, T.K. et al. (2016) The TIP60 Complex Regulates Bivalent Chromatin Recognition by 53BP1 through Direct H4K20me Binding and H2AK15 Acetylation. Mol Cell, 62, 409–421.

22. Pei, H., Zhang, L., Luo, K., Qin, Y., Chesi, M., Fei, F., Bergsagel, P.L., Wang, L., You, Z. and Lou, Z. (2011) MMSET regulates histone H4K20 methylation and 53BP1 accumulation at DNA damage sites. Nature, 470, 124–128.

23. Sharma, A.K., Bhattacharya, S., Khan, S.A., Khade, B. and Gupta, S. (2015) Dynamic alteration in H3 serine 10 phosphorylation is G1-phase specific during ionization radiation induced DNA damage response in human cells. Mutat Res, 773, 83–91.

24. Shimada, M., Niida, H., Zineldeen, D.H., Tagami, H., Tanaka, M., Saito, H. and Nakanishi, M. (2008) Chk1 is a histone H3 threonine 11 kinase that regulates DNA damage-induced transcriptional repression. Cell, 132, 221–232.

25. Tang, J., Cho, N.W., Cui, G., Manion, E.M., Shanbhag, N.M., Botuyan, M.V., Mer, G. and Greenberg, R.A. (2013) Acetylation limits 53BP1 association with damaged chromatin to promote homologous recombination. Nat Struct Mol Biol, 20, 317–325.

26. Guo, X., Bai, Y., Zhao, M., Zhou, M., Shen, Q., Yun, C.H., Zhang, H., Zhu, W.G. and Wang, J. (2018) Acetylation of 53BP1 dictates the DNA double strand break repair pathway. Nucleic Acids Res, 46, 689–703.

27. Jarrett, S.G., Carter, K.M., Bautista, R.M., He, D., Wang, C. and D’Orazio, J.A. (2018) Sirtuin 1-mediated deacetylation of XPA DNA repair protein enhances its interaction with ATR protein and promotes cAMP-induced DNA repair of UV damage. J Biol Chem, 293, 19025–19037.

28. Squatrito, M., Gorrini, C. and Amati, B. (2006) Tip60 in DNA damage response and growth control: many tricks in one HAT. Trends Cell Biol, 16, 433–442.

29. Tang, M., Li, Z., Zhang, C., Lu, X., Tu, B., Cao, Z., Li, Y., Chen, Y., Jiang, L., Wang, H. et al. (2019) SIRT7-mediated ATM deacetylation is essential for its deactivation and DNA damage repair. Sci Adv, 5, eaav1118.

30. Li, C., Irrazabal, T., So, C.C., Berru, M., Du, L., Lam, E., Ling, A.K., Gommerman, J.L., Pan-Hammarstrom, Q. and Martin, A. (2018) The H2B deubiquitinase Usp22 promotes antibody class switch recombination by facilitating non-homologous end joining. Nat Commun, 9, 1006.

31. Lin, D.W., Chung, B.P., Huang, J.W., Wang, X., Huang, L. and Kaiser, P. (2019) Microhomology-based CRISPR tagging tools for protein tracking, purification, and depletion. J Biol Chem, 294, 10877–10885.

32. Sakaue-Sawano, A., Kurokawa, H., Morimura, T., Hanyu, A., Hama, H., Osawa, H., Kashiwagi, S., Fukami, K., Miyata, T., Miyoshi, H. et al. (2008) Visualizing spatiotemporal dynamics of multicellular cell-cycle progression. Cell, 132, 487–498.

33. de la Barre, A.E., Gerson, V., Gout, S., Creaven, M., Allis, C.D. and Dimitrov, S. (2000) Core histone N-termini play an essential role in mitotic chromosome condensation. EMBO J, 19, 379–391.

34. Rappsilber, J., Mann, M. and Ishihama, Y. (2007) Protocol for micro-purification, enrichment, pre-fractionation and storage of peptides for proteomics using StageTips. Nat Protoc, 2, 1896–1906.

35. WeiqiangChen. (2022).

36. Hess, L., Moos, V., Lauber, A.A., Reiter, W., Schuster, M., Hartl, N., Lackner, D., Boenke, T., Koren, A., Guzzardo, P.M. et al. (2022) A toolbox for class I HDACs reveals isoform specific roles in gene regulation and protein acetylation. PLoS Genet, 18, e1010376.

37. Cox, J. and Mann, M. (2008) MaxQuant enables high peptide identification rates, individualized p.p.b.-range mass accuracies and proteome-wide protein quantification. Nat Biotechnol, 26, 1367–1372.

38. Plubell, D.L., Wilmarth, P.A., Zhao, Y., Fenton, A.M., Minnier, J., Reddy, A.P., Klimek, J., Yang, X., David, L.L. and Pamir, N. (2017) Extended Multiplexing of Tandem Mass Tags (TMT) Labeling Reveals Age and High Fat Diet Specific Proteome Changes in Mouse Epididymal Adipose Tissue. Mol Cell Proteomics, 16, 873–890.

39. Tyanova, S., Temu, T., Sinitcyn, P., Carlson, A., Hein, M.Y., Geiger, T., Mann, M. and Cox, J. (2016) The Perseus computational platform for comprehensive analysis of (prote)omics data. Nat Methods, 13, 731–740.

40. Nevers, Y., Kress, A., Defosset, A., Ripp, R., Linard, B., Thompson, J.D., Poch, O. and Lecompte, O. (2019) OrthoInspector 3.0: open portal for comparative genomics. Nucleic Acids Res, 47, D411–D418.

41. Joshi, P., Greco, T.M., Guise, A.J., Luo, Y., Yu, F., Nesvizhskii, A.I. and Cristea, I.M. (2013) The functional interactome landscape of the human histone deacetylase family. Mol Syst Biol, 9, 672.

42. Beli, P., Lukashchuk, N., Wagner, S.A., Weinert, B.T., Olsen, J.V., Baskcomb, L., Mann, M., Jackson, S.P. and Choudhary, C. (2012) Proteomic investigations reveal a role for RNA processing factor THRAP3 in the DNA damage response. Mol Cell, 46, 212–225.

43. Szklarczyk, D., Gable, A.L., Lyon, D., Junge, A., Wyder, S., Huerta-Cepas, J., Simonovic, M., Doncheva, N.T., Morris, J.H., Bork, P. et al. (2019) STRING v11: protein-protein association networks with increased coverage, supporting functional discovery in genome-wide experimental datasets. Nucleic Acids Res, 47, D607–D613.

44. Bastian, M., Heymann, S. and Jacomy, M. (2009) Gephi: An Open Source Software for Exploring and Manipulating Networks. Proceedings of the International AAAI Conference on Web and Social Media, 3, 361–362.

45. Jacomy, M., Venturini, T., Heymann, S. and Bastian, M. (2014) ForceAtlas2, a continuous graph layout algorithm for handy network visualization designed for the Gephi software. PLoS One, 9, e98679.

46. Andersson, B.S., Beran, M., Pathak, S., Goodacre, A., Barlogie, B. and McCredie, K.B. (1987) Ph-positive chronic myeloid leukemia with near-haploid conversion in vivo and establishment of a continuously growing cell line with similar cytogenetic pattern. Cancer Genet Cytogenet, 24, 335–343.

47. Nalawansha, D.A., Zhang, Y., Herath, K. and Pflum, M.K.H. (2018) HDAC1 Substrate Profiling Using Proteomics-Based Substrate Trapping. ACS Chem Biol, 13, 3315–3324.

48. Banks, C.A.S., Miah, S., Adams, M.K., Eubanks, C.G., Thornton, J.L., Florens, L. and Washburn, M.P. (2018) Differential HDAC1/2 network analysis reveals a role for prefoldin/CCT in HDAC1/2 complex assembly. Sci Rep, 8, 13712.

49. Barnes, C.E., English, D.M., Broderick, M., Collins, M.O. and Cowley, S.M. (2022) Proximity-dependent biotin identification (BioID) reveals a dynamic LSD1-CoREST interactome during embryonic stem cell differentiation. Mol Omics, 18, 31–44.

50. Hakimi, M.A., Bochar, D.A., Chenoweth, J., Lane, W.S., Mandel, G. and Shiekhattar, R. (2002) A core-BRAF35 complex containing histone deacetylase mediates repression of neuronal-specific genes. Proc Natl Acad Sci U S A, 99, 7420–7425.

51. Hakimi, M.A., Dong, Y., Lane, W.S., Speicher, D.W. and Shiekhattar, R. (2003) A candidate X-linked mental retardation gene is a component of a new family of histone deacetylase-containing complexes. J Biol Chem, 278, 7234–7239.

52. McClellan, D., Casey, M.J., Bareyan, D., Lucente, H., Ours, C., Velinder, M., Singer, J., Lone, M.D., Sun, W., Coria, Y. et al. (2019) Growth Factor Independence 1B-Mediated Transcriptional Repression and Lineage Allocation Require Lysine-Specific Demethylase 1-Dependent Recruitment of the BHC Complex. Mol Cell Biol, 39.

53. Kolle, D., Sarg, B., Lindner, H. and Loidl, P. (1998) Substrate and sequential site specificity of cytoplasmic histone acetyltransferases of maize and rat liver. FEBS Lett, 421, 109–114.

54. Gudkov, A.V., Kazarov, A.R., Thimmapaya, R., Axenovich, S.A., Mazo, I.A. and Roninson, I.B. (1994) Cloning mammalian genes by expression selection of genetic suppressor elements: association of kinesin with drug resistance and cell immortalization. Proc Natl Acad Sci U S A, 91, 3744–3748.

55. Montecucco, A., Zanetta, F. and Biamonti, G. (2015) Molecular mechanisms of etoposide. EXCLI J, 14, 95–108.

56. Mao, Z., Bozzella, M., Seluanov, A. and Gorbunova, V. (2008) DNA repair by nonhomologous end joining and homologous recombination during cell cycle in human cells. Cell Cycle, 7, 2902–2906.

57. Chao, O.S. and Goodman, O.B., Jr. (2014) Synergistic loss of prostate cancer cell viability by coinhibition of HDAC and PARP. Mol Cancer Res, 12, 1755–1766.

58. Maiso, P., Colado, E., Ocio, E.M., Garayoa, M., Martin, J., Atadja, P., Pandiella, A. and San-Miguel, J.F. (2009) The synergy of panobinostat plus doxorubicin in acute myeloid leukemia suggests a role for HDAC inhibitors in the control of DNA repair. Leukemia, 23, 2265–2274.

59. Min, K.Y., Lee, M.B., Hong, S.H., Lee, D., Jo, M.G., Lee, J.E., Choi, M.Y., You, J.S., Kim, Y.M., Park, Y.M. et al. (2021) Entinostat, a histone deacetylase inhibitor, increases the population of IL-10(+) regulatory B cells to suppress contact hypersensitivity. BMB Rep, 54, 534–539.

60. Bangert, A., Hacker, S., Cristofanon, S., Debatin, K.M. and Fulda, S. (2011) Chemosensitization of glioblastoma cells by the histone deacetylase inhibitor MS275. Anticancer Drugs, 22, 494–499.

61. Bayat Mokhtari, R., Baluch, N., Ka Hon Tsui, M., Kumar, S. T. S.H., Aitken, K., Das, B., Baruchel, S. and Yeger, H. (2017) Acetazolamide potentiates the anti-tumor potential of HDACi, MS-275, in neuroblastoma. BMC Cancer, 17, 156.

62. Fukuda, T., Wu, W., Okada, M., Maeda, I., Kojima, Y., Hayami, R., Miyoshi, Y., Tsugawa, K. and Ohta, T. (2015) Class I histone deacetylase inhibitors inhibit the retention of BRCA1 and 53BP1 at the site of DNA damage. Cancer Sci, 106, 1050–1056.

63. Goder, A., Mahendrarajah, N. and Kramer, O.H. (2017) Detection of Autophagy Induction After HDAC Inhibitor Treatment in Leukemic Cells. Methods Mol Biol, 1510, 3–10.

64. Subramanian, S., Bates, S.E., Wright, J.J., Espinoza-Delgado, I. and Piekarz, R.L. (2010) Clinical Toxicities of Histone Deacetylase Inhibitors. Pharmaceuticals (Basel), 3, 2751–2767.

65. Habibian, J. and Ferguson, B.S. (2018) The Crosstalk between Acetylation and Phosphorylation: Emerging New Roles for HDAC Inhibitors in the Heart. Int J Mol Sci, 20.

66. Li, J., Van Vranken, J.G., Pontano Vaites, L., Schweppe, D.K., Huttlin, E.L., Etienne, C., Nandhikonda, P., Viner, R., Robitaille, A.M., Thompson, A.H. et al. (2020) TMTpro reagents: a set of isobaric labeling mass tags enables simultaneous proteome-wide measurements across 16 samples. Nat Methods, 17, 399–404.

67. Elia, A.E., Boardman, A.P., Wang, D.C., Huttlin, E.L., Everley, R.A., Dephoure, N., Zhou, C., Koren, I., Gygi, S.P. and Elledge, S.J. (2015) Quantitative Proteomic Atlas of Ubiquitination and Acetylation in the DNA Damage Response. Mol Cell, 59, 867–881.

68. Bacevic, K., Lossaint, G., Achour, T.N., Georget, V., Fisher, D. and Dulic, V. (2017) Cdk2 strengthens the intra-S checkpoint and counteracts cell cycle exit induced by DNA damage. Sci Rep, 7, 13429.

69. Fung, T.K., Ma, H.T. and Poon, R.Y. (2007) Specialized roles of the two mitotic cyclins in somatic cells: cyclin A as an activator of M phase-promoting factor. Mol Biol Cell, 18, 1861–1873.

70. Huang, Y., Sramkoski, R.M. and Jacobberger, J.W. (2013) The kinetics of G2 and M transitions regulated by B cyclins. PLoS One, 8, e80861.

71. Sunada, S., Saito, H., Zhang, D., Xu, Z. and Miki, Y. (2021) CDK1 inhibitor controls G2/M phase transition and reverses DNA damage sensitivity. Biochem Biophys Res Commun, 550, 56–61.

72. Nam, C., Doi, K. and Nakayama, H. (2010) Etoposide induces G2/M arrest and apoptosis in neural progenitor cells via DNA damage and an ATM/p53-related pathway. Histol Histopathol, 25, 485–493.

73. Selvarajah, J., Elia, A., Carroll, V.A. and Moumen, A. (2015) DNA damage-induced S and G2/M cell cycle arrest requires mTORC2-dependent regulation of Chk1. Oncotarget, 6, 427–440.

74. Zhang, R., Zhu, L., Zhang, L., Xu, A., Li, Z., Xu, Y., He, P., Wu, M., Wei, F. and Wang, C. (2016) PTEN enhances G2/M arrest in etoposide-treated MCF-7 cells through activation of the ATM pathway. Oncol Rep, 35, 2707–2714.

75. Matsuoka, S., Ballif, B.A., Smogorzewska, A., McDonald, E.R., 3rd, Hurov, K.E., Luo, J., Bakalarski, C.E., Zhao, Z., Solimini, N., Lerenthal, Y. et al. (2007) ATM and ATR substrate analysis reveals extensive protein networks responsive to DNA damage. Science, 316, 1160-1166.

76. Clouaire, T., Rocher, V., Lashgari, A., Arnould, C., Aguirrebengoa, M., Biernacka, A., Skrzypczak, M., Aymard, F., Fongang, B., Dojer, N. et al. (2018) Comprehensive Mapping of Histone Modifications at DNA Double-Strand Breaks Deciphers Repair Pathway Chromatin Signatures. Mol Cell, 72, 250–262 e256.

77. Cook, P.J., Ju, B.G., Telese, F., Wang, X., Glass, C.K. and Rosenfeld, M.G. (2009) Tyrosine dephosphorylation of H2AX modulates apoptosis and survival decisions. Nature, 458, 591–596.

78. Schultz, D.C., Ayyanathan, K., Negorev, D., Maul, G.G. and Rauscher, F.J., 3rd. (2002) SETDB1: a novel KAP-1-associated histone H3, lysine 9-specific methyltransferase that contributes to HP1-mediated silencing of euchromatic genes by KRAB zinc-finger proteins. Genes Dev, 16, 919-932.

79. Tachibana, M., Sugimoto, K., Nozaki, M., Ueda, J., Ohta, T., Ohki, M., Fukuda, M., Takeda, N., Niida, H., Kato, H. et al. (2002) G9a histone methyltransferase plays a dominant role in euchromatic histone H3 lysine 9 methylation and is essential for early embryogenesis. Genes Dev, 16, 1779–1791.

80. Sims, J.J. and Cohen, R.E. (2009) Linkage-specific avidity defines the lysine 63-linked polyubiquitin-binding preference of rap80. Mol Cell, 33, 775–783.

81. Zhu, B., Zheng, Y., Pham, A.D., Mandal, S.S., Erdjument-Bromage, H., Tempst, P. and Reinberg, D. (2005) Monoubiquitination of human histone H2B: the factors involved and their roles in HOX gene regulation. Mol Cell, 20, 601–611.

82. Qureshi, I.A., Gokhan, S. and Mehler, M.F. (2010) REST and CoREST are transcriptional and epigenetic regulators of seminal neural fate decisions. Cell Cycle, 9, 4477–4486.

83. Song, Y., Dagil, L., Fairall, L., Robertson, N., Wu, M., Ragan, T.J., Savva, C.G., Saleh, A., Morone, N., Kunze, M.B.A. et al. (2020) Mechanism of Crosstalk between the LSD1 Demethylase and HDAC1 Deacetylase in the CoREST Complex. Cell Rep, 30, 2699–2711 e2698.

84. Cloud, A.S., Vargheese, A.M., Gunewardena, S., Shimak, R.M., Ganeshkumar, S., Kumaraswamy, E., Jensen, R.A. and Chennathukuzhi, V.M. (2022) Loss of REST in breast cancer promotes tumor progression through estrogen sensitization, MMP24 and CEMIP overexpression. BMC Cancer, 22, 180.

85. Ozdag, H., Teschendorff, A.E., Ahmed, A.A., Hyland, S.J., Blenkiron, C., Bobrow, L., Veerakumarasivam, A., Burtt, G., Subkhankulova, T., Arends, M.J. et al. (2006) Differential expression of selected histone modifier genes in human solid cancers. BMC Genomics, 7, 90.

86. Wu, J., Hu, L., Du, Y., Kong, F. and Pan, Y. (2015) Prognostic role of LSD1 in various cancers: evidence from a meta-analysis. Onco Targets Ther, 8, 2565–2570.

87. Schmidt, T., Samaras, P., Frejno, M., Gessulat, S., Barnert, M., Kienegger, H., Krcmar, H., Schlegl, J., Ehrlich, H.C., Aiche, S. et al. (2018) ProteomicsDB. Nucleic Acids Res, 46, D1271–D1281.

88. Rieckmann, J.C., Geiger, R., Hornburg, D., Wolf, T., Kveler, K., Jarrossay, D., Sallusto, F., Shen-Orr, S.S., Lanzavecchia, A., Mann, M. et al. (2017) Social network architecture of human immune cells unveiled by quantitative proteomics. Nat Immunol, 18, 583–593.

89. Howden, A.J.M., Hukelmann, J.L., Brenes, A., Spinelli, L., Sinclair, L.V., Lamond, A.I. and Cantrell, D.A. (2019) Quantitative analysis of T cell proteomes and environmental sensors during T cell differentiation. Nat Immunol, 20, 1542–1554.

90. Macinkovic, I., Theofel, I., Hundertmark, T., Kovac, K., Awe, S., Lenz, J., Forne, I., Lamp, B., Nist, A., Imhof, A. et al. (2019) Distinct CoREST complexes act in a cell-type-specific manner. Nucleic Acids Res, 47, 11649–11666.

91. Xiao, A., Li, H., Shechter, D., Ahn, S.H., Fabrizio, L.A., Erdjument-Bromage, H., Ishibe-Murakami, S., Wang, B., Tempst, P., Hofmann, K. et al. (2009) WSTF regulates the H2A.X DNA damage response via a novel tyrosine kinase activity. Nature, 457, 57–62.

92. Bamodu, O.A., Wang, Y.H., Ho, C.H., Hu, S.W., Lin, C.D., Tzou, K.Y., Wu, W.L., Chen, K.C. and Wu, C.C. (2021) Genetic Suppressor Element 1 (GSE1) Promotes the Oncogenic and Recurrent Phenotypes of Castration-Resistant Prostate Cancer by Targeting Tumor-Associated Calcium Signal Transducer 2 (TACSTD2). Cancers (Basel), 13.

93. Chai, P., Tian, J., Zhao, D., Zhang, H., Cui, J., Ding, K. and Liu, B. (2016) GSE1 negative regulation by miR-489-5p promotes breast cancer cell proliferation and invasion. Biochem Biophys Res Commun, 471, 123–128.

94. Ding, K., Tan, S., Huang, X., Wang, X., Li, X., Fan, R., Zhu, Y., Lobie, P.E., Wang, W. and Wu, Z. (2018) GSE1 predicts poor survival outcome in gastric cancer patients by SLC7A5 enhancement of tumor growth and metastasis. J Biol Chem, 293, 3949–3964.

95. Huang, M., Tailor, J., Zhen, Q., Gillmor, A.H., Miller, M.L., Weishaupt, H., Chen, J., Zheng, T., Nash, E.K., McHenry, L.K. et al. (2019) Engineering Genetic Predisposition in Human Neuroepithelial Stem Cells Recapitulates Medulloblastoma Tumorigenesis. Cell Stem Cell, 25, 433–446 e437.

96. Nicosia, L., Boffo, F.L., Ceccacci, E., Conforti, F., Pallavicini, I., Bedin, F., Ravasio, R., Massignani, E., Somervaille, T.C.P., Minucci, S. et al. (2022) Pharmacological inhibition of LSD1 triggers myeloid differentiation by targeting GSE1 oncogenic functions in AML. Oncogene, 41, 878–894.

97. Wang, W., Wang, S., Xu, A.M., Yuan, X., Huang, L. and Li, J. (2021) Overexpression of GSE1 Related to Trastuzumab Resistance in Gastric Cancer Cells. Biomed Res Int, 2021, 8834923.

98. Zhang, X., Tan, Z., Kang, T., Zhu, C. and Chen, S. (2019) Arsenic sulfide induces miR-4665-3p to inhibit gastric cancer cell invasion and migration. Drug Des Devel Ther, 13, 3037–3049.

99. Kalin, J.H., Wu, M., Gomez, A.V., Song, Y., Das, J., Hayward, D., Adejola, N., Wu, M., Panova, I., Chung, H.J. et al. (2018) Targeting the CoREST complex with dual histone deacetylase and demethylase inhibitors. Nat Commun, 9, 53.

100. Milelli, A., Marchetti, C., Turrini, E., Catanzaro, E., Mazzone, R., Tomaselli, D., Fimognari, C., Tumiatti, V. and Minarini, A. (2018) Novel polyamine-based Histone deacetylases-Lysine demethylase 1 dual binding inhibitors. Bioorg Med Chem Lett, 28, 1001–1004.

101. Perez-Riverol, Y., Bai, J., Bandla, C., Garcia-Seisdedos, D., Hewapathirana, S., Kamatchinathan, S., Kundu, D.J., Prakash, A., Frericks-Zipper, A., Eisenacher, M. et al. (2022) The PRIDE database resources in 2022: a hub for mass spectrometry-based proteomics evidences. Nucleic Acids Res, 50, D543–D552.

