## Supplementary material for "GSE1 links the HDAC1/CoREST co-repressor complex to DNA damage": Scripts used for MS shotgun data analysis: 01_proteingroups_data_processing_ver01.html

Quantitative proteomics analysis of HAP1 cells +/- etoposide treatment in different genetic backgrounds


### Quantitative proteomics analysis of HAP1 cells +/- etoposide treatment in different genetic backgrounds

##### submitted by Terezia Vcelkova (Seiser Lab)

###### performed by M. Hartl, Max Perutz Labs Mass Spectrometry Facility

###### Nov. 18, 2022

Note: This analysis was performed as an R Markdown Notebook in RStudio.
Not all code is displayed in the final report but is available as .rmd
file.

#### 1. Introduction

The experiments consists of 18 samples (3 genetic backgrounds: WT,
GSE1-KO, HDAC1-CI; 2 conditions: control, +etoposide; 3 biological
replicates), measured as two TMT-10plex sets. The tenth channel was a
pool of all 18 samples which was added to each set and used for
internal-reference-standard normalisation (IRS). The labeled peptides
were neutral-pH reversed phase fractionated (16 pooled for proteome, 8
phosphoproteome and acetylome), and enriched for acetylated peptides.
This results in a total of 64 measurements ([16 fraction proteome + 16
fractions PTM] x 2 sets). All runs were searched in MaxQuant and the
result files (proteinGroups.txt, phospho(STY)sites.txt,
acetyl(K)sites.txt) will be corrected for isotopic impurities during the
analysis. This analysis and Markdown script focuses on the proteome
data.

#### 2. Load data and quality control

The data were loaded and reverse database hits removed. As quality
control, we will create an overview of the data.

First, we compare the signal strength of all channels:

The distribution of channel intensities and of contaminants seems to
be largely comparable, as expected, and the intensity of contaminants is
rather low and uniform. Contaminants were thus removed for further
analysis steps, as well as proteins only identified by site (which
cannot be quantified correctly).

Protein groups identified and remaining after filtering:

| total | -reverse | -reverse & con | -rev & con & only by site |
| --- | --- | --- | --- |
| 11016 | 10356 | 10258 | 9793 |

It is known that multiple TMT-sets overlap only partially due to
stochasticiy in the MS/MS measurement. In this experiment we observe
about 93% overlap on protein group level between the two sets, which is
in the expected range:

#### 3.Correction for isotopic impurities

Isotopically labeled reagents used for TMT synthesis are not 100%
pure. We correct for these isotopic impurities using the data provided
by the manufacturer for the used lot of reagents, yielding corrected
data matrices for both sets.

#### 4. Normalisation procedures

To be able to compare the sample we need to apply two
normalisations:  
1. Global normalisation of all channels within one 10-plex set using
median scaling. 2. Normalisation between both 10-plex sets using
internal-reference-standard normalisation (IRS) according to Plubell et
al. (doi: 10.1074/mcp.M116.065524).

##### Median scaling

All channels of one set are scaled to the mean of all channel median
intensities. Then only the intersection (overlap) of all three
experiments is selected for IRS normalisation.

##### IRS normalisation

For IRS normalisation the geometric mean of all reference channel
intensities is calculated (per protein), and then all values for each
set are scaled to this common reference.

To inspect the effect of IRS-normalisation we perform PCA analysis
before and after the procedure and also show the effect of the
normalization in boxplots.

The PCA shows that IRS-normalization removed the batch effects and
that the replicates of each group cluster well in the PCA.

For further inspection we calculate Pearson correlation coefficients
and plot them in a heatmap. The samples and replicates are highly
correlated with all coefficients >0.95.

Results are stored as full matrix with corrected values:

```
write.table(df_prot_norm, file = "./output/proteinGroups_corrected.txt", sep = "\t", col.names = TRUE, row.names=FALSE)
```

#### 5. Group comparison and statistical analysis

For quantitative statistical analysis the normalised dataset was
further filtered for protein groups having at least 2 razor or unique
peptides and for all protein groups with an intensity >= the 0.5%
quantile of the mean top 3 intense channels. 8574 of a total of 8742,
113 protein groups remained after filtering.

The resulting matrix was then used to calculate LIMMA statistics to
determine statistically significant differential expression changes
between sample groups.

The following comparisons were made:  
Gse1\_KO\_ctrl-WT\_ctrl, HDAC1.CI\_ctrl-WT\_ctrl, Gse1\_KO\_ctrl-HDAC1.CI\_ctrl,
Gse1\_KO\_eto-WT\_eto, HDAC1.CI\_eto-WT\_eto, Gse1\_KO\_eto-HDAC1.CI\_eto,
Gse1\_KO\_eto-Gse1\_KO\_ctrl, HDAC1.CI\_eto-HDAC1.CI\_ctrl, WT\_eto-WT\_ctrl

Significantly regulated proteins (at 5% FDR) for all
comparisons:

|  | Gse1\_KO\_ctrl-WT\_ctrl | HDAC1.CI\_ctrl-WT\_ctrl | Gse1\_KO\_ctrl-HDAC1.CI\_ctrl | Gse1\_KO\_eto-WT\_eto | HDAC1.CI\_eto-WT\_eto | Gse1\_KO\_eto-HDAC1.CI\_eto | Gse1\_KO\_eto-Gse1\_KO\_ctrl | HDAC1.CI\_eto-HDAC1.CI\_ctrl | WT\_eto-WT\_ctrl |
| --- | --- | --- | --- | --- | --- | --- | --- | --- | --- |
| Down | 2776 | 3145 | 2752 | 2859 | 3352 | 2679 | 165 | 379 | 326 |
| NotSig | 2672 | 2549 | 2534 | 2511 | 2443 | 2347 | 8021 | 7856 | 7560 |
| Up | 3126 | 2880 | 3288 | 3204 | 2779 | 3548 | 388 | 339 | 688 |

As a graphical overview, we also present the results as volcano plots
with p-values (not corrected for multiple testing) or adj.p.values
(corrected). Red indicates p-values or adjusted p-values below 0.05,
blue above, respectively:

All data are finally stored in a matrix for further processing
(including Protein group IDs & gene names):

```
df_prot_limma <- data.frame(df_p, tt_exp_limma)
write.table(df_prot_limma, file = "./output/protein_groups_limma_results.txt", sep = "\t", row.names = FALSE, quote = FALSE)
```
