## Supplementary material for "GSE1 links the HDAC1/CoREST co-repressor complex to DNA damage": Scripts used for MS shotgun data analysis: 01_proteingroups_data_processing_ver01.Rmd

````
---
title: "Quantitative proteomics analysis of HAP1 cells +/- etoposide treatment in different genetic backgrounds"
author: "data analysis performed by M. Hartl, Max Perutz Labs Mass Spectrometry Facility"
date: "Nov. 18, 2022"
output:
  html_document:
    df_print: paged
subtitle: submitted by Terezia Vcelkova (Seiser Lab)
---

Note: This analysis was performed as an [R Markdown](http://rmarkdown.rstudio.com) Notebook in RStudio. Not all code is displayed in the final report but is available as .rmd file.

## 1. Introduction

The experiments consists of 18 samples (3 genetic backgrounds: WT, GSE1-KO, HDAC1-CI; 2 conditions: control, +etoposide; 3 biological replicates), measured as two TMT-10plex sets. The tenth channel was a pool of all 18 samples which was added to each set and used for internal-reference-standard normalisation (IRS). The labeled peptides were neutral-pH reversed phase fractionated (16 pooled for proteome, 8 phosphoproteome and acetylome), and enriched for acetylated peptides. This results in a total of 64 measurements ([16 fraction proteome + 16 fractions PTM] x 2 sets). All runs were searched in MaxQuant and the result files (proteinGroups.txt, phospho(STY)sites.txt, acetyl(K)sites.txt) will be corrected for isotopic impurities during the analysis.
This analysis and Markdown script focuses on the proteome data. 

```{r load libraries, fig.show='hide', message=FALSE, warning=FALSE, include=FALSE, paged.print=FALSE, results='hide'}
## first part of the script was written by Moritz Madern
library(ggVennDiagram)
library(ggplot2)
library(RColorBrewer)
library(MSnbase)
library(knitr)
library(limma)
library(qvalue)
library(scales)
library(reshape2)

display.brewer.all()
group_colors <- c(brewer.pal(12, "Set3"), brewer.pal(6, "Dark2")) 
```


```{r read data, fig.show='hide', message=FALSE, warning=FALSE, include=FALSE, paged.print=FALSE, results='hide'}
# read in prot data
df_prot <- read.delim("./input/proteinGroups.txt", header=TRUE, sep="\t", stringsAsFactors = FALSE)
names(df_prot)
n_all <- nrow(df_prot)

# remove reverse hits
df_prot <- df_prot[!df_prot$Reverse== "+",]
n_rev <- nrow(df_prot)
```

## 2. Load data and quality control

The data were loaded and reverse database hits removed. As quality control, we will create an overview of the data.

First, we compare the signal strength of all channels:

```{r overview stats, echo=FALSE, message=FALSE, warning=FALSE, paged.print=FALSE, results='hide'}
# check channel intensities
#par(mfrow=c(1,2))
df_plot_int <- df_prot[,grep(pattern = "corrected", colnames(df_prot))]
colnames_plot_int <- c(paste0(rep("eto", times = 10), c(1:10)), paste0(rep("ctrl", times = 10), c(1:10)))                
colnames(df_plot_int) <- colnames_plot_int

barplot(colSums(df_plot_int), 
        las = 3, 
        cex.main = 0.8,
        main = "Protein intensity sum per channel", 
        xlab = "channel",
        ylab = "Intensity")
boxplot(log2(df_plot_int), 
        las = 3,
        cex.main = 0.8,
        main = "Protein intensity sum per channel", 
        xlab = "channel",
        ylab = "Intensity [log2]")

# check contamination levels
barplot(colSums(df_plot_int[df_prot$Potential.contaminant == "+",]), 
        las = 3, 
        cex.main = 0.8,
        main = "Contaminant intensity sum per channel", 
        xlab = "channel",
        ylab = "Intensity")

# remove contaminants
df_prot <- df_prot[!df_prot$Potential.contaminant == "+",]
n_rev_con <- nrow(df_prot)
df_prot <- df_prot[!df_prot$Only.identified.by.site == "+",]
n_rev_con_only <- nrow(df_prot)

# extract reporter intensity matrices for each set in a separate matrix
ind_m1 <- which(grepl(names(df_prot), pattern="corrected[.][0-9]+[. ]ETO"))
m1 <- as.matrix(df_prot[,ind_m1])

ind_m2 <- which(grepl(names(df_prot), pattern="corrected[.][0-9]+[. ]untreated"))
m2 <- as.matrix(df_prot[,ind_m2])

```

The distribution of channel intensities and of contaminants seems to be largely comparable, as expected, and the intensity of contaminants is rather low and uniform. Contaminants were thus removed for further analysis steps, as well as proteins only identified by site (which cannot be quantified correctly).

Protein groups identified and remaining after filtering:

```{r table proteins, echo=FALSE, message=FALSE, warning=FALSE}
n_proteins <- data.frame(n_all, n_rev, n_rev_con, n_rev_con_only)
colnames(n_proteins) <- c("total", "-reverse", "-reverse & con", "-rev & con & only by site")
kable(n_proteins)
```

It is known that multiple TMT-sets overlap only partially due to stochasticiy in the MS/MS measurement. In this experiment we observe about 93% overlap on protein group level between the two sets, which is in the expected range:

```{r venn diagram, echo=FALSE, message=FALSE, warning=FALSE, paged.print=FALSE, results='hide'}
## EXTRACT INFO FOR VENN DIAGRAM; THEN PLOT OVERLAP BETWEEN PLEXES IN VENN DIAGRAM -------------------------------------------------------------------------------------------------

# for each set, extract protein IDs that are non-zero 
list_1 <- df_prot$Protein.IDs[!apply(m1==0, MARGIN = 1, FUN=all)]
list_2 <- df_prot$Protein.IDs[!apply(m2==0, MARGIN = 1, FUN=all)]
list_nonNA <- list(first_plex=list_1, second_plex=list_2)
sapply(list_nonNA, FUN=length)

# plot venn diagram
ggVennDiagram(list_nonNA, label="both") +
   scale_fill_gradient(low="#deebf7",high = "#3182bd")
```

## 3.Correction for isotopic impurities

Isotopically labeled reagents used for TMT synthesis are not 100% pure. We correct for these isotopic impurities using the data provided by the manufacturer for the used lot of reagents, yielding corrected data matrices for both sets.

```{r Isotopic impurity correction, echo = FALSE, results = FALSE}
## This part of the script was originally written by Moritz Madern and adapted by MH
## ISOTOPIC IMPURITY CORRECTION -----------------------------------------------------------------------------------------

# read in impurity matrix (necessary for impurity correction)
impurity_matrix <- as.matrix(read.csv(file="./input/impurity10.txt", sep=",", header=TRUE, row.names = 1))
#rowSums(impurity_matrix)
impurity_matrix
#t(impurity_matrix)  # transposed
inverted_transposed_impurity_matrix <- solve(t(impurity_matrix))

# note: impurity matrix itself needs to be manually created from information on product sheet


# CORRECT FOR ISOTOPIC IMPURITIES IN PLEX1 -------------------------------------------------------------------------------------------------------------------------

# note: this is done by solving a linear equation system for every row in your reporter intensity matrix.
# The equation is:                                               t(impurity_matrix) * corrected_reporter_int  =  observed_reporter_int,
# thus solving this for corrected_reporter_int results in:                            corrected_reporter_int  =  (t(impurity_matrix))^(-1) * observed_reporter_int

# check if the order of reporter ions corresponds to order in impurity matrix (***check if that is the case, it is a requirement!***)
data.frame(txt_file_order = colnames(m1), impurity_matrix_order= rownames(t(impurity_matrix)))

# make a copy of m1 called m1_corrected (this is where the corrected results are stored), then do the correction in a loop
m1_corrected <-  m1
#dim(m1_corrected)


for (i in 1:nrow(m1)){
  
  # extract row_i
  row_i <- m1[i,]
  row_i
  
  # calculate corrected intensities, replace any negative values with 0
  row_i_corrected <- inverted_transposed_impurity_matrix %*% row_i
  row_i_corrected <- pmax(row_i_corrected,0)
  
  # store in m1_corrected 
  m1_corrected[i,] <- row_i_corrected
  
}  

# compare colSums before and after
par(mfrow=c(1,2))
barplot(colSums(m1, na.rm=TRUE), main="Plex 1 before correction", 
        las=1, 
        xlab = "channel",
        ylab = "Intensity",
        names = colnames_plot_int[1:10])
barplot(colSums(m1_corrected, na.rm=TRUE), main="Plex 1 after correction", 
        las=1, 
        xlab = "channel",
        ylab = "Intensity",
        names = colnames_plot_int[1:10])

# replace 0 with NAs in the corrected reporter intensity matrix
m1_corrected[m1_corrected==0] <- NA

# give rownames
rownames(m1) <- df_prot$Protein.IDs
#length(df_prot$Protein.IDs)

## CORRECT FOR ISOTOPIC IMPURITIES IN PLEX2 -------------------------------------------------------------------------------------------------------------------------

# check if the order of reporter ions corresponds to order in impurity matrix (***check if that is the case, it is a requirement!***)
data.frame(txt_file_order = colnames(m2), impurity_matrix_order= rownames(t(impurity_matrix)))


# make a copy of m2 called m2_corrected (this is where the corrected results are stored), then do the correction in a loop
m2_corrected <-  m2
#dim(m2_corrected)


for (i in 1:nrow(m2)){
  
  # extract row_i
  row_i <- m2[i,]
  row_i
  
  # calculate corrected intensities, replace any negative values with 0
  row_i_corrected <- inverted_transposed_impurity_matrix %*% row_i
  row_i_corrected <- pmax(row_i_corrected,0)
  
  # store in 2_corrected 
  m2_corrected[i,] <- row_i_corrected
  
}  


# compare colSums before and after
par(mfrow=c(1,2))
barplot(colSums(m2, na.rm=TRUE), main="Plex 2 before correction", 
        las=1, 
        xlab = "channel",
        ylab = "Intensity",
        names = colnames_plot_int[11:20])
barplot(colSums(m2_corrected, na.rm=TRUE), main="Plex 2 after correction", 
        las=1, 
        xlab = "channel",
        ylab = "Intensity",
        names = colnames_plot_int[11:20])


# replace 0 with NAs in the corrected reporter intensity matrix
m2_corrected[m2_corrected==0] <- NA


# give rownames
rownames(m2) <- df_prot$Protein.IDs
```

## 4. Normalisation procedures

To be able to compare the sample we need to apply two normalisations:\
1. Global normalisation of all channels within one 10-plex set using median scaling. 2. Normalisation between both 10-plex sets using internal-reference-standard normalisation (IRS) according to Plubell et al. (doi: 10.1074/mcp.M116.065524).

### Median scaling

All channels of one set are scaled to the mean of all channel median intensities. Then only the intersection (overlap) of all three experiments is selected for IRS normalisation.

```{r median normalisation, echo = FALSE, results='hide'}

### FIRST NORMALIZE WITHIN EACH PLEX TO OBTAIN EQUAL MEDIANS ---------------------------------------------------------------------------------------

# calculate average median across both sets
plex1_avg_median <- mean(apply(m1_corrected, MARGIN = 2, FUN = median, na.rm=TRUE))
plex2_avg_median <- mean(apply(m2_corrected, MARGIN = 2, FUN = median, na.rm=TRUE))

grand_median_mean <- mean(c(plex1_avg_median, plex2_avg_median))

# do normalization
m1_c_n <- sweep(m1_corrected, STATS= apply(m1_corrected, MARGIN = 2, FUN = median, na.rm=TRUE), FUN="/", MARGIN = 2)*grand_median_mean
apply(m1_c_n, MARGIN = 2, FUN = median, na.rm=TRUE)
n_factor_m1 <- grand_median_mean / apply(m1_corrected, MARGIN = 2, FUN = median, na.rm=TRUE)

m2_c_n <- sweep(m2_corrected, STATS= apply(m2_corrected, MARGIN = 2, FUN = median, na.rm=TRUE), FUN="/", MARGIN = 2)*grand_median_mean
apply(m2_c_n, MARGIN = 2, FUN = median, na.rm=TRUE)
n_factor_m2 <- grand_median_mean / apply(m2_corrected, MARGIN = 2, FUN = median, na.rm=TRUE)

par(mfrow=c(1,2))
par(mar = c(1,4,2,2) )
boxplot(log2(m1_c_n), ylab = "intensity [log2]", main = "Plex 1 after normal.", names = colnames_plot_int[1:10])
boxplot(log2(m2_c_n), main = "Plex 2 after normal.", ylab = "intensity [log2]", names = colnames_plot_int[11:20])

## SELECT INTERSECTING PROTEINS ONLY; DELETE THE REST ---------------------------------------

# find the intersection of all plexes
table(table((df_prot$Protein.IDs)))   # check if some Protein.IDs are duplicates (should not be the case); do not use Gene.names here, they will probably have duplicates
intersect_all_plex <- intersect(list_nonNA[[1]], list_nonNA[[2]])
length(intersect_all_plex)

bool_keep <- df_prot$Protein.IDs %in% intersect_all_plex
sum(bool_keep)

# delete non-overlapping protein.IDs names
df_prot <- df_prot[bool_keep,]
dim(df_prot)
m1_c_n <- m1_c_n[bool_keep,]
m2_c_n <- m2_c_n[bool_keep,]
```

### IRS normalisation

For IRS normalisation the geometric mean of all reference channel intensities is calculated (per protein), and then all values for each set are scaled to this common reference.

```{r IRS normalisation, echo = FALSE, results='hide'}
## NOW NORMALIZE BETWEEN ALL 2 PLEXES VIA REFERENCE CHANNEL ---------------------------------------------------------------

# extract reference channels (in this experiment it was always channel 15)
plex1_standard <- m1_c_n[,10, drop=FALSE]
plex2_standard <- m2_c_n[,10, drop=FALSE]

# get geometric mean of averaged references (one per plex)
final_reference <- (plex1_standard * plex2_standard)^(1/2)
final_reference[1:5]
plex1_standard[1:5] 
plex2_standard[1:5]


# calculate scaling factors that scale each plex_standard to the final reference
scale_factors_plex1 <- final_reference/plex1_standard
scale_factors_plex2 <- final_reference/plex2_standard

# Test:
(plex1_standard*scale_factors_plex1)[1:10] 
(plex2_standard*scale_factors_plex2)[1:10]

# scale intensity matrix by multiplying scale_factors for every row (protein), for every plex
plex1_irs <- sweep(m1_c_n, FUN="*", STATS=scale_factors_plex1, MARGIN=1)
plex2_irs <- sweep(m2_c_n, FUN="*", STATS=scale_factors_plex2, MARGIN=1)

# Test:
# average reference dublicates in each plex
plex1_standard_after_irs <- rowMeans(plex1_irs[,10, drop=FALSE])
plex2_standard_after_irs <- rowMeans(plex2_irs[,10, drop=FALSE])

plex1_standard_after_irs[1:5]
plex2_standard_after_irs[1:5]

# create normalized dataframe df_prot_norm
df_prot_norm <- df_prot
df_prot_norm[,ind_m1] <- plex1_irs
df_prot_norm[,ind_m2] <- plex2_irs

```

To inspect the effect of IRS-normalisation we perform PCA analysis before and after the procedure and also show the effect of the normalization in boxplots.

```{r PCA after IRS, echo = FALSE, results='hide'}
## COMPARE PCA BEFORE NORMALIZATION VIA REFERENCE CHANNEL ---------------------------------------------------

# specify group vector:
groups  <- c("WT_eto", "Gse1_KO_eto", "WT_eto", "Gse1_KO_eto", "WT_eto", "Gse1_KO_eto", "HDAC1-CI_eto", "HDAC1-CI_eto", "HDAC1-CI_eto", "REF_eto", "WT_ctrl", "Gse1_KO_ctrl", "WT_ctrl", "Gse1_KO_ctrl", "WT_ctrl", "Gse1_KO_ctrl", "HDAC1-CI_ctrl", "HDAC1-CI_ctrl", "HDAC1-CI_ctrl", "REF_ctrl")

table(groups)
names(group_colors) <- unique(groups)
group_colors
groups
pca_colors <- group_colors[groups]
pca_colors

# extract intensities before normalization
m_all_before <- as.matrix(df_prot[,c(ind_m1,ind_m2)])
m_all_before[m_all_before<=100] <- NA   # replace 0 with NA
m_all_before <- log(m_all_before, base=2)
m_all_before[is.na(m_all_before)] <- 0

# berechne PCA
pca_res <- prcomp(t(m_all_before))
rot_mat <- pca_res$rotation
res_final <- as.matrix(scale(t(m_all_before), center=TRUE, scale=FALSE)) %*% rot_mat

# calculate % variance explained
eigenvectors <- pca_res$sdev^2
anteil_var_pca1 <- round(eigenvectors[1]/sum(eigenvectors),digits=3)
anteil_var_pca2 <- round(eigenvectors[2]/sum(eigenvectors),digits=3)

# prepare plot before IRS
# plot PCA before normalization
par(xpd=TRUE)
par(mfrow=c(1,1))
par(mar=c(3,3,2,12))
par(mgp=c(2,0.7,0))
plot(res_final[,"PC1"], res_final[,"PC2"], pch=rep(c(1,2,3),each=16), col=pca_colors, yaxt="n", xaxt="n", ylab="", xlab="", cex=2, lwd=5)
axis(side=2, cex.axis = 0.7)
axis(side=1, cex.axis = 0.7)
title(xlab=paste0("PC1  ","(",anteil_var_pca1*100,"%",")"), ylab=paste0("PC2  ","(",anteil_var_pca2*100,"%",")"), cex.lab=0.8, main="PCA before normalization between Plexes")
legend("right", legend=unique(groups), col=group_colors[unique(groups)], bty="n", inset=-0.35, pch=16, cex=1)

## COMPARE PCA AFTER NORMALIZATION VIA REFERENCE CHANNEL ---------------------------------------------------

# extract intensities after normalization
m_all_after <- as.matrix(df_prot_norm[,c(ind_m1,ind_m2)])
m_all_after[m_all_after<=100] <- NA   # replace 0 with NA
m_all_after <- log(m_all_after, base=2)
m_all_after[is.na(m_all_after)] <- 0


# berechne PCA
pca_res <- prcomp(t(m_all_after))
rot_mat <- pca_res$rotation
res_final <- as.matrix(scale(t(m_all_after), center=TRUE, scale=FALSE)) %*% rot_mat


# calculate % variance explained
eigenvectors <- pca_res$sdev^2
anteil_var_pca1 <- round(eigenvectors[1]/sum(eigenvectors),digits=3)
anteil_var_pca2 <- round(eigenvectors[2]/sum(eigenvectors),digits=3)

# plot PCA after normalization
par(xpd=TRUE)
par(mfrow=c(1,1))
par(mar=c(3,3,2,12))
par(mgp=c(2,0.7,0))

plot(res_final[,"PC1"], res_final[,"PC2"], col=pca_colors, pch=rep(c(1,2,3),each=16), yaxt="n", xaxt="n", ylab="", xlab="", cex=2, lwd=5)
axis(side=2, cex.axis = 0.7)
axis(side=1, cex.axis = 0.7)
title(xlab=paste0("PC1  ","(",anteil_var_pca1*100,"%",")"), ylab=paste0("PC2  ","(",anteil_var_pca2*100,"%",")"), cex.lab=0.8, main="PCA after normalization between Plexes")
legend("right", legend=unique(groups), col=group_colors[unique(groups)], bty="n", inset=-0.35, pch=16, cex=1)

# draw boxplots
boxplot(m_all_before, 
        main = "Intensity before median & IRS normalisation", 
        xlab = "channel",
        ylab = "intensity [log2]", 
        names = colnames_plot_int[1:20])
boxplot(m_all_after, 
        main = "Intensity after median & IRS normalisation", 
        xlab = "channel",
        ylab = "intensity [log2]", 
        names = colnames_plot_int[1:20])

## EXPORT DATAFRAME WITH CORRECTED INTENSITIES  --------------------------------------------------------------------------
```

The PCA shows that IRS-normalization removed the batch effects and that the replicates of each group cluster well in the PCA.

For further inspection we calculate Pearson correlation coefficients and plot them in a heatmap. The samples and replicates are highly correlated with all coefficients \>0.95. 

```{r correlation analysis after IRS, echo = FALSE, results='hide'}
#copy matrix
m_all_after_cor <- m_all_after

#make names vector

cornames <- paste0(groups, "_", c(1,1,2,2,3,3,1,2,3,1,1,1,2,2,3,3,1,2,3,1))

#add proper names
colnames(m_all_after_cor) <- cornames

#remove reference channels
m_all_after_cor <- m_all_after_cor[,-c(grep(cornames, pattern = "REF"))]
m_all_after_cor <- m_all_after_cor[, order(colnames(m_all_after_cor))]

#the following codes for constructing Pearson correlation heatmaps is based on code from http://www.sthda.com/english/wiki/ggplot2-quick-correlation-matrix-heatmap-r-software-and-data-visualization, distributed under the creative commons license CC BY-NC-SA 3.0 US

#make correlation matrix
cormat <- round(cor(m_all_after_cor, method = "pearson"), 3)

# Get lower triangle of the correlation matrix
  get_lower_tri<-function(cormat){
    cormat[upper.tri(cormat)] <- NA
    return(cormat)
  }
  # Get upper triangle of the correlation matrix
  get_upper_tri <- function(cormat){
    cormat[lower.tri(cormat)]<- NA
    return(cormat)
  }
  
upper_tri <- get_upper_tri(cormat)
upper_tri


melted_cormat <- melt(upper_tri, na.rm = TRUE)
# Heatmap

# Create a ggheatmap
ggheatmap <- ggplot(melted_cormat, aes(Var2, Var1, fill = value))+
 geom_tile(color = "white")+
 scale_fill_gradient(low = "white", high = "red", 
   limit = c(0.95,1), space = "Lab", 
    name="Pearson\nCorrelation") +
  theme_minimal()+ # minimal theme
 theme(axis.text.x = element_text(angle = 90, vjust = 0.4, hjust = 1))+
 coord_fixed()+
  ggtitle("Heatmap of Pearson correlation coefficient matrix")

# Print the heatmap
print(ggheatmap)

```

Results are stored as full matrix with corrected values:

```{r write corrected and normalised matrix}
write.table(df_prot_norm, file = "./output/proteinGroups_corrected.txt", sep = "\t", col.names = TRUE, row.names=FALSE)
```

## 5. Group comparison and statistical analysis

```{r load data for limma & filter, include = FALSE}
# define channel names
channel_names <- cornames
unique_groups <- unique(groups)

# reduce to intensity columns
colnames(df_prot_norm)
df_p <- df_prot_norm[,grep(pattern = "corrected", colnames(df_prot_norm))]
# rename channels
colnames(df_p) <- channel_names
# add descriptive and filter columns
df_p <- data.frame(df_prot_norm[,c("Protein.IDs", "Gene.names", "Razor...unique.peptides","id")], df_p)
dim(df_p)
# min 2 razor peptides
df_p <- df_p[df_p$Razor...unique.peptides >= 2,]
dim(df_p)
# check rows with missing values and remove them
table(rowSums(is.na(df_p[,5:24])))
df_p <- df_p[rowSums(is.na(df_p[,5:24])) == 0, ]
dim(df_p)

# calculate mean of top three intensities per row
v_top3_int_mean_p <- apply(df_p[,5:24], 1, function(x) mean(sort(x, decreasing = TRUE)[1:3]))
hist(log(v_top3_int_mean_p,10))
quantile(v_top3_int_mean_p, probs = c(0.005, 0.01, 0.05))
# apply cut-off at 0.5% quantile
int_cut_off <- quantile(v_top3_int_mean_p, probs = c(0.005))
df_p <- df_p[v_top3_int_mean_p >= int_cut_off,]
dim(df_p)

```

For quantitative statistical analysis the normalised dataset was further filtered for protein groups having at least 2 razor or unique peptides and for all protein groups with an intensity \>= the 0.5% quantile of the mean top 3 intense channels. `r dim(df_p)[1]` of a total of `r dim(df_prot)` protein groups remained after filtering.

The resulting matrix was then used to calculate LIMMA statistics to determine statistically significant differential expression changes between sample groups.

```{r LIMMA, echo=FALSE, results = 'hide', message = FALSE, warning = FALSE}
#copy and log transform data
d_limma <- log2(df_p[,5:24])
row.names(d_limma) <- df_p[,1]
#remove reference channels
d_limma <- d_limma[-c(10,20)]
head(d_limma)
# set up the design matrix (with experiments as batch effect)
group <- as.factor(make.names(groups[-c(10,20)]))
#batch <- as.factor(c(rep("eto", 9), rep("ctrl", 9))) #as treatment and batch are the same introducing an identical batch variable would not make sense. We rely on the batch normalisation instead and accept that technical bias might not be fully separated from treatment effects. This was clear before the experiment and the decisions was taken that the genotype comparison within a treatment is more important than the treatment comparison within the same genotype. Moreover, available literature data can be used to validate the treatment effects. 
design <- model.matrix(~ 0 + group)
colnames(design) <- sub(pattern = ("group"), replacement = "", colnames(design))
colnames(design)
design

# make the contrast
comparisons <- c("Gse1_KO_ctrl-WT_ctrl", "HDAC1.CI_ctrl-WT_ctrl", "Gse1_KO_ctrl-HDAC1.CI_ctrl", "Gse1_KO_eto-WT_eto", "HDAC1.CI_eto-WT_eto", "Gse1_KO_eto-HDAC1.CI_eto", "Gse1_KO_eto-Gse1_KO_ctrl", "HDAC1.CI_eto-HDAC1.CI_ctrl", "WT_eto-WT_ctrl")
contrast <- makeContrasts(contrasts = comparisons, levels = design)
contrast

# do the linear model fitting
fit <- lmFit(d_limma, design)

# get the fit for the contrast of interest
fit2 <- contrasts.fit(fit, contrast)

# do the empirical Bayes moderation of the test statistic (with trended variance)
fit2 <- eBayes(fit2, trend = TRUE)

# show how many proteins are differentially abundant for each contrast
summary(decideTests(fit2))

# store results
tt_exp_limma <- topTable(fit2, coef = 1, sort.by = "none", number = Inf)

for(i in 2:length(comparisons)) {
  tt_exp_limma <- data.frame(tt_exp_limma, topTable(fit2, coef = i, sort.by = "none", number = Inf))
  print(length(colnames(tt_exp_limma)))
}

colnames(tt_exp_limma) <- paste0(colnames(tt_exp_limma)[1:6],".", rep(comparisons, each = 6))

```

The following comparisons were made:\
`r comparisons`

Significantly regulated proteins (at 5% FDR) for all comparisons:\
`r kable(summary(decideTests(fit2)))`

As a graphical overview, we also present the results as volcano plots with p-values (not corrected for multiple testing) or adj.p.values (corrected). Red indicates p-values or adjusted p-values below 0.05, blue above, respectively:

```{r Volcano and histogramm LIMMA all, echo=FALSE}

par(mfrow = c(1,2))
par(mar = c(5,5,2,0.5))

i_logFC <- grep("logFC", colnames(tt_exp_limma))

for (i in i_logFC) { 

  # store data in column as x and ys
  x <- tt_exp_limma[,i]
  y_pval <- -log10(tt_exp_limma[, i+3])
  y_apval <- -log10(tt_exp_limma[, i+4])
  
  # Plot pval
  plot(x, y_pval,
       xlab = colnames(tt_exp_limma)[i],
       ylab = colnames(tt_exp_limma)[i+3],
       main = paste(sub("logFC.", "", colnames(tt_exp_limma)[i]), "p-value"),
       cex.main = 0.8,
       cex.lab = 0.8,
       pch = 16,
       col = alpha(c("blue","red")[as.factor(tt_exp_limma[, i+3] <= 0.05)], 0.3))
  # Plot adj. pval
  plot(x, y_apval,
       xlab = colnames(tt_exp_limma)[i],
       ylab = colnames(tt_exp_limma)[i+4],
       main = paste(sub("logFC.", "", colnames(tt_exp_limma)[i]), " adj. p-value"),
       cex.main = 0.8,
       cex.lab = 0.8,
       pch = 16,
       col = alpha(c("blue","red")[as.factor(tt_exp_limma[, i+4] <= 0.05)], 0.3))
}
```

All data are finally stored in a matrix for further processing (including Protein group IDs & gene names):

```{r}
df_prot_limma <- data.frame(df_p, tt_exp_limma)
write.table(df_prot_limma, file = "./output/protein_groups_limma_results.txt", sep = "\t", row.names = FALSE, quote = FALSE)
```
````
