## Supplementary material for "GSE1 links the HDAC1/CoREST co-repressor complex to DNA damage": Scripts used for MS shotgun data analysis: 02_acetylKsites_data_processing_ver01.html

Quantitative analysis of protein lysine acetylation sites in HAP1 cells +/- etoposide treatment in different genetic backgrounds


### Quantitative analysis of protein lysine acetylation sites in HAP1 cells +/- etoposide treatment in different genetic backgrounds

###### data analysis performed by M. Hartl, Max Perutz Labs Mass Spectrometry Facility

###### November 18, 2022

Note: This analysis was performed as an R Markdown Notebook in RStudio.
Not all code will be displayed in the final report but is available as
.rmd file. Parts of the code were kindly provided by Moritz Madern and
further adapted.

#### 1. Introduction

The experiments consists of 18 samples (3 genetic backgrounds: WT,
GSE1-KO, HDAC1-CI; 2 conditions: control, +etoposide; 3 biological
replicates), measured as two TMT-10plex sets. The tenth channel was a
pool of all 18 samples which was added to each set and used for
internal-reference-standard normalisation (IRS). The labeled peptides
were neutral-pH reversed phase fractionated (16 pooled for proteome, 8
phosphoproteome and acetylome), and enriched for acetylated peptides.
This results in a total of 64 measurements ([16 fraction proteome + 16
fractions PTM] x 2 sets). All runs were searched in MaxQuant and the
result files (proteinGroups.txt, phospho(STY)sites.txt,
acetyl(K)sites.txt) will be corrected for isotopic impurities during the
analysis. This analysis and Markdown script focuses on the acetylome
data.

#### 2. Load data and correction of isotopic impurities

The data were loaded and rearranged for further processing. This
includes splitting the quantitative values for singly, doubly, or
multiply acetylated sites in separate entries (rows). Then, we compare
the signal strength of all channels before and after impurity correction
(in the same manner as for the proteome data). The data look as
expected.

#### 3. Within-set median normalisation

Before normalisation entries with no quantitative information have to
be removed (many of them were generated in the previous step in which
quantitative information for different acetylation states was
split).Then the data are median normalized (channel-wise). Overall
acetylome levels are rather stable and median normalisation is suitable
(which was also inspected by scatter plots, not shown here):

##### Principal component analysis

As quality control we perform PCA. As expected, we see the same
TMT-labeling set dependent batch effect as on the proteome level, which
makes IRS normalisation necessary.

#### 4. Between-set IRS normalisation

For IRS normalisation the geometric mean of all reference channel
intensities is calculated (per protein), and then all values for each
set are scaled to this common reference. We again inspect the results
using PCA. Plotting PC1 vs. PC2 clearly shows that the TMT-set dependent
batch effect was removed by the normalisation.

In addition we inspect the Pearson correlation between samples after
normalisation. An overall higher variation at the level of lysine
acetylation as compared to the proteome is expected, due to the general
lower signal of lysine acetylated peptides and the additional
aggregation of peptides on protein level. The clustering of experimental
groups in the PCA and the overall rather high correlation indicate a
reproducible dataset that allows further inspection in differential
analysis.

#### 5. Site-to-protein normalisation

To be able to differentiate site-level from protein level-changes,
the site data are normalized to protein data. Before this step, the the
acetyl-K sites are further filtered to increase reliability and
robustness of the data. This means that addition to the 1% FDR cut-off
and 40 score cut-off already applied in MaxQuant, we apply a score
filter of >=75, and an intensity cut-off at the 5% quantile of the
mean of the three most abundant channel intensities per site.  
The normalised site-data are then again inspected by PCA.

#### 6. Differential abundance analysis using LIMMA

LIMMA is performed with the “trend” option and using the following
model: (~ 0 + group).

The following significantly regulated sites (at 5% FDR) were
determined:

|  | Gse1\_KO\_ctrl-WT\_ctrl | HDAC1.CI\_ctrl-WT\_ctrl | Gse1\_KO\_ctrl-HDAC1.CI\_ctrl | Gse1\_KO\_eto-WT\_eto | HDAC1.CI\_eto-WT\_eto | Gse1\_KO\_eto-HDAC1.CI\_eto | Gse1\_KO\_eto-Gse1\_KO\_ctrl | HDAC1.CI\_eto-HDAC1.CI\_ctrl | WT\_eto-WT\_ctrl |
| --- | --- | --- | --- | --- | --- | --- | --- | --- | --- |
| Down | 1639 | 3119 | 368 | 478 | 513 | 389 | 77 | 71 | 1258 |
| NotSig | 12797 | 7903 | 14445 | 14567 | 14220 | 14782 | 15364 | 15329 | 13231 |
| Up | 1050 | 4464 | 673 | 441 | 753 | 315 | 45 | 86 | 997 |

The results are stored to:
“acK-sites\_filtered\_proteinnormalized\_LIMMA.txt” as well as in a reduced
format as
“acK-sites\_filtered\_proteinnormalized\_LIMMA\_condensed.txt”.

##### Volcano plots

Plots for all comparisons of interest are generated and stored,
marking significant sites (at 5% FDR and > 2-fold or < 0.5-fold
difference).
