## Supplementary material for "GSE1 links the HDAC1/CoREST co-repressor complex to DNA damage": Scripts used for MS shotgun data analysis: 02_acetylKsites_data_processing_ver01.Rmd

````
---
title: "Quantitative analysis of protein lysine acetylation sites in HAP1 cells +/- etoposide treatment in different genetic backgrounds"
author: "data analysis performed by M. Hartl, Max Perutz Labs Mass Spectrometry Facility"
date: "November 18, 2022"
output: html_document
---

Note: This analysis was performed as an [R Markdown](http://rmarkdown.rstudio.com) Notebook in RStudio. Not all code will be displayed in the final report but is available as .rmd file. Parts of the code were kindly provided by Moritz Madern and further adapted.

## 1. Introduction

The experiments consists of 18 samples (3 genetic backgrounds: WT, GSE1-KO, HDAC1-CI; 2 conditions: control, +etoposide; 3 biological replicates), measured as two TMT-10plex sets. The tenth channel was a pool of all 18 samples which was added to each set and used for internal-reference-standard normalisation (IRS). The labeled peptides were neutral-pH reversed phase fractionated (16 pooled for proteome, 8 phosphoproteome and acetylome), and enriched for acetylated peptides. This results in a total of 64 measurements ([16 fraction proteome + 16 fractions PTM] x 2 sets). All runs were searched in MaxQuant and the result files (proteinGroups.txt, phospho(STY)sites.txt, acetyl(K)sites.txt) will be corrected for isotopic impurities during the analysis.
This analysis and Markdown script focuses on the acetylome data. 

```{r load packages, echo=FALSE, results='hide', warning=FALSE, message=FALSE}
library(ggVennDiagram)
library(ggplot2)
library(RColorBrewer)
library(MSnbase)
library(knitr)
library(limma)
library(dendextend)
library(plotly)
library(ggrepel)
library(reshape2)

```


```{r Parameters, echo=FALSE, results='hide'}

## filepath of ac site txt-file
filename_path = "./input/Acetyl (K)Sites.txt"  

## filepath of impurity matrix
impurity_path = "./input/impurity10.txt"  

## define groups vector
groups  <- c("WT_eto", "Gse1_KO_eto", "WT_eto", "Gse1_KO_eto", "WT_eto", "Gse1_KO_eto", "HDAC1.CI_eto", "HDAC1.CI_eto", "HDAC1.CI_eto", "REF_eto", "WT_ctrl", "Gse1_KO_ctrl", "WT_ctrl", "Gse1_KO_ctrl", "WT_ctrl", "Gse1_KO_ctrl", "HDAC1.CI_ctrl", "HDAC1.CI_ctrl", "HDAC1.CI_ctrl", "REF_ctrl")


## define sample vector

samplenames <- paste0(groups, "_", c(1,1,2,2,3,3,1,2,3,1,1,1,2,2,3,3,1,2,3,1))


## define max number of missing values per group
max_nr_missing = 0

## create colorfactor
colorfactor <- as.factor(groups)
colors <- colorRampPalette(brewer.pal(8, "Set1"))(length(unique(colorfactor)))
names(colors) <- levels(colorfactor)
  
```


```{r write required function, echo=FALSE, results='hide'}

### impurity correction function ###
impurity_correction <- function(impuritymatrix, df, reporterIon_columnnames) {
  
  ## read in impurity matrix to be used to correct for isotopic impurities
  writeLines("Impurity Matrix:")
  print(impuritymatrix)
  inverted_transposed_impuritymatrix <- solve(t(impuritymatrix))  # using linear algebra, this matrix can be used to calculate the corrected intensities
    
  ## manually calculate impurity correction:
  m_uncorrected <- as.matrix(df[,reporterIon_columnnames])
  m_uncorrected[is.na(m_uncorrected)] <- 0
  rownames(m_uncorrected) <- df$unique_id
  m_corrected <- m_uncorrected
    
  for (i in 1:nrow(m_uncorrected)) {
      
    ## extract row_i
    row_i <- m_uncorrected[i,]
      
    ## calculate corrected intensities, replace any negative values with 0
    row_i_corrected <- inverted_transposed_impuritymatrix %*% row_i
    row_i_corrected <- pmax(row_i_corrected,0)
      
    ## store in m_corrected 
    m_corrected[i,] <- row_i_corrected
  }
    
  ## finally, replace values that were originally < 1  with 0 ("i.e. NA"). Else they would mimic real observed intensities, which they are not!
  m_corrected[m_uncorrected < 1] <- 0
    
  ## plot before and after correction:
  par(mfrow=c(1,1))
  barplot(colSums(m_uncorrected, na.rm = TRUE), las=2, main="colsums before impurity correction")
  barplot(colSums(m_corrected,na.rm=TRUE), las=2, main="colsums after impurity correction")
    
  ## replace uncorrected intensities in df with corrected intensities:
  df[,reporterIon_columnnames] <- m_corrected
  
  return(df)
}

### PCA plot function ###
PCA_plot <- function(m, colorfactor){
  
  # replace NAs with 0
  m[is.na(m)] <- 0
  
  # compute PCA
  pca_res <- prcomp(t(m))
  rot_mat <- pca_res$rotation
  res_final <- as.matrix(scale(t(m), center=TRUE, scale=FALSE)) %*% rot_mat
  eigenvectors <- pca_res$sdev^2
  anteil_var_pca1 <- round(eigenvectors[1]/sum(eigenvectors),digits=3)
  anteil_var_pca2 <- round(eigenvectors[2]/sum(eigenvectors),digits=3)
  anteil_var_pca3 <- round(eigenvectors[3]/sum(eigenvectors),digits=3)
  anteil_var_pca4 <- round(eigenvectors[4]/sum(eigenvectors),digits=3)
  anteil_var_pca5 <- round(eigenvectors[5]/sum(eigenvectors),digits=3)
  anteil_var_pca6 <- round(eigenvectors[6]/sum(eigenvectors),digits=3)
  anteil_var_pca7 <- round(eigenvectors[7]/sum(eigenvectors),digits=3)
  
  ## create colorfactor
  colorfactor <- as.factor(colorfactor)
  colors <- colorRampPalette(brewer.pal(8, "Set1"))(length(unique(colorfactor)))
  names(colors) <- levels(colorfactor)
  
  #plot PCA
  par(xpd=TRUE)
  par(mfrow=c(1,1))
  par(mar=c(5,5,5,7.5))
  par(mgp=c(2,0.7,0))
  
  
  plot(res_final[,"PC1"], res_final[,"PC2"], col=colors[colorfactor], pch=16, cex=1.5,lwd=3, yaxt="n", xaxt="n", ylab="", xlab="")
  axis(side=2, cex.axis = 0.7)
  axis(side=1, cex.axis = 0.7)
  title(xlab=paste0("PC1  ","(",anteil_var_pca1*100,"%",")"), ylab=paste0("PC2  ","(",anteil_var_pca2*100,"%",")"), cex.lab=0.8)
  par(xpd=TRUE)
  legend("right", legend=names(colors), col=colors, bty="n", cex=0.75,pch=16, inset=-0.225)

  # #plot PCA 1vs3
  # par(xpd=TRUE)
  # par(mfrow=c(1,1))
  # par(mar=c(5,5,5,7.5))
  # par(mgp=c(2,0.7,0))
  # 
  # 
  # plot(res_final[,"PC1"], res_final[,"PC3"], col=colors[colorfactor], pch=16, cex=1.5,lwd=3, yaxt="n", xaxt="n", ylab="", xlab="")
  # axis(side=2, cex.axis = 0.7)
  # axis(side=1, cex.axis = 0.7)
  # title(xlab=paste0("PC1  ","(",anteil_var_pca1*100,"%",")"), ylab=paste0("PC3  ","(",anteil_var_pca3*100,"%",")"), cex.lab=0.8)
  # par(xpd=TRUE)
  # legend("right", legend=names(colors), col=colors, bty="n", cex=0.75,pch=16, inset=-0.225)  
  # 
  # #plot PCA 1vs4
  # par(xpd=TRUE)
  # par(mfrow=c(1,1))
  # par(mar=c(5,5,5,7.5))
  # par(mgp=c(2,0.7,0))
  # 
  # 
  # plot(res_final[,"PC1"], res_final[,"PC4"], col=colors[colorfactor], pch=16, cex=1.5,lwd=3, yaxt="n", xaxt="n", ylab="", xlab="")
  # axis(side=2, cex.axis = 0.7)
  # axis(side=1, cex.axis = 0.7)
  # title(xlab=paste0("PC1  ","(",anteil_var_pca1*100,"%",")"), ylab=paste0("PC4  ","(",anteil_var_pca4*100,"%",")"), cex.lab=0.8)
  # par(xpd=TRUE)
  # legend("right", legend=names(colors), col=colors, bty="n", cex=0.75,pch=16, inset=-0.225)    
      
  ## plot percent of variance explained per principle component
  v_percent_explained <- c(anteil_var_pca1, anteil_var_pca2, anteil_var_pca3, anteil_var_pca4, anteil_var_pca5, anteil_var_pca6, anteil_var_pca7)*100
  names(v_percent_explained) <- paste0("PC ",1:7)
  barplot(v_percent_explained, ylab="% variance explained", names.arg = names(v_percent_explained), las=2)
  
  
}


### heatmap plot ###
heatmap_plot <- function(m, colfactor, samplenames){
  
  
  ## create colorfactor
  colorfactor <- as.factor(colorfactor)
  colors <- rainbow(length(unique(colorfactor)))
  names(colors) <- levels(colorfactor)
  
  
  ## replaces NAs with 0
  m[is.na(m)] <- 0
  colnames(m) <- samplenames
  
  ## plot heatmap
  goldpalette <- colorRampPalette(colors = c("#6baed6","#deebf7","#fed976","#feb24c","#fd8d3c","#fc4e2a"))
  par(mfrow=c(1,1))
  heatmap(m, Rowv = FALSE, Colv=FALSE, margins=c(9,9), cexCol = 1,scale="none",labRow="",col =goldpalette(40), ColSideColors = colors[groups], cex.lab=2)

}

```

## 2. Load data and correction of isotopic impurities
  
The data were loaded and rearranged for further processing. This includes splitting the quantitative values for singly, doubly, or multiply acetylated sites in separate entries (rows). Then, we compare the signal strength of all channels before and after impurity correction (in the same manner as for the proteome data). The data look as expected. 

```{r Read in data and check column names, then filter out unnecessary duplicate columns, echo=FALSE, results='hide'}

## read in file
df <-  read.delim(filename_path, sep="\t", header=TRUE)
names(df)
dim(df)


## save metadata:
df_meta <- df[,!grepl(names(df), pattern="^Reporter[.]intensity")]


## save reporter ion intensity with column names matching pattern: "Reporter[.]intensity[.][0-9]*___[123]", for each ending separately
df_int_1 <- df[,grepl(names(df), pattern="Reporter.intensity.\\d*[.]\\w*___[1]") ]
df_int_2 <- df[,grepl(names(df), pattern="Reporter.intensity.\\d*[.]\\w*___[2]") ]
df_int_3 <- df[,grepl(names(df), pattern="Reporter.intensity.\\d*[.]\\w*___[3]") ]


## add reporter intensities (for each ending separately) to metadata to create 3 separate data_frames. Also add #site column
df_1 <- cbind(df_meta, df_int_1)
df_2 <- cbind(df_meta, df_int_2)
df_3 <- cbind(df_meta, df_int_3)
df_1$Number.Sites <- 1
df_2$Number.Sites <- 2
df_3$Number.Sites <- 3


## replace old reporter names with uniform name
bool_reporter <- grepl(names(df_1), pattern="Reporter[.]intensity")
new_name <- c(paste0("Reporter.intensity_eto_", 1:10), paste0("Reporter.intensity_ctrl_", 1:10))
names(df_1)[bool_reporter] <- new_name
names(df_2)[bool_reporter] <- new_name
names(df_3)[bool_reporter] <- new_name


## merge all three df into one
df_all <- rbind(df_1, df_2, df_3)
dim(df_all)
table(df_all$Number.Sites)

```


```{r Do impurity correction on reporter ion intensities, echo=FALSE, results='hide'}

## check names of df_all, and find reporter intensity columns
names_reporter <- grep(names(df_all), pattern="Reporter[.]intensity_", value=TRUE)
names_reporter

## check summary of reporter ions
summary(df_all[,names_reporter])

## do impurity correction
df_all_corr <- impurity_correction(df = df_all, 
                                   reporterIon_columnnames = names_reporter[1:10],
                                   impuritymatrix = as.matrix(read.csv(file=impurity_path, sep=",", header=TRUE, row.names = 1)))

df_all_corr <- impurity_correction(df = df_all, 
                                   reporterIon_columnnames = names_reporter[11:20],
                                   impuritymatrix = as.matrix(read.csv(file=impurity_path, sep=",", header=TRUE, row.names = 1)))

```

## 3. Within-set median normalisation

Before normalisation entries with no quantitative information have to be removed (many of them were generated in the previous step in which quantitative information for different acetylation states was split).Then the data are median normalized (channel-wise). Overall acetylome levels are rather stable and median normalisation is suitable (which was also inspected by scatter plots, not shown here):

```{r Filter based on missing values, echo=FALSE, results='hide', fig.keep='none'}

## dimensions before filtering
writeLines("dimensions before filtering")
dim(df_all_corr)

## extract reporter ion intensities
m_reporter <- as.matrix(df_all_corr[,names_reporter])
m_reporter[m_reporter <= 1] <- NA

## overview missing values
barplot(table(is.na(m_reporter)), main="number of missing values (is.na) before filtering")

## kick rows with too many missing values
bool_kick <- apply(is.na(m_reporter), MARGIN=1, FUN=sum) > max_nr_missing
df_all_corr_filtered <- df_all_corr[!bool_kick,]
writeLines("dimensions after filtering")
dim(df_all_corr_filtered)

## overview of missing values after filtering
m_reporter <- as.matrix(df_all_corr_filtered[,names_reporter])
m_reporter[m_reporter <= 1] <- NA
barplot(table(is.na(m_reporter)), main="number of missing values (is.na) after filtering")

```


```{r Do normalization, echo=FALSE, results='hide'}


## extract reporter intensities, replace intensities smaller 1 with NA
m1_corrected <- m_reporter[, 1:10]
m2_corrected <- m_reporter[, 11:20]

### FIRST NORMALIZE WITHIN EACH PLEX TO OBTAIN EQUAL MEDIANS ---------------------------------------------------------------------------------------

# calculate mean of medians for each channel for each plex
plex1_avg_median <- mean(apply(m1_corrected, MARGIN = 2, FUN = median, na.rm=TRUE))
plex2_avg_median <- mean(apply(m2_corrected, MARGIN = 2, FUN = median, na.rm=TRUE))

# calculate mean of mean of medians across all three plexes
grand_median_mean <- mean(c(plex1_avg_median, plex2_avg_median))

# do normalization by dividing each value by column median and multiplying by grand median mean
m1_c_n <- sweep(m1_corrected, STATS= apply(m1_corrected, MARGIN = 2, FUN = median, na.rm=TRUE), FUN="/", MARGIN = 2)*grand_median_mean
apply(m1_c_n, MARGIN = 2, FUN = median, na.rm=TRUE) #check: all medians should be equal now

m2_c_n <- sweep(m2_corrected, STATS= apply(m2_corrected, MARGIN = 2, FUN = median, na.rm=TRUE), FUN="/", MARGIN = 2)*grand_median_mean
apply(m2_c_n, MARGIN = 2, FUN = median, na.rm=TRUE)


#combine all three into one matrix
m_reporter_norm <- cbind(m1_c_n, m2_c_n)

## plot colSums after normalization
barplot(colSums(m_reporter_norm, na.rm=TRUE), main="total intensity after norm", las=2, col=colors[groups], names.arg = samplenames)
boxplot(log2(m_reporter_norm), main="intensity after norm", col=colors[groups])
#apply(m_reporter_norm, FUN=summary, MARGIN = 2)

## do pair plotting, only using 6 columns
#pairs(log2(m_reporter_norm[,c(1,2,7,11,12,17)]), cex=0.1)

## replace old intensities with normalized intensities
df_all_corr_filtered_norm <- df_all_corr_filtered
df_all_corr_filtered_norm[,names_reporter] <- m_reporter_norm

```

### Principal component analysis  
As quality control we perform PCA. As expected, we see the same TMT-labeling set dependent batch effect as on the proteome level, which makes IRS normalisation necessary.

```{r make PCA, echo=FALSE, results='hide'}

## extract reporter intensity matrix
m_reporter <- as.matrix(df_all_corr_filtered_norm[,names_reporter])
#barplot(colSums(m_reporter, na.rm=TRUE))


## plot PCA
PCA_plot(m=log2(m_reporter),
         colorfactor = groups)


```

## 4. Between-set IRS normalisation
  
For IRS normalisation the geometric mean of all reference channel intensities is calculated (per protein), and then all values for each set are scaled to this common reference. We again inspect the results using PCA. Plotting PC1 vs. PC2 clearly shows that the TMT-set dependent batch effect was removed by the normalisation. 
  
```{r IRS normalisation, echo = FALSE, results='hide'}
# extract reference channels (in this experiment it was always channel 10)
plex1_standard <- m1_c_n[,10, drop=FALSE]
plex2_standard <- m2_c_n[,10, drop=FALSE]

# calculate geometric mean of averaged references (one per plex)
final_reference <- (plex1_standard * plex2_standard)^(1/2)

# calculate scaling factors that scale each plex_standard to the final reference
scale_factors_plex1 <- final_reference/plex1_standard
scale_factors_plex2 <- final_reference/plex2_standard

# Test: now all values for the reference channels should be the same
(plex1_standard*scale_factors_plex1)[1:10] 
(plex2_standard*scale_factors_plex2)[1:10]

# scale intensity matrix by multiplying scale_factors for every row (protein), for every plex
plex1_irs <- sweep(m1_c_n, FUN="*", STATS=scale_factors_plex1, MARGIN=1)
plex2_irs <- sweep(m2_c_n, FUN="*", STATS=scale_factors_plex2, MARGIN=1)


# Check:
# columnmeans of the reference channels should be equal
(colMeans(plex1_irs[,10, drop=FALSE]) == colMeans(plex2_irs[,10, drop=FALSE]))
# create combined matrix
m_reporter_norm_irs <- cbind(plex1_irs, plex2_irs)

## replace former intensities with irs-normalized intensities
df_all_corr_filtered_norm_irs <- df_all_corr_filtered_norm
df_all_corr_filtered_norm_irs[,names_reporter] <- m_reporter_norm_irs
```
```{r make PCA after IRS normalisation, echo=FALSE, results='hide'}

## plot PCA
PCA_plot(m=log2(m_reporter_norm_irs),
         colorfactor = groups)
```

In addition we inspect the Pearson correlation between samples after normalisation. An overall higher variation at the level of lysine acetylation as compared to the proteome is expected, due to the general lower signal of lysine acetylated peptides and the additional aggregation of peptides on protein level. The clustering of experimental groups in the PCA and the overall rather high correlation indicate a reproducible dataset that allows further inspection in differential analysis. 

```{r Pearson correlation matrix, echo=FALSE, results='hide'}
#copy matrix
m_all_after_cor <- m_reporter_norm_irs

#the following codes for constructing Pearson correlation heatmaps is based on code from http://www.sthda.com/english/wiki/ggplot2-quick-correlation-matrix-heatmap-r-software-and-data-visualization, distributed under the creative commons license CC BY-NC-SA 3.0 US

#make names vector
cornames <- paste0(groups, "_", c(1,1,2,2,3,3,1,2,3,1,1,1,2,2,3,3,1,2,3,1))

#add proper names
colnames(m_all_after_cor) <- cornames

#remove reference channels
m_all_after_cor <- m_all_after_cor[,-c(grep(cornames, pattern = "REF"))]
m_all_after_cor <- m_all_after_cor[, order(colnames(m_all_after_cor))]

#make correlation matrix
cormat <- round(cor(m_all_after_cor, method = "pearson"), 3)

# Get lower triangle of the correlation matrix
  get_lower_tri<-function(cormat){
    cormat[upper.tri(cormat)] <- NA
    return(cormat)
  }
# Get upper triangle of the correlation matrix
  get_upper_tri <- function(cormat){
    cormat[lower.tri(cormat)]<- NA
    return(cormat)
  }
  
upper_tri <- get_upper_tri(cormat)
upper_tri

melted_cormat <- melt(upper_tri, na.rm = TRUE)

# Heatmap
# Create a ggheatmap
ggheatmap <- ggplot(melted_cormat, aes(Var2, Var1, fill = value))+
 geom_tile(color = "white")+
 scale_fill_gradient(low = "white", high = "red", 
   limit = c(0.80,1), space = "Lab", 
    name="Pearson\nCorrelation") +
  theme_minimal()+ # minimal theme
 theme(axis.text.x = element_text(angle = 90, vjust = 0.4, hjust = 1))+
 coord_fixed()+
  ggtitle("Heatmap of Pearson correlation coefficient matrix")

# Print the heatmap
print(ggheatmap)

# reorder_cormat <- function(cormat){
# # Use correlation between variables as distance
# dd <- as.dist((1-cormat)/2)
# hc <- hclust(dd)
# cormat <-cormat[hc$order, hc$order]
# }
# 
# # Reorder the correlation matrix
# cormat_re <- reorder_cormat(cormat)
# upper_tri_re <- get_upper_tri(cormat_re)
# # Melt the correlation matrix
# melted_cormat_re <- melt(upper_tri_re, na.rm = TRUE)
# # Create a ggheatmap
# ggheatmap_re <- ggplot(melted_cormat_re, aes(Var2, Var1, fill = value))+
#  geom_tile(color = "white")+
#  scale_fill_gradient(low = "white", high = "red",
#    limit = c(0.80,1), space = "Lab",
#     name="Pearson\nCorrelation") +
#   theme_minimal()+ # minimal theme
#  theme(axis.text.x = element_text(angle = 90, vjust = 0.4, hjust = 1))+
#  coord_fixed()+
#   ggtitle("Pearson correlation coefficient heatmap, clustered by coefficient")
# # Print the heatmap
# print(ggheatmap_re)

```

## 5. Site-to-protein normalisation
To be able to differentiate site-level from protein level-changes, the site data are normalized to protein data. Before this step, the the acetyl-K sites are further filtered to increase reliability and robustness of the data. This means that addition to the 1% FDR cut-off and 40 score cut-off already applied in MaxQuant, we apply a score filter of >=75, and an intensity cut-off at the 5% quantile of the mean of the three most abundant channel intensities per site.   
The normalised site-data are then again inspected by PCA.   

```{r write table IRS, echo=FALSE, results='hide'}

## generate output table
colnames(df_all_corr_filtered_norm_irs)[grepl(pattern = "Reporter.intensity_", colnames(df_all_corr_filtered_norm_irs))] <- paste0("Kac_", samplenames)
write.table(df_all_corr_filtered_norm_irs, file = paste0("./output/Acetyl(K)Sites_results_IRS.txt"), sep = "\t", col.names = TRUE, row.names=FALSE, quote=FALSE)
```


```{r ptm to protein normalisation, echo=FALSE, results='hide'}
#load data
ptm <- df_all_corr_filtered_norm_irs 
prot <- read.delim("./output/protein_groups_limma_results.txt")

dim(ptm)
ptm <- ptm[ptm$Score >= 75,] #Score >= 75, which is about about 6% of sites for this dataset
dim(ptm)

##filter sites based on intensity
#make a vector with the mean of the top three intensities per site
v_top3_int_mean <- apply(ptm[,grepl(colnames(ptm), pattern="Kac_")], 1, function(x) mean(sort(x, decreasing = TRUE)[1:3]))
#check distribution and quantiles
#hist(log10(v_top3_int_mean), breaks=50, main = "Histogram of the mean log10 intensity \n of the three most intense channels per site")
summary(v_top3_int_mean)
quantile(v_top3_int_mean, probs = c(0.005, 0.01, 0.05))
# apply cut-off at 5% quantile (above intensity of 1000)
int_cut_off <- quantile(v_top3_int_mean, probs = c(0.05))
ptm <- ptm[v_top3_int_mean >= int_cut_off,]
dim(ptm)

#check whether there are missing values in the PTM data, FALSE means no
sum(apply(ptm[,grepl(colnames(ptm), pattern="Reporter.intensity")], 2, function(x) sum(x == 0))) != 0
#check for NAs
v_NA_ptm <- is.na(rowSums(ptm[,grep(colnames(ptm), pattern = "Reporter.intensity")]))
table(v_NA_ptm)

#### Filtering of protein data already performed before####

#### Merge PTM with proteome data ####

#reduce to those columns needed
prot <- prot[, c(which(colnames(prot)=="id"), which(colnames(prot)=="Razor...unique.peptides"), c(5:24), grep(colnames(prot), pattern="logFC"), grep(colnames(prot), pattern="P.Value"), grep(colnames(prot), pattern="adj.P.Val"))]

#rename all columns to be distinguishable from phos-sites table with a "P_" prefix
colnames(prot) <- paste0("P_", colnames(prot))

#problem: in the sites table some entries have multiple protein IDs. simple matching by merge does not work. for this reason first the best match needs to be selected. 1. find all entries with ";", 2. split them, 3. search which of those has highest number of razor peptides, 4. make a new column with only one entry, 5. use that for merge.

#check how many of them there are
length(grep(";", ptm$Protein.group.IDs,))

l_protids_dups <- strsplit(ptm$Protein.group.IDs, split = ";") #create list that splits multiple entries separated by ";"

Protein.group.IDs_quant <- character(nrow(ptm)) #initialize vector for loop

for (i in 1:length(l_protids_dups)){
  l_i <- as.numeric(l_protids_dups[[i]]) #converts list to numeric!
  df_i <- prot[prot$P_id %in% l_i,] #creates df from prot for proteins in list at position i
  ind_id <- which.max(df_i$P_Razor...unique.peptides) #writes index for df with highest number of Razor peptides
  if (length(df_i$P_id[ind_id]!=0)) {
    Protein.group.IDs_quant[i] <- df_i$P_id[ind_id]
  } else {
    Protein.group.IDs_quant[i] <- NA
  } #writes protein groups id with highest number of razor in vector; if there is no matching protein ID (because it was filtered out before, it will write NA instead)
}

table(is.na(Protein.group.IDs_quant)) # check how many NAs, i.e. sites without protein info, are there

#add vector to original df
ptm <- cbind(ptm, Protein.group.IDs_quant)

#merge both tables, omit sites with no protein match
ptm_prot <- merge(ptm, prot, by.x = "Protein.group.IDs_quant", by.y = "P_id")
#same but keep sites without protein match for documentation
ptm_prot_allsites <- merge(ptm, prot, by.x = "Protein.group.IDs_quant", by.y = "P_id", all.x = TRUE)

#calculate mean protein reporter intensity for each row
P_mean_int <- rowMeans(ptm_prot[, grepl(colnames(ptm_prot), pattern="^P_.*_[1-3]")], na.rm = TRUE)

#extract site and protein reporter intensities for further calculations
ptm_int <- ptm_prot[,grep("^Kac_", colnames(ptm_prot))]
prot_int <- ptm_prot[,grep("^P_.*_[1-3]", colnames(ptm_prot))]
#divide site intensity by protein intensity
ptm_int_n <- ptm_int / prot_int
#multiply each value again by mean protein intensity
ptm_int_n <- ptm_int_n * P_mean_int 
#plot(log2(ptm_int[,1]), log2(ptm_int_n[,1])) #just to check normalisation visually
#pairs(log2(ptm_int_n[,c(1,3,5,2,4,6)]), cex=0.1)

#rename colnames
colnames(ptm_int_n) <- paste0("pnorm_", colnames(ptm_int_n))

#add values back to df
ptm_prot <- data.frame(ptm_prot, ptm_int_n)

## plot PCA
PCA_plot(m=log2(ptm_int_n),
         colorfactor = groups)

#### write output tables ####
#sites and proteins matched, protein-normalized sites included rows with missing protein values omitted
write.table(ptm_prot, file = "./output/acK-sites_filtered_proteinnormalized.txt", row.names = FALSE, quote = FALSE, sep = "\t")
# #sites and proteins matched, all rows included
# write.table(ptm_prot_allsites, file = "acK-sites_matched_to_proteins_full_MS2.txt", row.names = FALSE, quote = FALSE, sep = "\t")

```

## 6. Differential abundance analysis using LIMMA

LIMMA is performed with the "trend" option and using the following model: (~ 0 + group).

```{r Do Differential expression testing, echo=FALSE, results='hide'}

#copy and log transform data
d_limma <- log2(ptm_int_n)
head(d_limma)

# set up the design matrix
group <- as.factor(make.names(groups))
#batch <- as.factor(rep(c(1,2), each = 10))
design <- model.matrix(~ 0 + group)
colnames(design) <- sub(pattern = ("group"), replacement = "", colnames(design))
colnames(design)
design

#limma
# do the linear model fitting
fit <- lmFit(d_limma, design)

# make the contrast
comparisons <- c("Gse1_KO_ctrl-WT_ctrl", "HDAC1.CI_ctrl-WT_ctrl", "Gse1_KO_ctrl-HDAC1.CI_ctrl", "Gse1_KO_eto-WT_eto", "HDAC1.CI_eto-WT_eto", "Gse1_KO_eto-HDAC1.CI_eto", "Gse1_KO_eto-Gse1_KO_ctrl", "HDAC1.CI_eto-HDAC1.CI_ctrl", "WT_eto-WT_ctrl")

contrast <- makeContrasts(contrasts = comparisons, levels = design)

# get the fit for the contrast of interest
fit2 <- contrasts.fit(fit, contrast)

# do the empirical Bayes moderation of the test statistic (with trended variance)
fit2 <- eBayes(fit2, trend = TRUE)

# show how many proteins are differentially abundant for each contrast
summary(decideTests(fit2))

# store results
tt_exp_limma <- topTable(fit2, coef = 1, sort.by = "none", number = Inf) #writes toptable for coeff1

for(i in 2:length(comparisons)) {
  tt_exp_limma <- data.frame(tt_exp_limma, topTable(fit2, coef = i, sort.by = "none", number = Inf))
  print(length(colnames(tt_exp_limma)))
} #adds toptable for remaining coefficients

colnames(tt_exp_limma) <- paste0(colnames(tt_exp_limma)[1:6],".", rep(comparisons, each = 6)) #rename columns by adding comparisons

#combine limma data with full table
df_full_set <- data.frame(ptm_prot, tt_exp_limma)

#The following comparisons were made:  
comparisons

#Significantly regulated acetylated peptides (at 5% FDR) for all comparisons:  
summary(decideTests(fit2))

```

The following significantly regulated sites (at 5% FDR) were determined:  
`r kable(summary(decideTests(fit2)))`

The results are stored to: "acK-sites_filtered_proteinnormalized_LIMMA.txt" as well as in a reduced format as "acK-sites_filtered_proteinnormalized_LIMMA_condensed.txt".

### Volcano plots
Plots for all comparisons of interest are generated and stored, marking significant sites (at 5% FDR and > 2-fold or < 0.5-fold difference).  

```{r write final table, echo=FALSE, results='hide', warning=FALSE, message=FALSE}

## generate full output table
write.table(df_full_set, file = "./output/acK-sites_filtered_proteinnormalized_LIMMA.txt", col.names = TRUE, row.names = FALSE, quote = FALSE, sep = "\t")

##generate condensed output table
df_full_condensed <- df_full_set 
#make vector of desired column names
sel_columns <- c("Proteins", "Positions.within.proteins", "Leading.proteins", 
"Protein", "Protein.names", "Gene.names", "Fasta.headers", "Localization.prob", 
"Score.diff", "PEP", "Score", "Delta.score", "Score.for.localization", 
"Number.of.Acetyl..K.", "Sequence.window", "Acetyl..K..Probabilities", 
"Position.in.peptide", "id", "Protein.group.IDs", "Positions", 
"Position", "Number.Sites", "P_Razor...unique.peptides", "P_logFC.Gse1_KO_ctrl.WT_ctrl", 
"P_logFC.HDAC1.CI_ctrl.WT_ctrl", "P_logFC.Gse1_KO_ctrl.HDAC1.CI_ctrl", 
"P_logFC.Gse1_KO_eto.WT_eto", "P_logFC.HDAC1.CI_eto.WT_eto", 
"P_logFC.Gse1_KO_eto.HDAC1.CI_eto", "P_logFC.Gse1_KO_eto.Gse1_KO_ctrl", 
"P_logFC.HDAC1.CI_eto.HDAC1.CI_ctrl", "P_logFC.WT_eto.WT_ctrl", 
"P_P.Value.Gse1_KO_ctrl.WT_ctrl", "P_P.Value.HDAC1.CI_ctrl.WT_ctrl", 
"P_P.Value.Gse1_KO_ctrl.HDAC1.CI_ctrl", "P_P.Value.Gse1_KO_eto.WT_eto", 
"P_P.Value.HDAC1.CI_eto.WT_eto", "P_P.Value.Gse1_KO_eto.HDAC1.CI_eto", 
"P_P.Value.Gse1_KO_eto.Gse1_KO_ctrl", "P_P.Value.HDAC1.CI_eto.HDAC1.CI_ctrl", 
"P_P.Value.WT_eto.WT_ctrl", "P_adj.P.Val.Gse1_KO_ctrl.WT_ctrl", 
"P_adj.P.Val.HDAC1.CI_ctrl.WT_ctrl", "P_adj.P.Val.Gse1_KO_ctrl.HDAC1.CI_ctrl", 
"P_adj.P.Val.Gse1_KO_eto.WT_eto", "P_adj.P.Val.HDAC1.CI_eto.WT_eto", 
"P_adj.P.Val.Gse1_KO_eto.HDAC1.CI_eto", "P_adj.P.Val.Gse1_KO_eto.Gse1_KO_ctrl", 
"P_adj.P.Val.HDAC1.CI_eto.HDAC1.CI_ctrl", "P_adj.P.Val.WT_eto.WT_ctrl", 
"pnorm_Kac_Gse1_KO_ctrl_1", "pnorm_Kac_Gse1_KO_ctrl_2", "pnorm_Kac_Gse1_KO_ctrl_3", 
"pnorm_Kac_Gse1_KO_eto_1", "pnorm_Kac_Gse1_KO_eto_2", "pnorm_Kac_Gse1_KO_eto_3", 
"pnorm_Kac_HDAC1.CI_ctrl_1", "pnorm_Kac_HDAC1.CI_ctrl_2", "pnorm_Kac_HDAC1.CI_ctrl_3", 
"pnorm_Kac_HDAC1.CI_eto_1", "pnorm_Kac_HDAC1.CI_eto_2", "pnorm_Kac_HDAC1.CI_eto_3", 
"pnorm_Kac_REF_ctrl_1", "pnorm_Kac_REF_eto_1", "pnorm_Kac_WT_ctrl_1", 
"pnorm_Kac_WT_ctrl_2", "pnorm_Kac_WT_ctrl_3", "pnorm_Kac_WT_eto_1", 
"pnorm_Kac_WT_eto_2", "pnorm_Kac_WT_eto_3", "logFC.Gse1_KO_ctrl.HDAC1.CI_ctrl", 
"logFC.Gse1_KO_ctrl.WT_ctrl", "logFC.Gse1_KO_eto.Gse1_KO_ctrl", 
"logFC.Gse1_KO_eto.HDAC1.CI_eto", "logFC.Gse1_KO_eto.WT_eto", 
"logFC.HDAC1.CI_ctrl.WT_ctrl", "logFC.HDAC1.CI_eto.HDAC1.CI_ctrl", 
"logFC.HDAC1.CI_eto.WT_eto", "logFC.WT_eto.WT_ctrl", "P.Value.Gse1_KO_ctrl.HDAC1.CI_ctrl", 
"P.Value.Gse1_KO_ctrl.WT_ctrl", "P.Value.Gse1_KO_eto.Gse1_KO_ctrl", 
"P.Value.Gse1_KO_eto.HDAC1.CI_eto", "P.Value.Gse1_KO_eto.WT_eto", 
"P.Value.HDAC1.CI_ctrl.WT_ctrl", "P.Value.HDAC1.CI_eto.HDAC1.CI_ctrl", 
"P.Value.HDAC1.CI_eto.WT_eto", "P.Value.WT_eto.WT_ctrl", "adj.P.Val.Gse1_KO_ctrl.HDAC1.CI_ctrl", 
"adj.P.Val.Gse1_KO_ctrl.WT_ctrl", "adj.P.Val.Gse1_KO_eto.Gse1_KO_ctrl", 
"adj.P.Val.Gse1_KO_eto.HDAC1.CI_eto", "adj.P.Val.Gse1_KO_eto.WT_eto", 
"adj.P.Val.HDAC1.CI_ctrl.WT_ctrl", "adj.P.Val.HDAC1.CI_eto.HDAC1.CI_ctrl", 
"adj.P.Val.HDAC1.CI_eto.WT_eto", "adj.P.Val.WT_eto.WT_ctrl", 
"AveExpr.Gse1_KO_ctrl.HDAC1.CI_ctrl", "AveExpr.Gse1_KO_ctrl.WT_ctrl", 
"AveExpr.Gse1_KO_eto.Gse1_KO_ctrl", "AveExpr.Gse1_KO_eto.HDAC1.CI_eto", 
"AveExpr.Gse1_KO_eto.WT_eto", "AveExpr.HDAC1.CI_ctrl.WT_ctrl", 
"AveExpr.HDAC1.CI_eto.HDAC1.CI_ctrl", "AveExpr.HDAC1.CI_eto.WT_eto", 
"AveExpr.WT_eto.WT_ctrl")

#check for entries with missing Gene name
df_full_condensed[df_full_condensed$Gene.names == "", c(1,7)]
gn_empty <- length(df_full_condensed[df_full_condensed$Gene.names == "", "Gene.names"]) #number of empty gene names

#replace empty gene names with [none] followed by a counter
df_full_condensed[df_full_condensed$Gene.names == "", "Gene.names"] <- paste0("[none]",c(1:gn_empty))
df_full_condensed[grepl(df_full_condensed$Gene.names, pattern = "none"), c(1,7)]

#check for multiple entries in gene name
table(grepl(df_full_condensed$Gene.names, pattern = ";"))
#make a vector with single gene names
single_gene_names <- sub(";.*", "", df_full_condensed$Gene.names)

#make unique site identifier
unique_site_ID <- paste0(single_gene_names, "_K", df_full_set$Position, "_", df_full_set$Number.Sites, "ac")

#reduce to desired columns
df_full_condensed <- df_full_condensed[, colnames(df_full_condensed) %in% sel_columns]

#add unique site ID
df_full_condensed <- cbind(unique_site_ID, df_full_condensed)
colnames(df_full_condensed)
colnames(df_full_condensed)[52:107] <- paste0("Kac_", colnames(df_full_condensed[52:107]))
colnames(df_full_condensed)

#write output table
write.table(df_full_condensed, file = "./output/acK-sites_filtered_proteinnormalized_LIMMA_condensed.txt", col.names = TRUE, row.names = FALSE, quote = FALSE, sep = "\t")
```
``` {r volcano plots, echo=FALSE, results='hide', warning=FALSE, message=FALSE}
#Volcano plots with p-values (not corrected for multiple testing) or adj.p.values (corrected). Red indicates p-values or adjusted p-values below 0.05, blue above, respectively

i_logFC <- grep("logFC", colnames(tt_exp_limma))

for (i in i_logFC) { 

    #define regulated datapoints: all sites >|< +/-1 log2 FC and significant in LIMMA
    b_reg <- (abs(tt_exp_limma[,i]) >= 1) & (tt_exp_limma[,i+4] <= 0.05 )

    #change level information
    levels(b_reg) <- c("non-signif.", "signif.")
  
    #volcano plot
    p <- ggplot(tt_exp_limma, mapping = aes(x= tt_exp_limma[,i], y=-log(tt_exp_limma[,i+3], base =10))) +
      geom_vline(xintercept = -1, linetype = "solid") +
      geom_vline(xintercept = 1, linetype = "solid") +
      geom_point(tt_exp_limma[!b_reg,], 
                 alpha = 0.2, pch = 16, size = 2, 
                 mapping = aes(x=tt_exp_limma[!b_reg,i], y = -log(tt_exp_limma[!b_reg,i+3], base =10), col=b_reg[!b_reg])) +
      geom_point(tt_exp_limma[b_reg,], 
                 alpha = 0.5, pch = 16, size = 2, 
                 mapping = aes(x=tt_exp_limma[b_reg,i], y=-log(tt_exp_limma[b_reg,i+3], base =10), col=b_reg[b_reg])) +
      scale_color_manual(values=c("gray", "red")) +
      labs(title = sub("-", " vs. ", sub("logFC.", "", colnames(tt_exp_limma)[i])), color = "Significant acetylation sites (<5% FDR):") +
      theme_classic() +
      scale_x_continuous(name="Acetylation site log2 fold-change", limits=c(min(tt_exp_limma[,i_logFC]) - 0.5, max(tt_exp_limma[,i_logFC]) + 0.5), n.breaks = round(max(tt_exp_limma[,i_logFC]) + abs(min(tt_exp_limma[,i_logFC])))) +
      scale_y_continuous(name="-log10 LIMMA p-value", limits=c(0, max(-log10(tt_exp_limma[,i_logFC+3])) + 0.5)) 
    
    print(p)
    ggsave(plot = p, path = "output/figs", filename = paste0("Kac_", colnames(tt_exp_limma)[i], ".pdf"))
}

```
````
