## Supplementary material for "GSE1 links the HDAC1/CoREST co-repressor complex to DNA damage": Scripts used for MS shotgun data analysis: 03_phosphosites_data_processing_ver01.html

Quantitative analysis of protein phosporylation sites in HAP1 cells +/- etoposide treatment in different genetic backgrounds


### Quantitative analysis of protein phosporylation sites in HAP1 cells +/- etoposide treatment in different genetic backgrounds

#### 6. Differential abundance analysis using LIMMA

LIMMA is performed with the “trend” option and using the following
model: (~ 0 + group).

The following significantly regulated sites (at 5% FDR) were
determined:

|  | Gse1\_KO\_ctrl-WT\_ctrl | HDAC1.CI\_ctrl-WT\_ctrl | Gse1\_KO\_ctrl-HDAC1.CI\_ctrl | Gse1\_KO\_eto-WT\_eto | HDAC1.CI\_eto-WT\_eto | Gse1\_KO\_eto-HDAC1.CI\_eto | Gse1\_KO\_eto-Gse1\_KO\_ctrl | HDAC1.CI\_eto-HDAC1.CI\_ctrl | WT\_eto-WT\_ctrl |
| --- | --- | --- | --- | --- | --- | --- | --- | --- | --- |
| Down | 6597 | 4591 | 5536 | 6567 | 4742 | 6248 | 2756 | 3921 | 4825 |
| NotSig | 14004 | 15968 | 15663 | 13989 | 15642 | 14299 | 20187 | 17524 | 15269 |
| Up | 4001 | 4043 | 3403 | 4046 | 4218 | 4055 | 1659 | 3157 | 4508 |

The results are stored to:
“acK-sites\_filtered\_proteinnormalized\_LIMMA.txt” as well as in a reduced
format as
“acK-sites\_filtered\_proteinnormalized\_LIMMA\_condensed.txt”.

##### Volcano plots

Plots for all comparisons of interest are generated and stored,
marking significant sites (at 5% FDR and > 2-fold or < 0.5-fold
difference).
