## Supplementary material for "GSE1 links the HDAC1/CoREST co-repressor complex to DNA damage": Scripts used for MS shotgun data analysis: readme.txt

This folder contains all data to reproduce the data analysis after MaxQuant search. The three R-markdown script in this root directory should be run consecutively (1, 2, 3) on the data in the "input" folder and will generate all files in the "output" folder as well as the html files when used with the knitr package (e.g. in RStudio). The html-files contain a commented summary of the analysis, including plots for illustration. 
