## Supplementary figures and images for "GSE1 links the HDAC1/CoREST co-repressor complex to DNA damage"

### Kac_logFC.Gse1_KO_ctrl-HDAC1.CI_ctrl.pdf

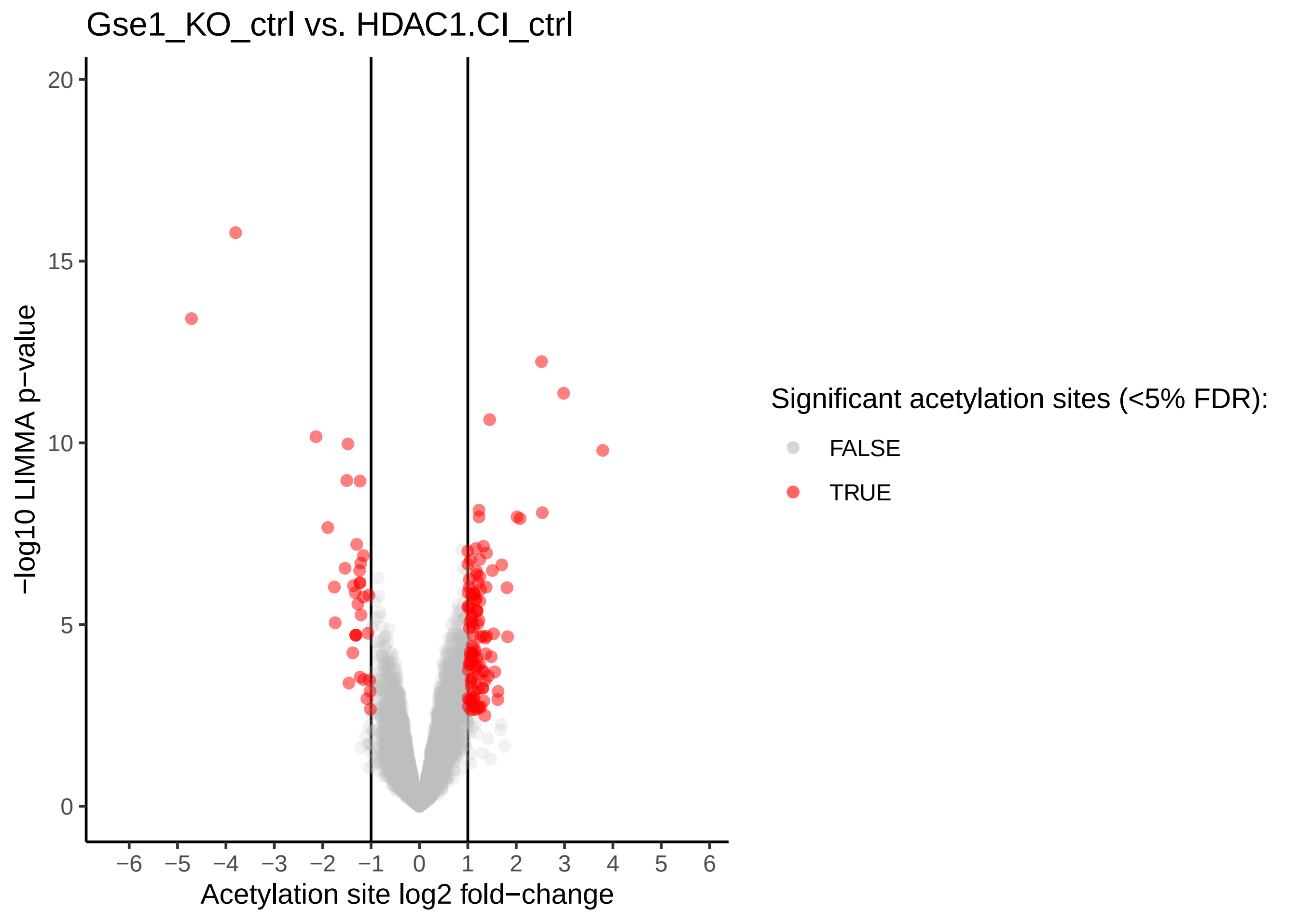

### Kac_logFC.Gse1_KO_ctrl-WT_ctrl.pdf

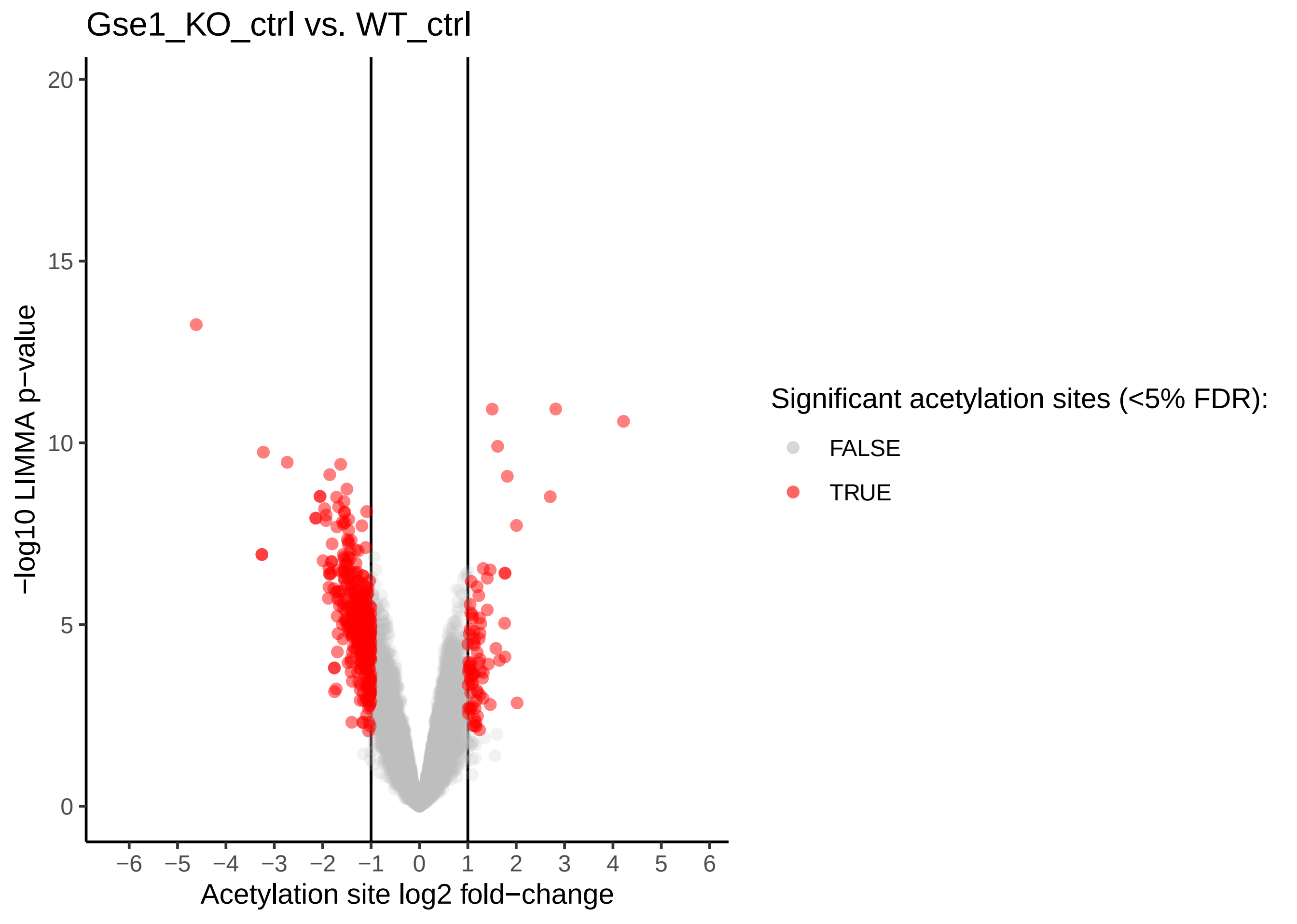

### Kac_logFC.Gse1_KO_eto-Gse1_KO_ctrl.pdf

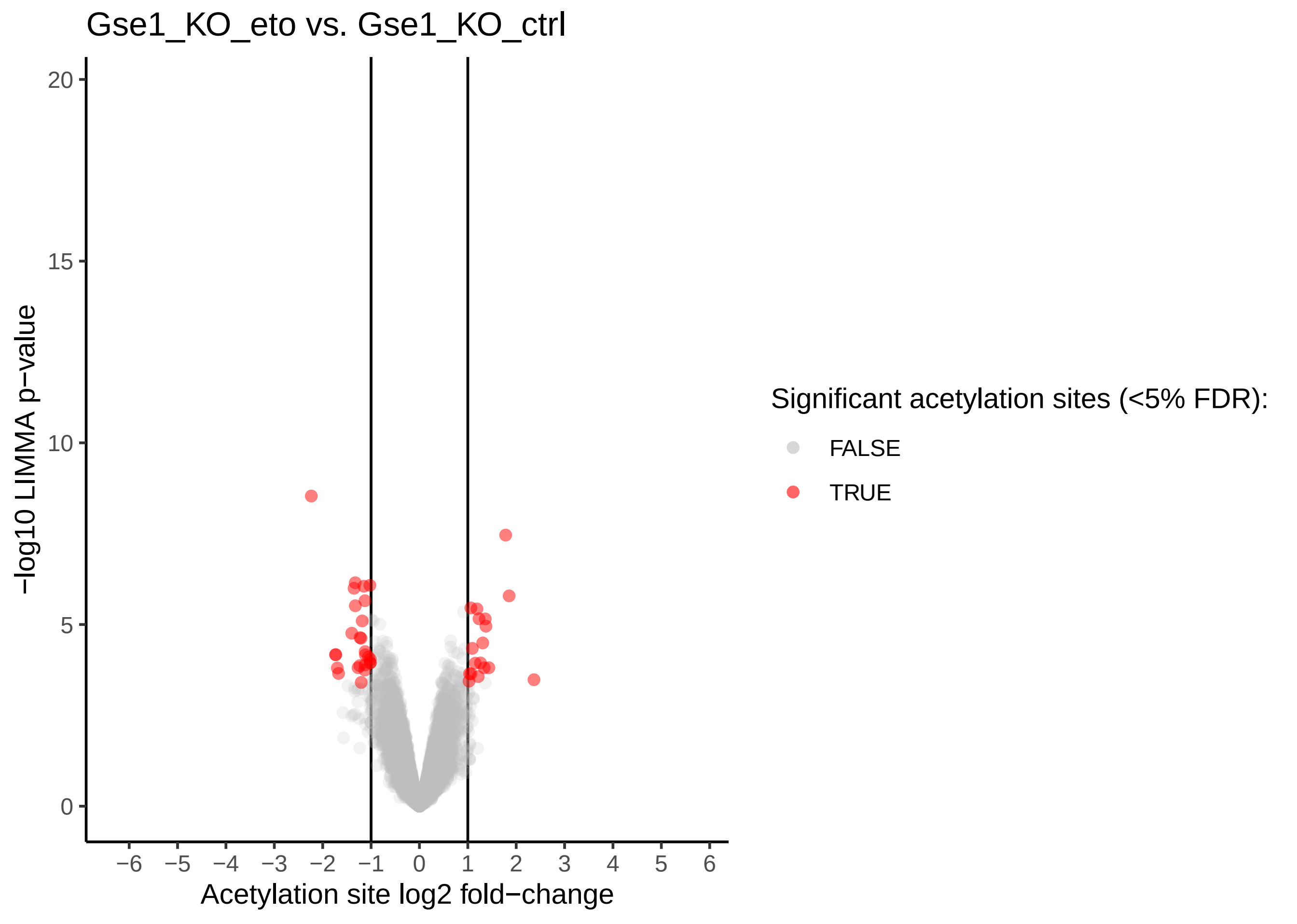

### Kac_logFC.Gse1_KO_eto-HDAC1.CI_eto.pdf

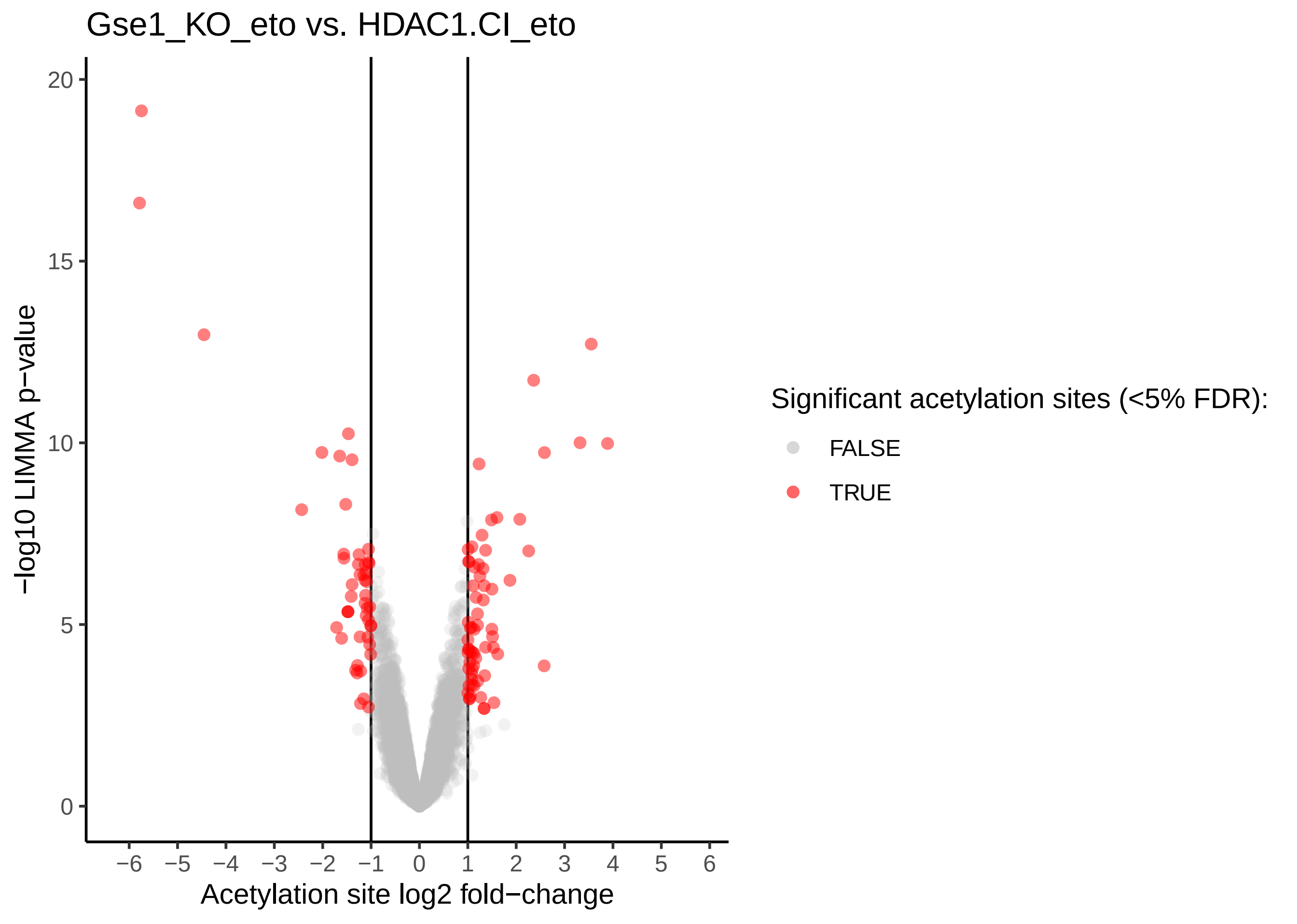

### Kac_logFC.Gse1_KO_eto-WT_eto.pdf

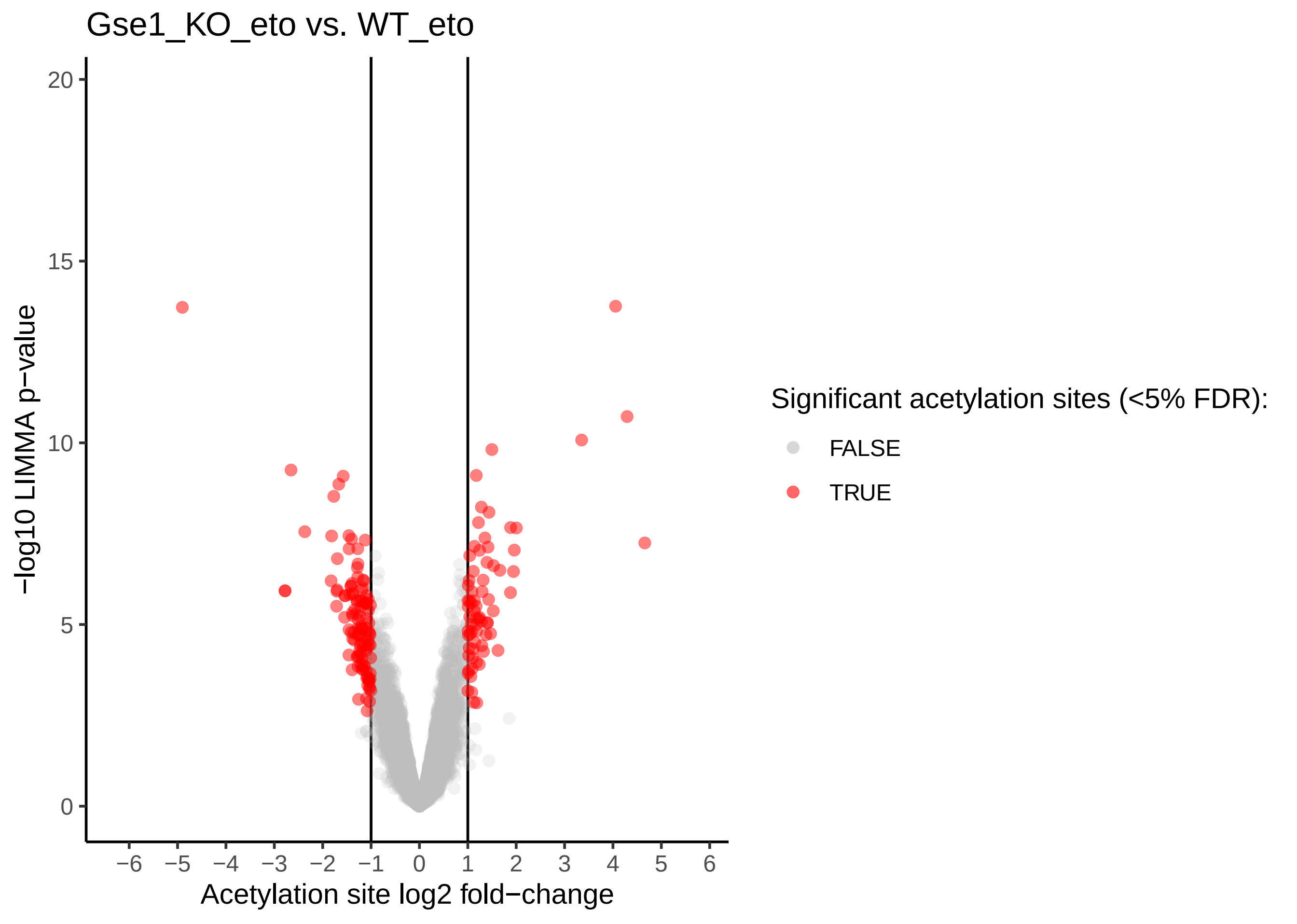

### Kac_logFC.HDAC1.CI_ctrl-WT_ctrl.pdf

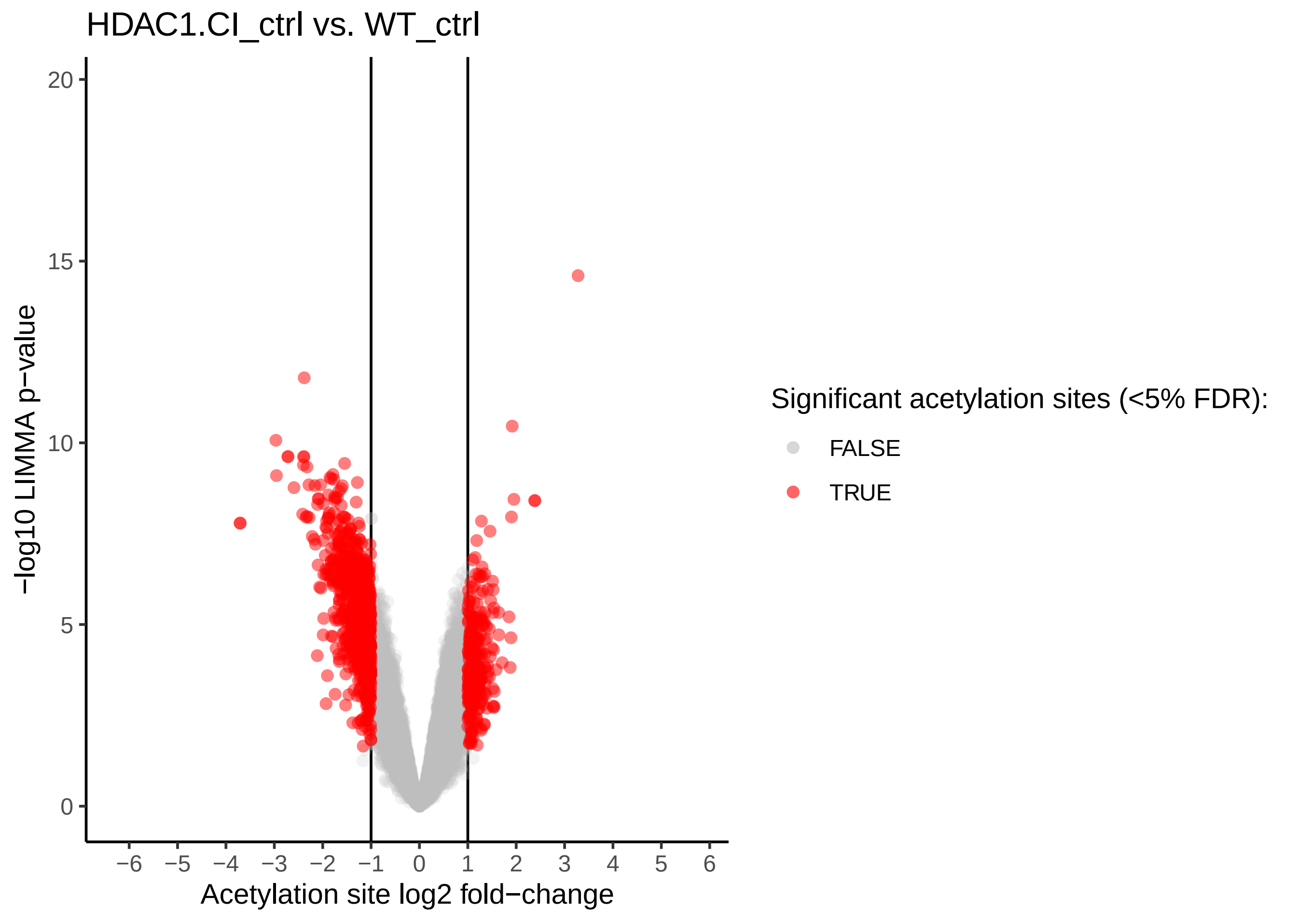

### Kac_logFC.HDAC1.CI_eto-HDAC1.CI_ctrl.pdf

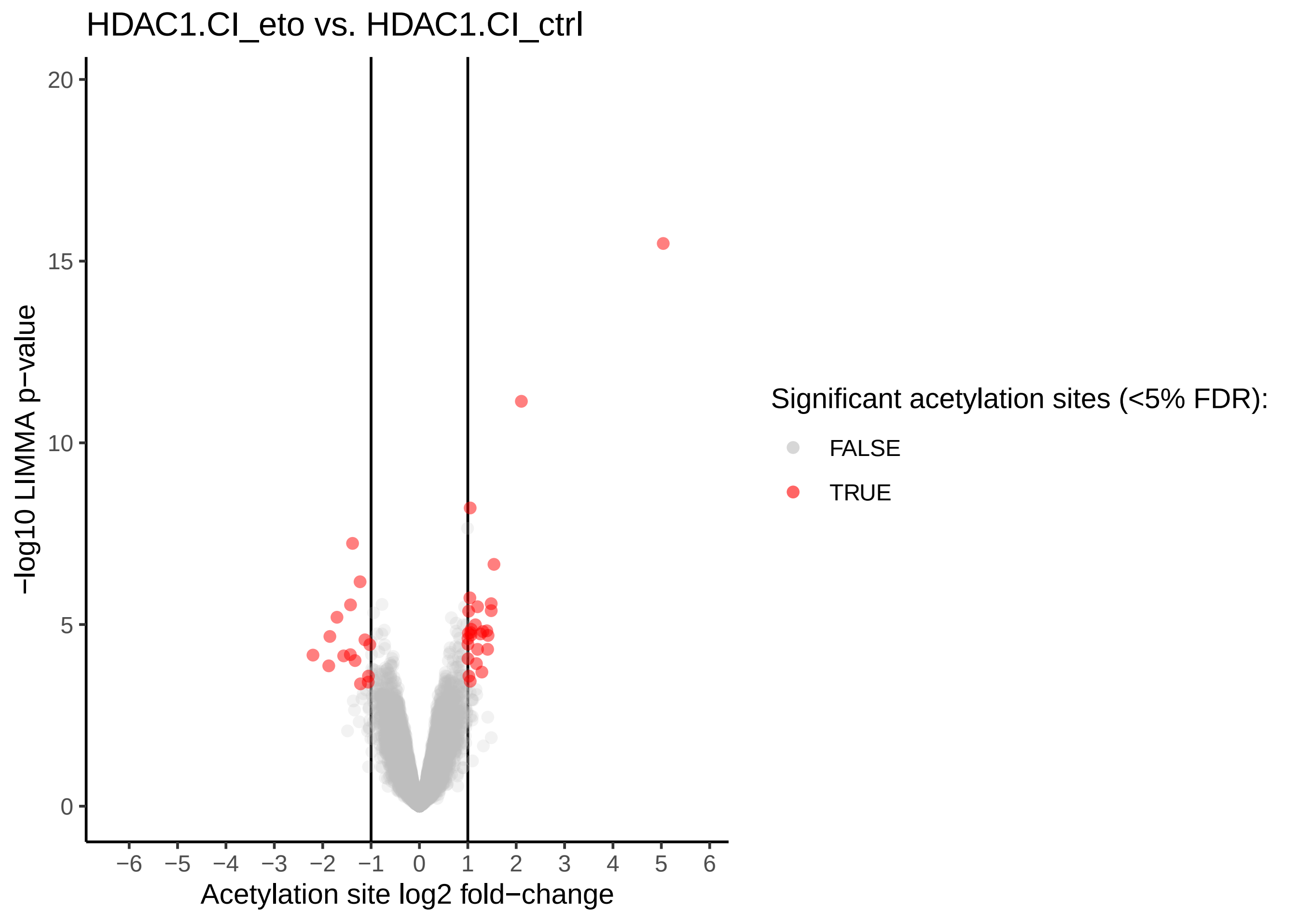

### Kac_logFC.HDAC1.CI_eto-WT_eto.pdf

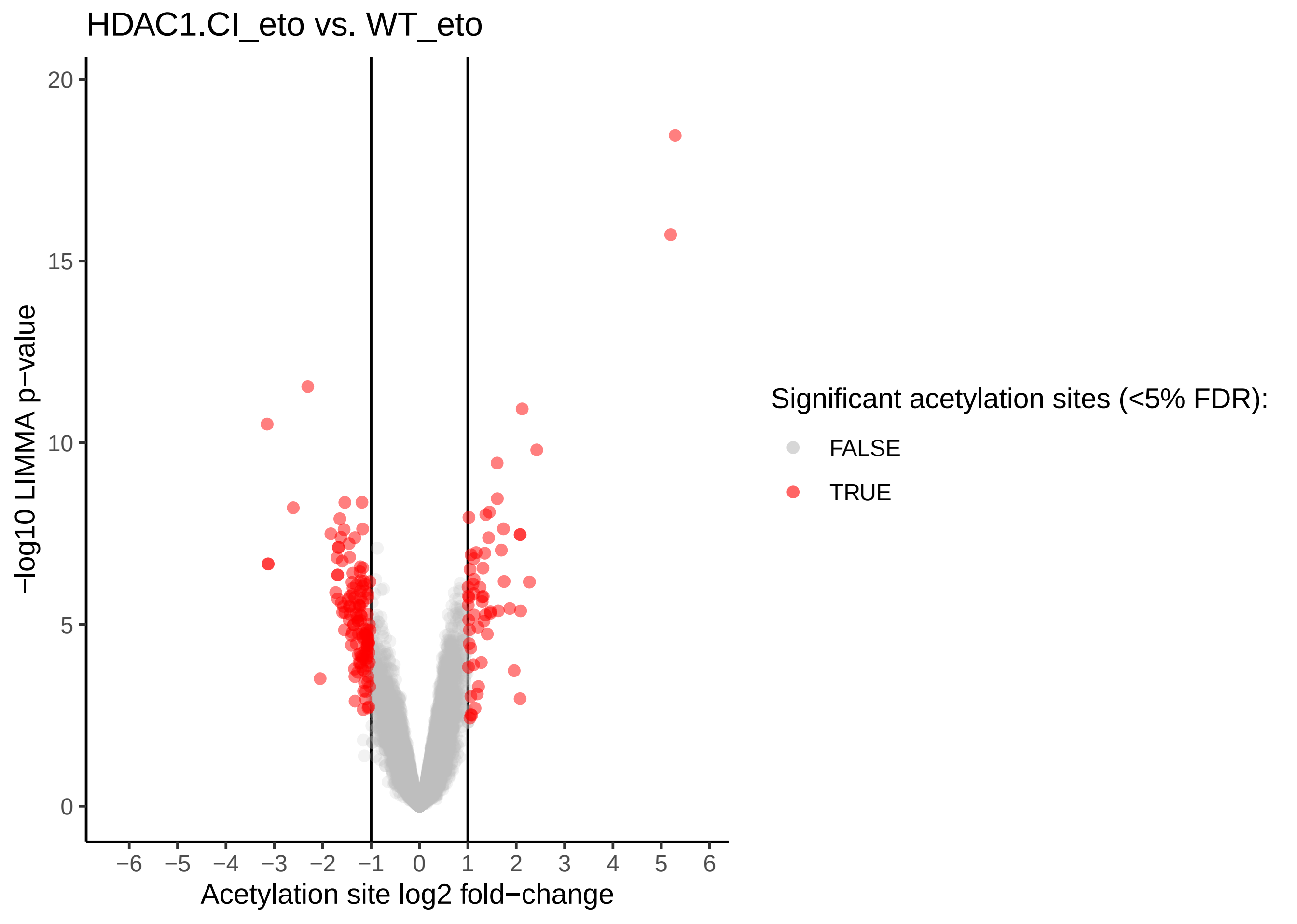

### Kac_logFC.WT_eto-WT_ctrl.pdf

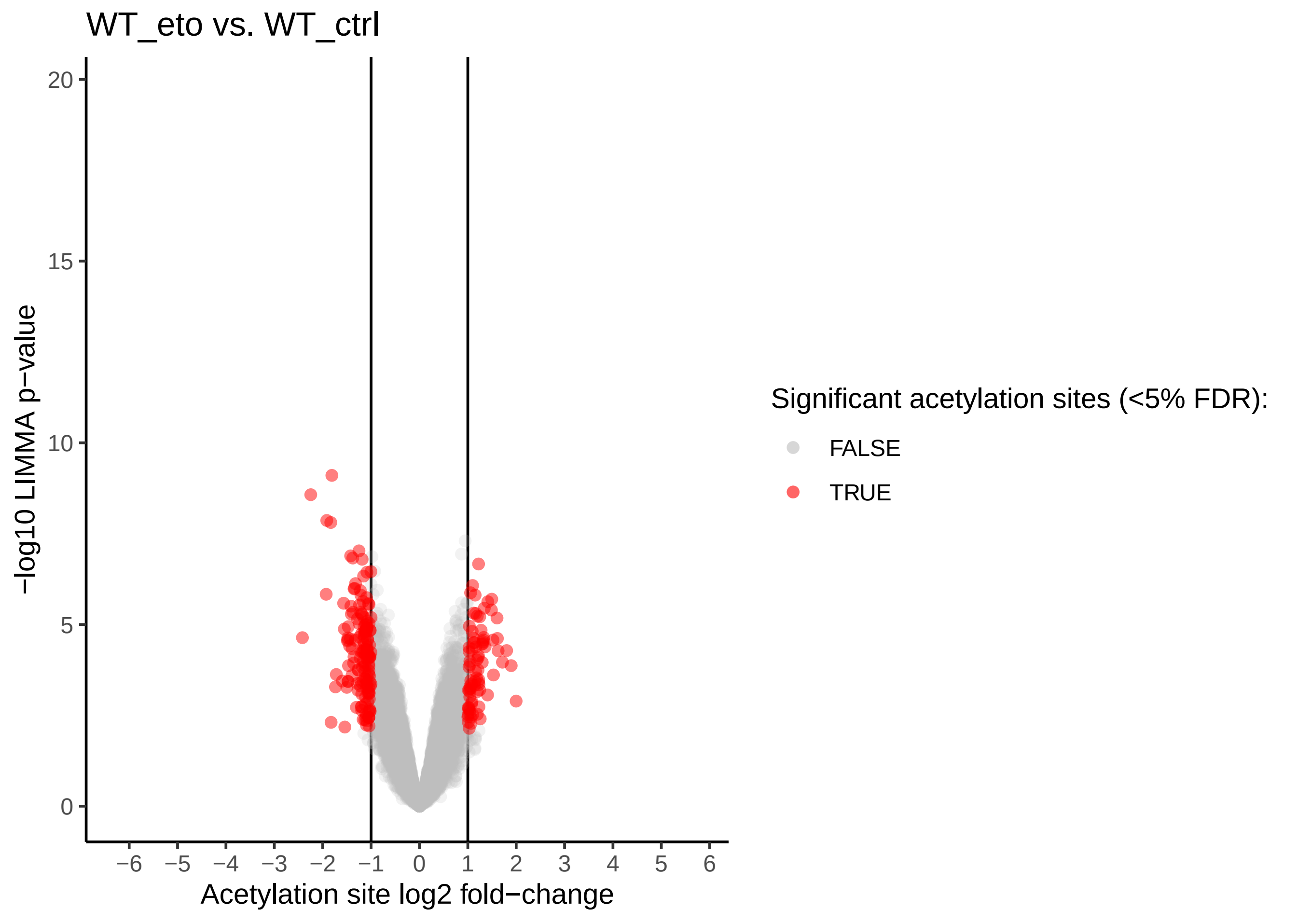

### ph_logFC.Gse1_KO_ctrl-HDAC1.CI_ctrl.pdf

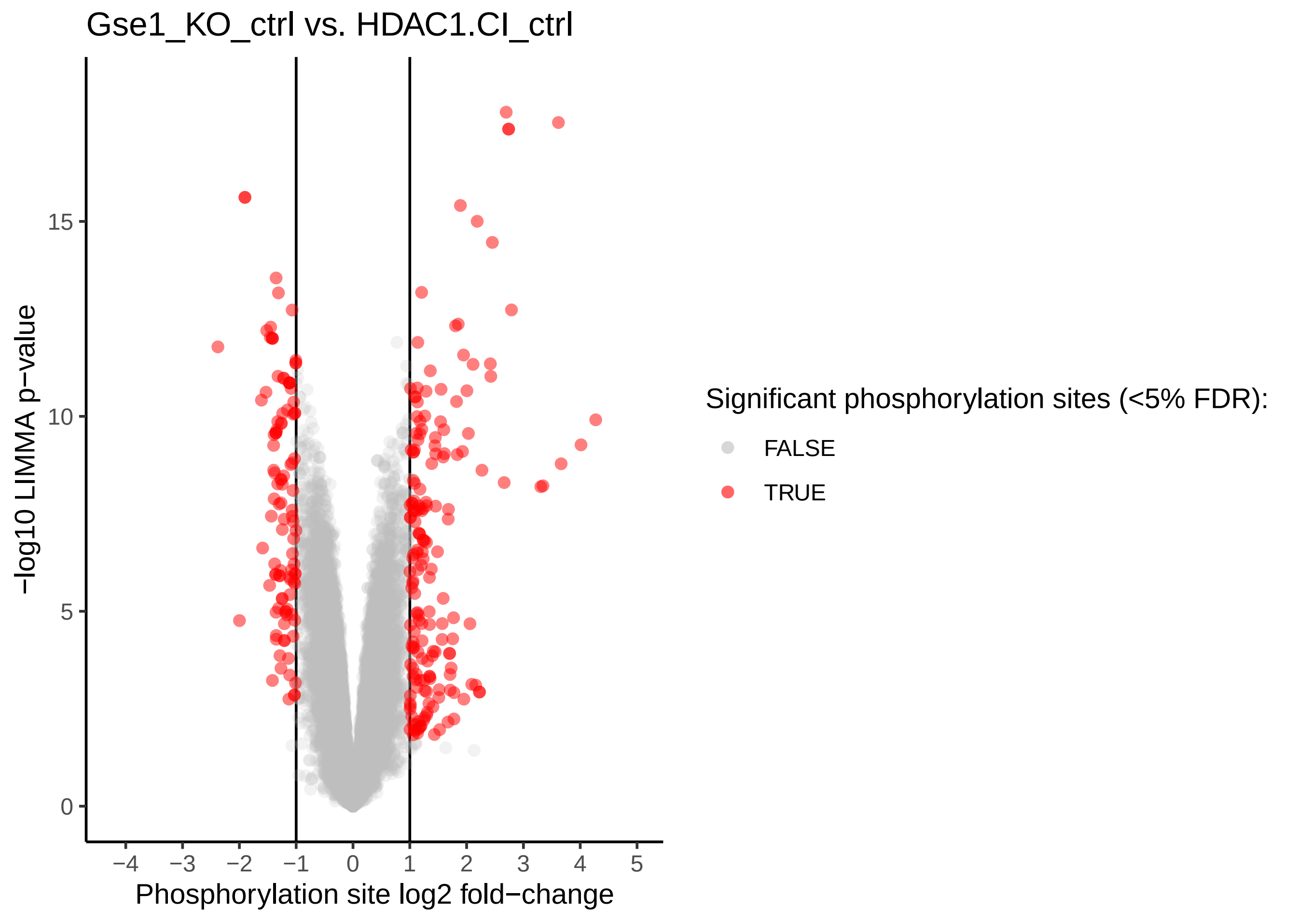

### ph_logFC.Gse1_KO_ctrl-WT_ctrl.pdf

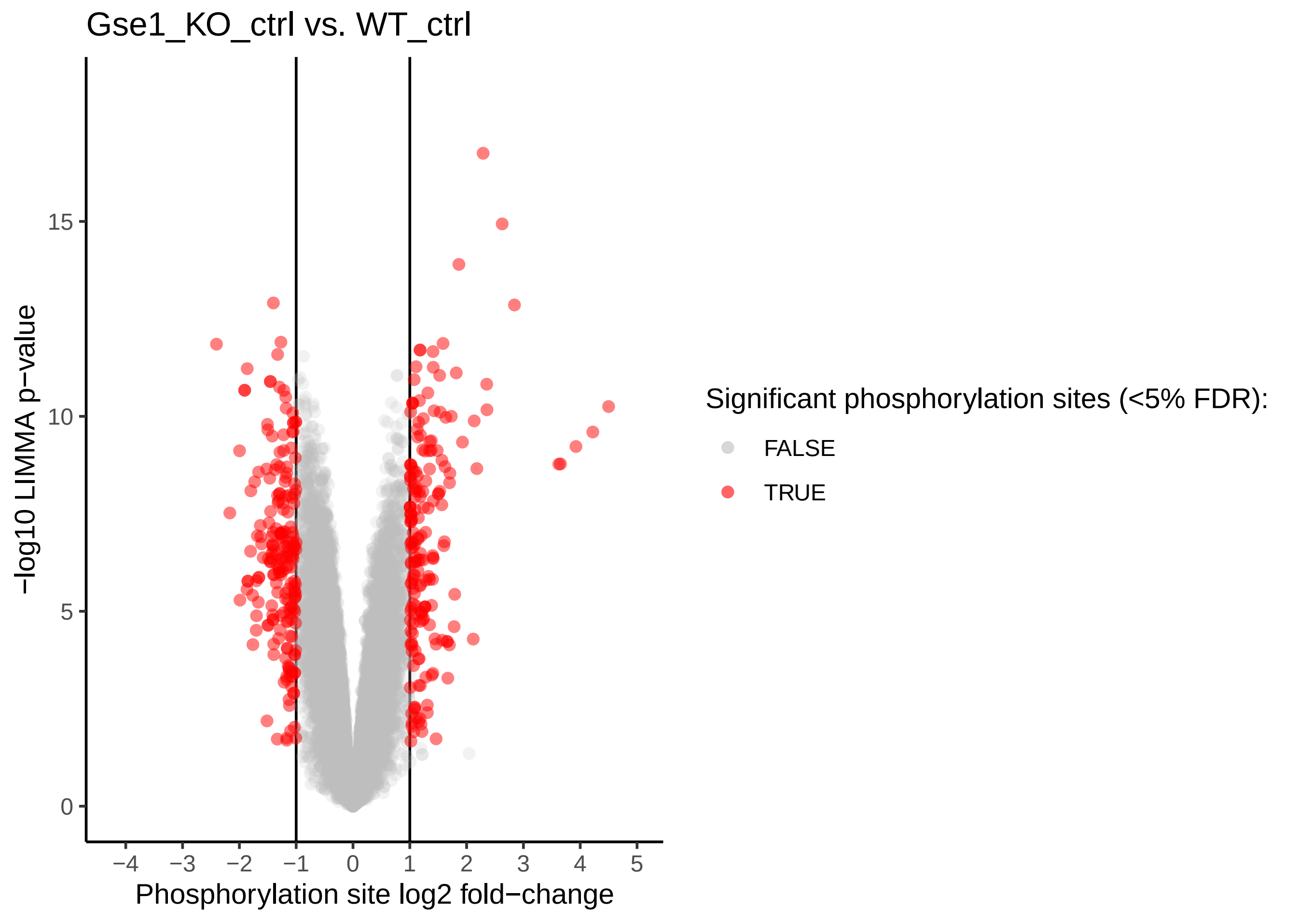

### ph_logFC.Gse1_KO_eto-Gse1_KO_ctrl.pdf

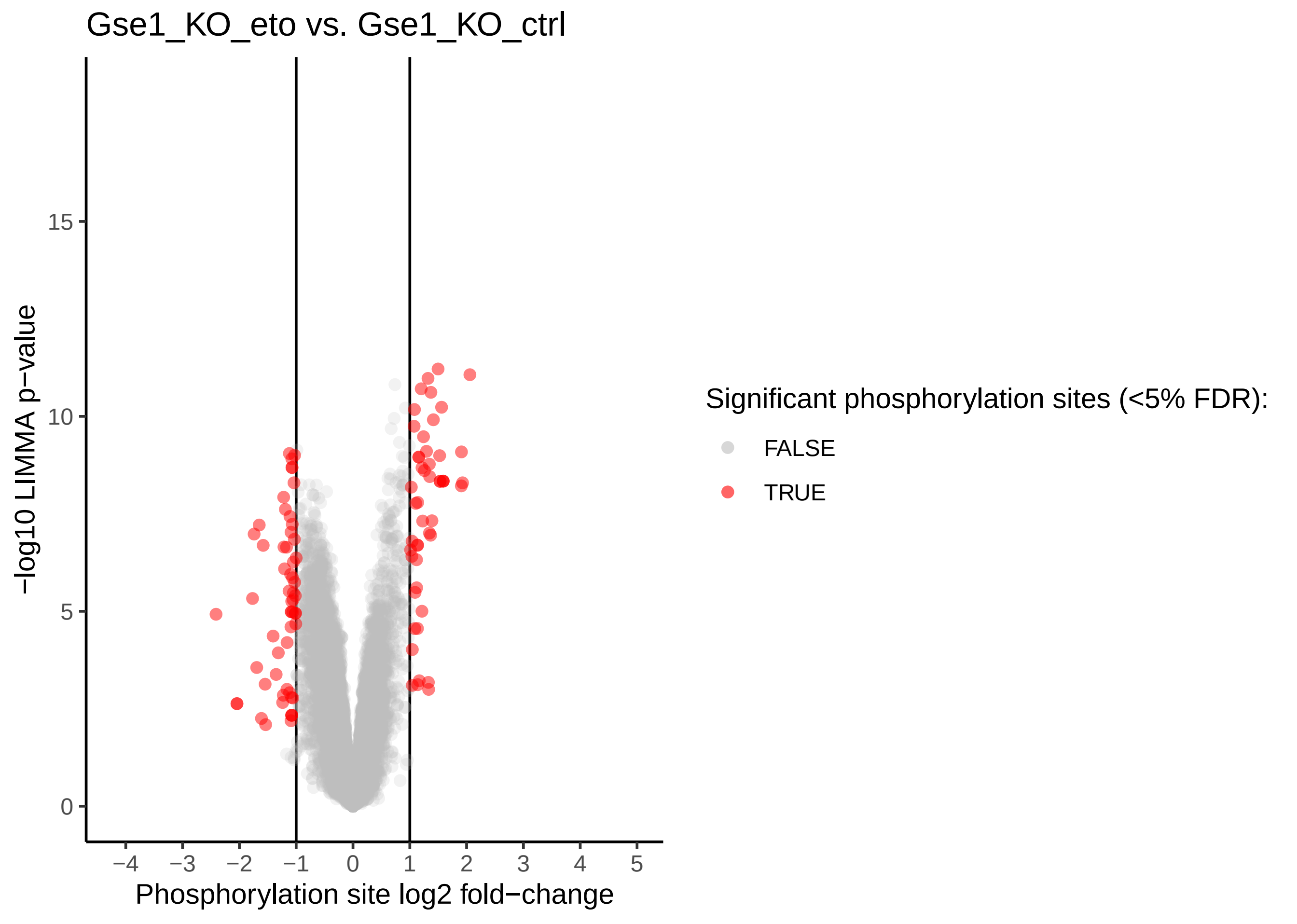

### ph_logFC.Gse1_KO_eto-HDAC1.CI_eto.pdf

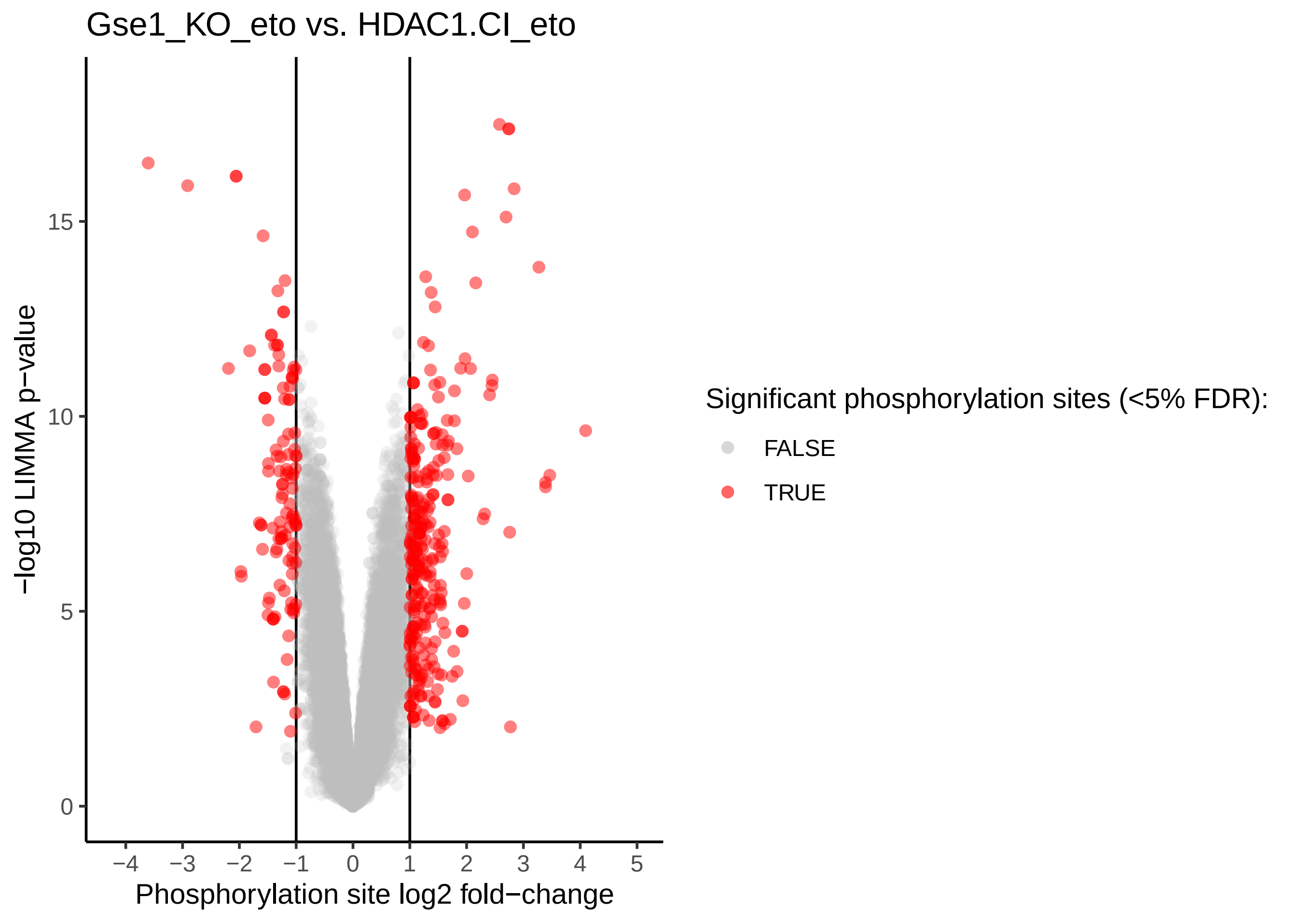

### ph_logFC.Gse1_KO_eto-WT_eto.pdf

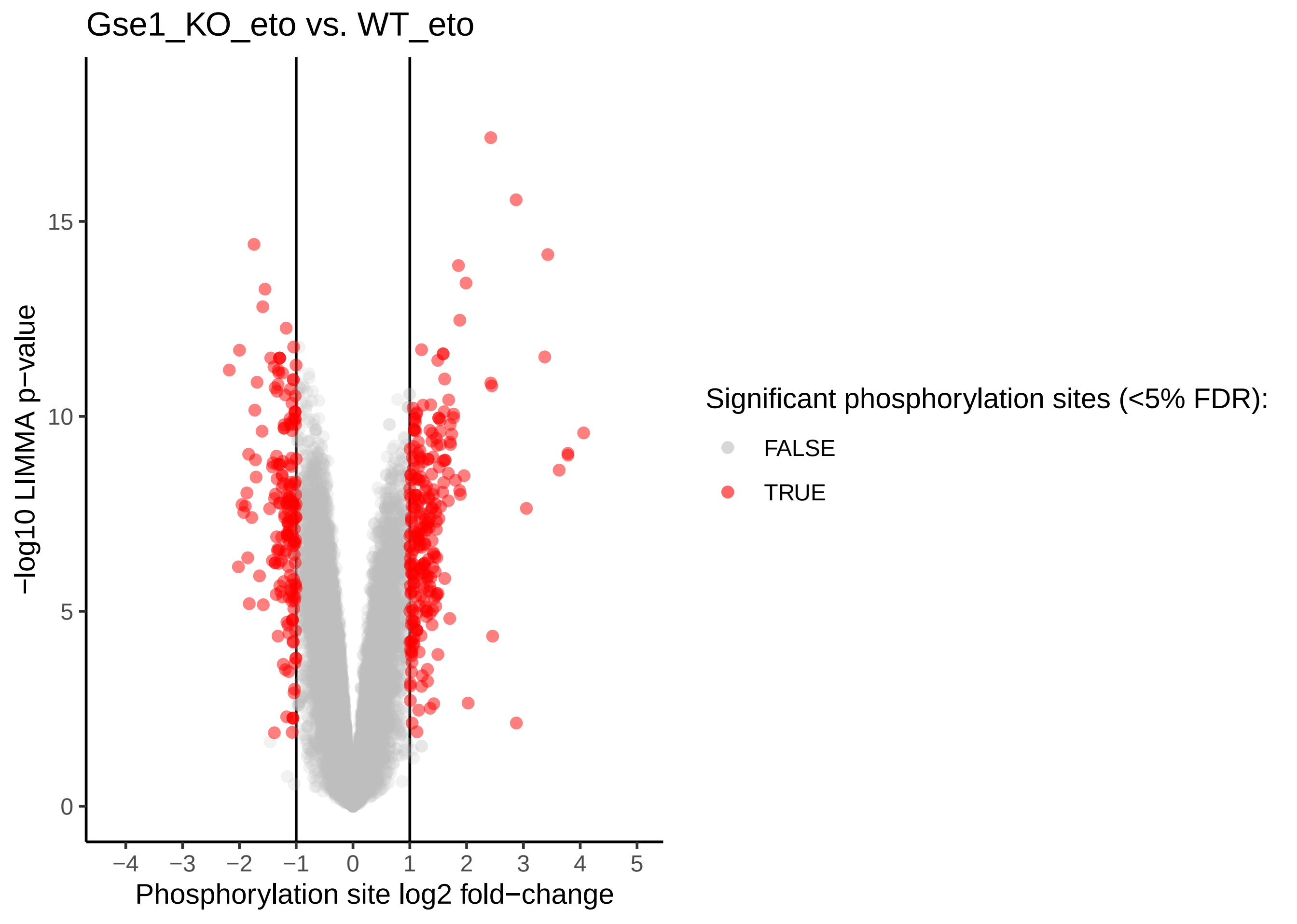

### ph_logFC.HDAC1.CI_ctrl-WT_ctrl.pdf

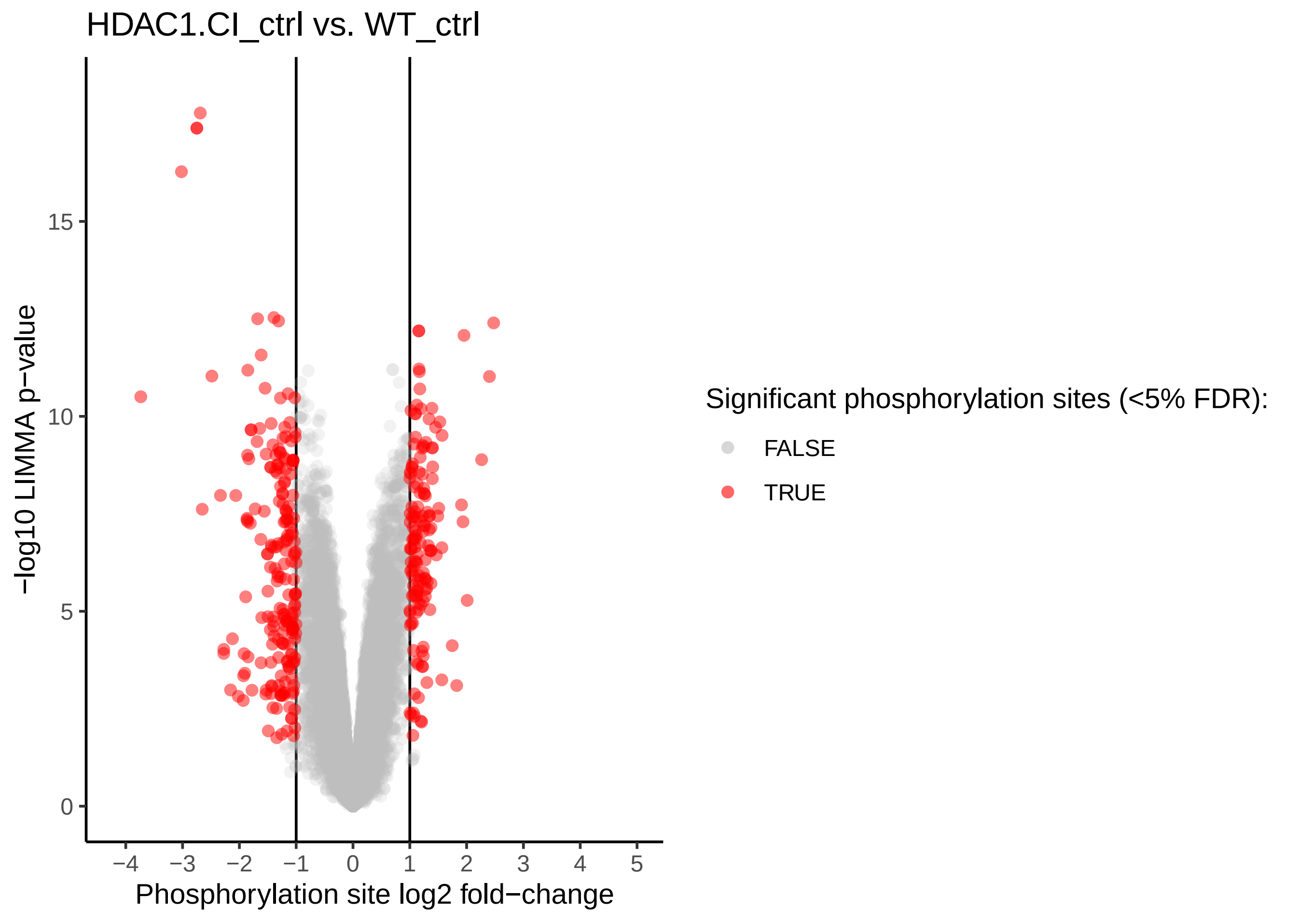

### ph_logFC.HDAC1.CI_eto-HDAC1.CI_ctrl.pdf

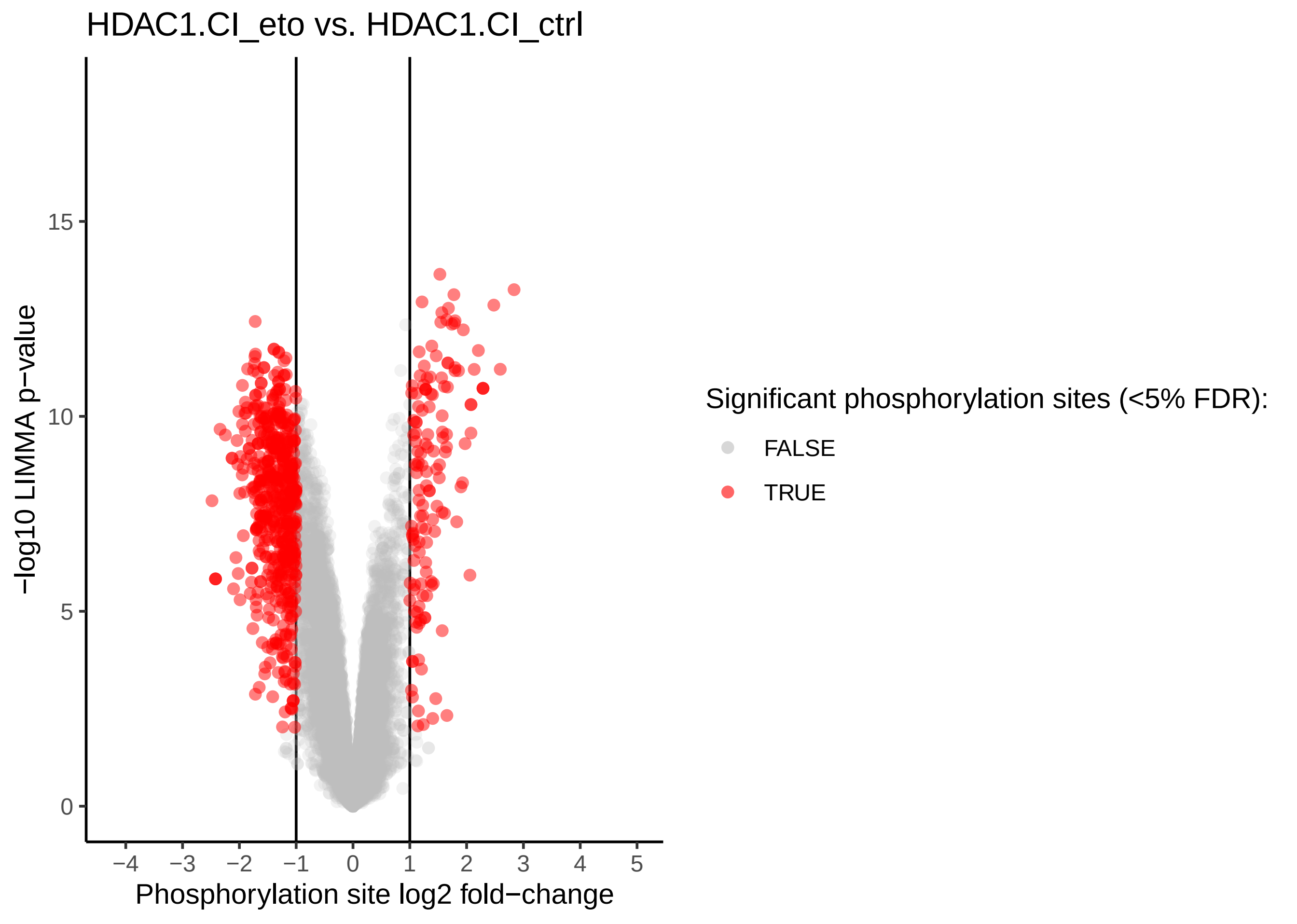

### ph_logFC.HDAC1.CI_eto-WT_eto.pdf

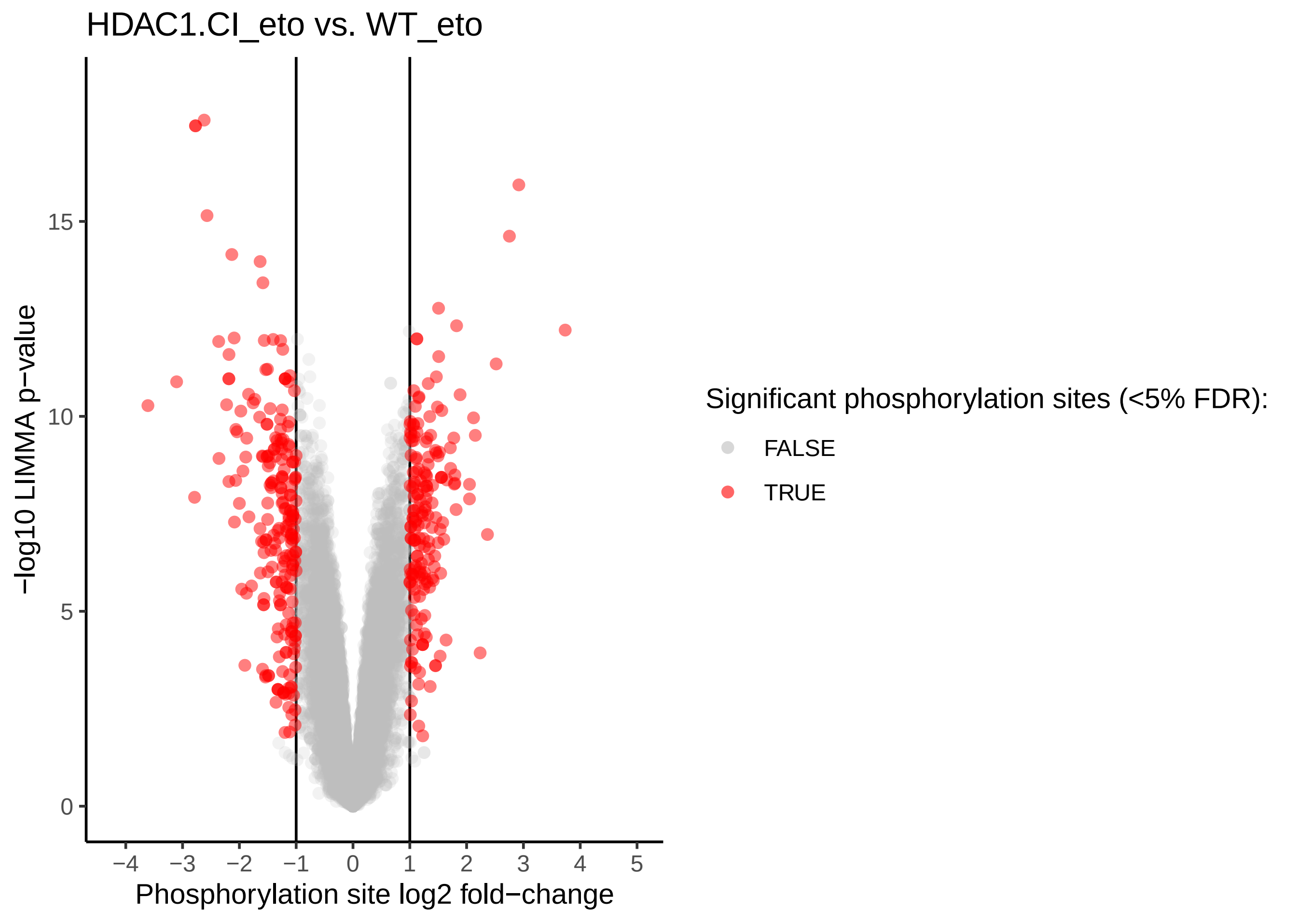

### ph_logFC.WT_eto-WT_ctrl.pdf

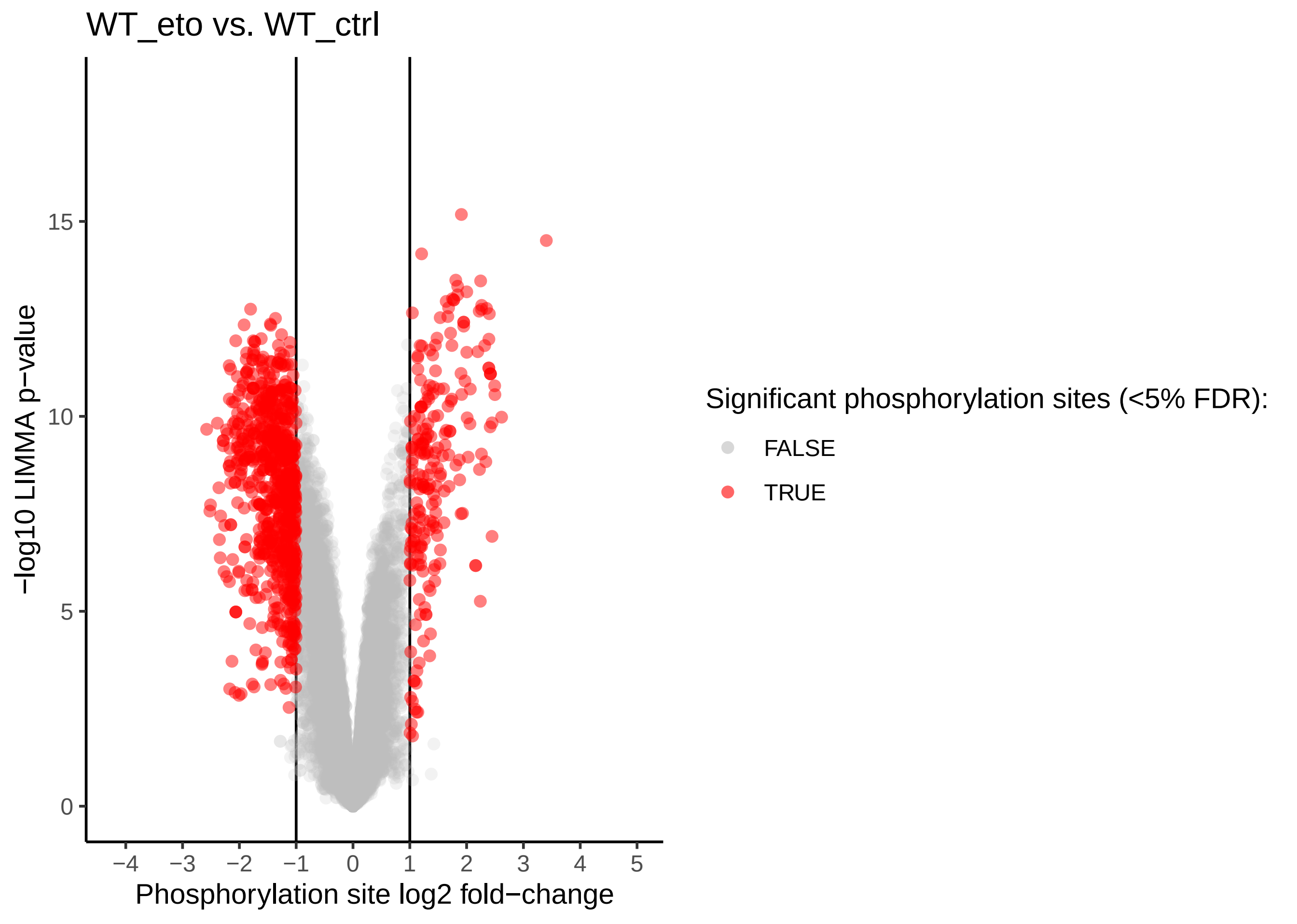
